# MicroRNA-541-3p alters lipoproteins to reduce atherosclerosis by degrading Znf101 and Casz1 transcription factors

**DOI:** 10.1101/2023.11.01.565110

**Authors:** Abulaish Ansari, Pradeep Kumar Yadav, Liye Zhou, Binu Prakash, Bhargavi Gangula, Laraib Ijaz, Amanda Christiano, Sameer Ahmad, Antoine Rimbert, M. Mahmood Hussain

## Abstract

High apoB-containing low-density lipoproteins (LDL) and low apoA1-containing high-density lipoproteins (HDL) are associated with atherosclerosis. In search of a molecular regulator that could simultaneously and reciprocally control both LDL and HDL levels, we screened a microRNA (miR) library using human hepatoma Huh-7 cells. We identified miR-541-3p that both decreases apoB and increases apoA1 expression by inducing mRNA degradation of two different transcription factors, Znf101 and Casz1. Znf101 enhances apoB expression while Casz1 represses apoA1 expression. The hepatic knockdown of orthologous *Zfp961* and *Casz1* genes in mice altered plasma lipoproteins and reduced atherosclerosis without causing hepatic lipid accumulation, most likely by lowering hepatic triglyceride production, increasing HDL cholesterol efflux capacity, and reducing lipogenesis. Notably, human genetic variants in the *MIR541*, *ZNF101*, and *CASZ1* loci are significantly associated with plasma lipids and lipoprotein levels. This study identifies miR-541-3p and Znf101/Casz1 as potential therapeutic agent and targets, respectively, to reduce plasma lipoproteins and atherosclerosis without causing liver steatosis.

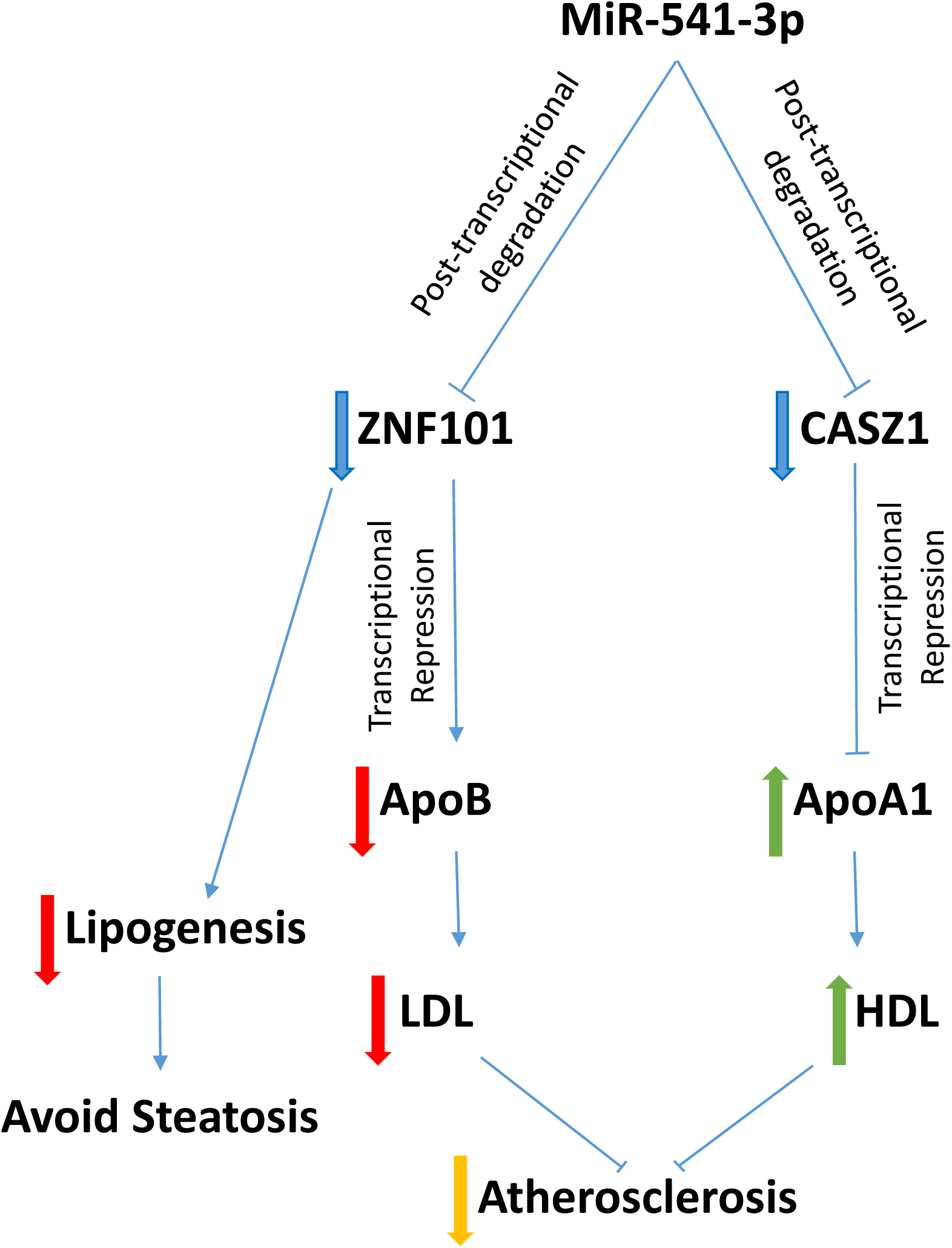
Graphical Abstract. A schematic diagram depicting the role of miR-541-3p in the control of plasma lipoproteins and atherosclerosis. Our data show that miR-541-3p downregulates *ZNF101* and *CASZ1* by enhancing post-transcriptional degradation of mRNAs after interacting with their 3′-UTRs. Furthermore, our data indicate that *ZNF101* is an enhancer of *APOB*, and *CASZ1* is a repressor of *APOA1*. Hepatic knockdown (KD) of *Zfp961*, an orthologue of *ZNF101*, reduces plasma apoB-containing lipoproteins, whereas KD of *Casz1* increases high density lipoproteins in mice. Furthermore, we show that hepatic KDs of these transcription factors reduces atherosclerosis in mice induced by the expression of mutant PCSK9.

## Introduction

MicroRNAs (miRs), small (∼22 nucleotides) non-coding RNAs, bind via their seed sequences to the 3’-untranslated regions (3’-UTRs) of target mRNAs to either degrade them or to decrease translation^1,2^. MiRs regulate many genes; therefore, miR mimics, or their inhibitors, anti-miRs, can be used to understand regulation of different interrelated biological pathways. Such knowledge will be useful in developing drugs to target multiple pathways for favorable clinical outcomes. Several miR-based drugs are in clinical trials^3–5^.

High plasma cholesterol levels are risk factors for atherosclerotic cardiovascular disease (ASCVD)^6,7^. Cholesterol is carried in the blood by apoB-containing low-density lipoproteins (apoB-Lps) and apoA1-containing high-density lipoproteins (HDL). ApoB-Lps are synthesized and secreted by the liver and the intestine to deliver endogenous and dietary lipids to peripheral tissues. Excess accumulation of modified apoB-Lps in the plasma, and their uptake by macrophages, contributes to ASCVD^8,9^. HDL extracts and transports free cholesterol from macrophages to the liver for excretion from the body^10,11^. Hence, low HDL is also associated with ASCVD. Therefore, understanding physiological and molecular mechanisms controlling plasma low-density lipoproteins (LDL) and HDL levels will aid in developing new therapeutics against ASCVD. To this end, LDL-lowering therapies are widely used to reduce heart disease. Yet, a significant portion of the population does not respond and/or is intolerant to these drugs^12,13^. Therefore, drugs have been developed to lower LDL by inhibiting hepatic production of apoB-Lps^14^. However, these approaches are associated with hepatosteatosis^15^. Furthermore, increasing plasma HDL therapeutically has not been effective in lowering ASCVD^11,16^. Hence, there is a need to develop new LDL lowering therapies and to increase functional HDL levels. Notably, agents that concurrently decrease LDL and increase HDL have not been identified.

## Results

### MiR-541-3p reciprocally regulates apoB and apoAI production

We transfected 1237 human miR mimics into human hepatoma Huh-7 cells and quantified secreted apoB and apoA1 (**Extended Data** Fig 1A**, Extended Data Table 1**). The screening performed in duplicate plates showed high correlation (**Extended Data** Fig 1B), and good internal reproducibility as different members of the same miR families that share the same seed sequence similarly reduced apoB and apoA1 secretion (**Extended Data** Fig 1C). Additionally, miR-30c-5p and miR-33a-5p reduced apoB and apoA1, respectively, (**Extended Table 1**) consistent with earlier studies^17,18^. We then selected 24 miR mimics that showed differential effects on apoB and apoA1 secretion for additional screening. Of these, 6 miR mimics increased apoA1 secretion (**Extended Data** Fig 1D) whereas 13 miR mimics decreased apoB secretion (**Extended Data** Fig 1E). In this study, we selected miR-541-3p for detail studies because it decreased apoB secretion by 58±9%, and increased apoAI secretion by 95±3%. Increasing concentrations of miR-541-3p and antimiR-541-3p significantly decreased and increased, respectively, media and cellular apoB in dose-dependent manner, as measured by ELISA and Western blotting (**Fig 1A, Extended Data** Fig 2A-C). In contrast, miR-541-3p significantly increased secreted and cellular apoA1 in a dose-dependent manner, whereas antimiR-541-3p decreased apoA1 protein (**Fig 1B, Extended Data** Fig 2D-F). Similar reciprocal modulation of apoB and apoA1 secretion was observed in human primary hepatocytes (**Fig 1C**). MiR-541-3p had no effect on *MTTP* or *ABCA1* mRNA levels, proteins critical for apoB-Lps and HDL biosynthesis (**Extended Data** Fig 2G). These studies suggest that miR-541-3p simultaneously and reciprocally regulates apoB and apoA1 production.

**Figure 1.**
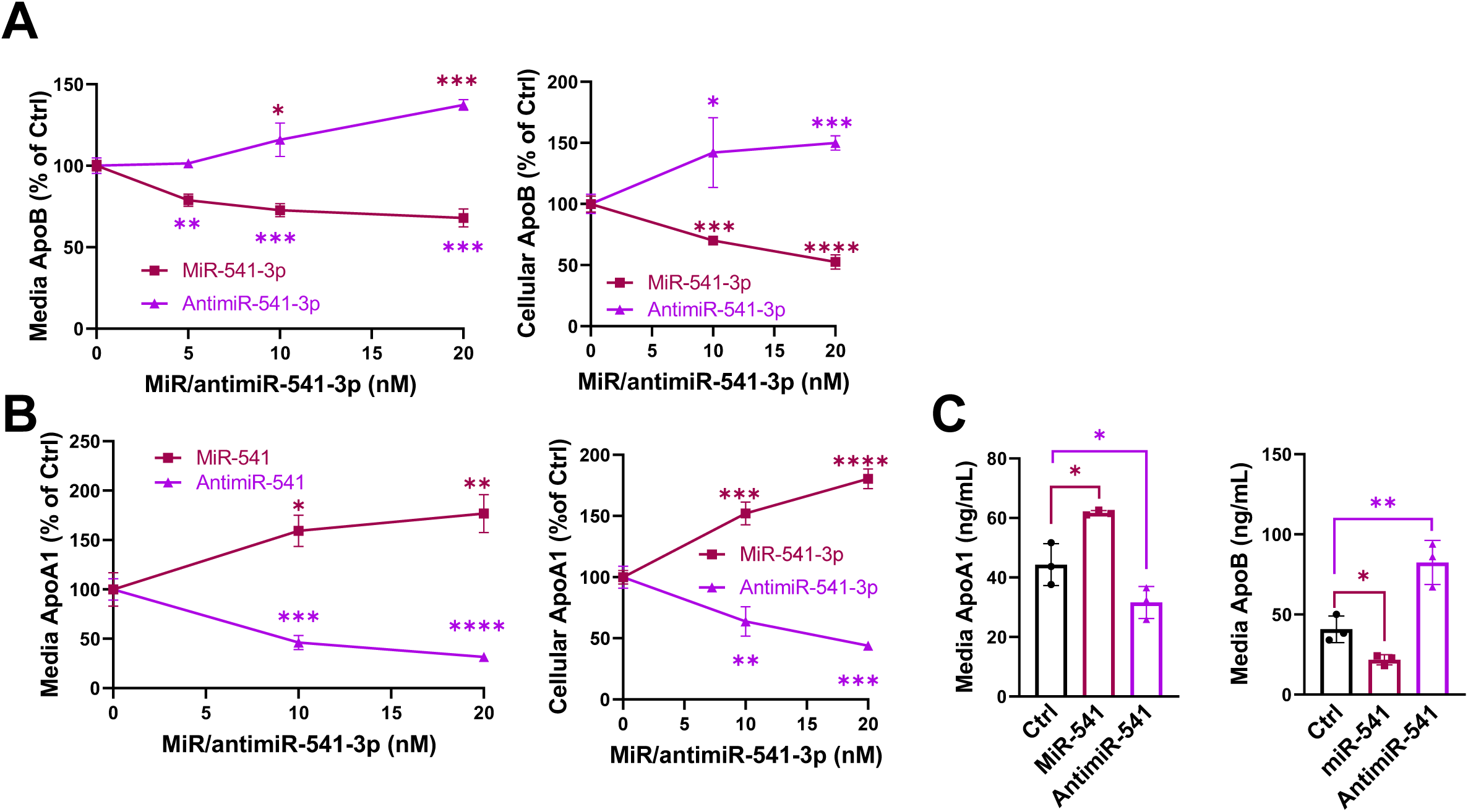
MiR-541-3p reciprocally regulates apoB and apoA1 secretion in human liver cells. (**A-B**) Huh-7 cells in 6-well plates were reverse transfected in triplicate with different amounts of miR-541-3p mimics or antimiR-541-3p. Control cells were transfected with a control miR (20 nM) and were used as 100% control. After 48 h, media was changed and collected after overnight incubation. ApoB (A) and apoA1 (B) levels were quantified in media and in cell lysates in triplicate by ELISA and normalized to cellular protein levels. Data are representative of 3 experiments. *, P < 0.05; **, P < 0.01; ***, P < 0.001; ****, P < 0.0001, two-way ANOVA, non-parametric. (**C**) Human primary hepatocytes were seeded in 24-well plates. After one day, they were transfected in triplcate with miR-541-3p mimics or antimiR-541-3p (10 nM, n = 3). After 16 h, media was replaced with fresh media. After 24 h, media was used to quantify apoB and apoA1 by ELISA in triplicates. *, P < 0.05; **, P < 0.01; unpaired t-test.

### MiR-541-3p mimics decrease apoB production by reducing Znf101 expression

To understand mechanisms how miR-541-3p regulates apoB expression, we transfected Huh-7 cells with miR-541-3p mimic and antimiR-541-3p and quantified mRNA levels. MiR-541-3p mimics decreased *APOB* mRNA, while anti-miR-541-3p had the opposite effect (**Fig 2A**). Since miR-541-3p is not predicted to pair with the 3’-UTR of *APOB* mRNA or *APOB* promoter sequences, we hypothesized that miR-541-3p might indirectly modulate mRNA by targeting transcription factors (TFs) regulating *APOB* expression. Using online tools, we identified seven TFs that could regulate apoB expression, are targets of miR-541-3p, and are expressed in Huh-7 cells (**Extended Data** Fig 3A-B). Notably, miR-541-3p mimics significantly decreased the expression of *ZNF101* mRNA, while antimiR-541-3p increased it (**Fig 2B, Extended Data** Fig 3C). Reducing Znf101 with siZnf101 significantly decreased apoB transcript levels and protein secretion without affecting apoA1 (**Fig 2C**). The expression of miR-541-3p mimics, independently or combined with siZnf101, revealed that both miR-541-3p and Znf101 regulate apoB through the same pathway (**Fig 2D, Extended Data** Fig 3D). These studies show that miR-541-3p regulates Znf101 to decrease apoB expression.

**Fig 2.**
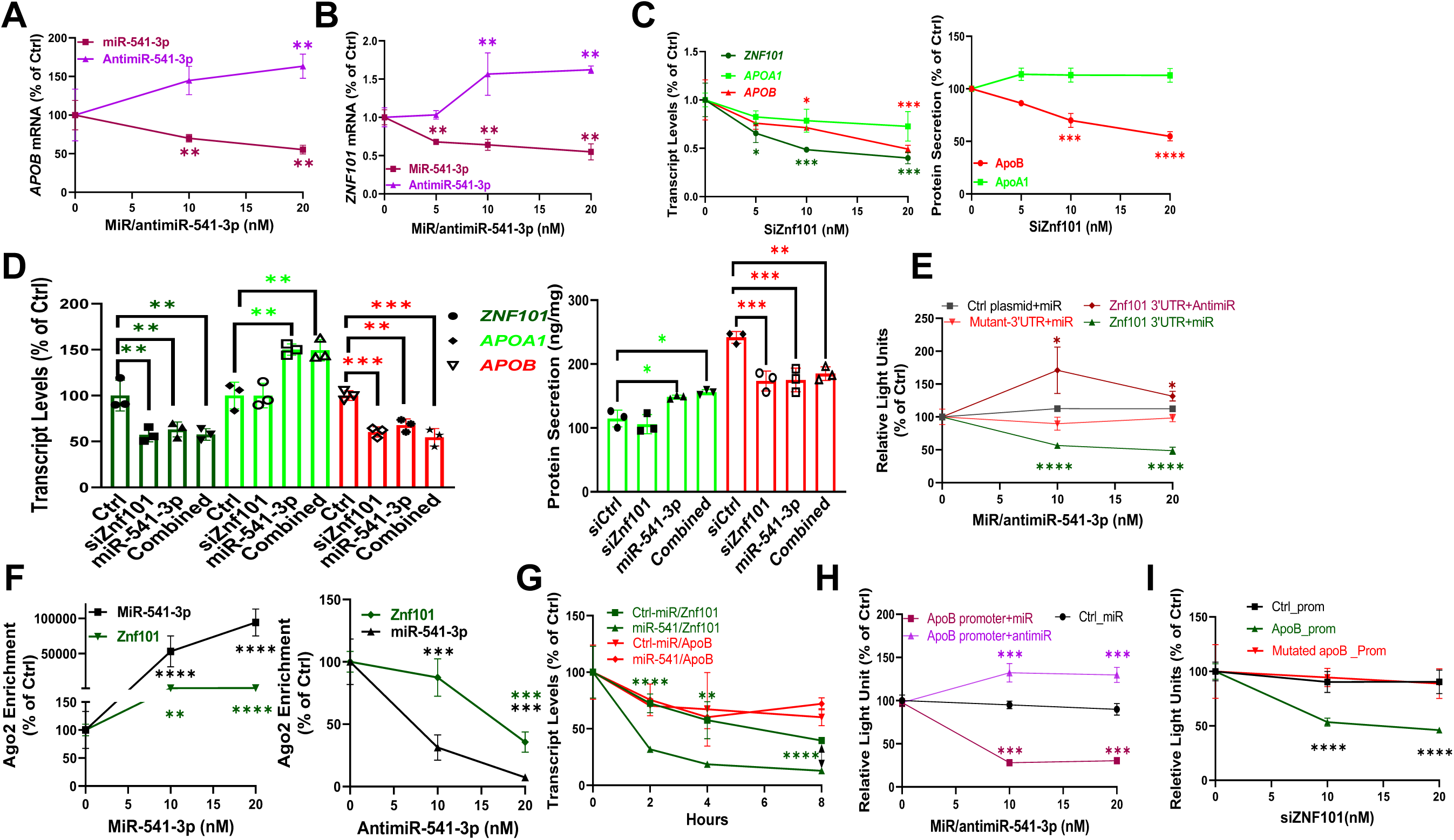
Regulation of apoB by miR-541-3p. **(A-B)** Huh-7 cells were transfected in triplicate with different amounts of miR-541-3p mimics or antimiR-541-3p, and *APOB* and *ZNF101* mRNA levels were quantified in triplicate. **(C)** Cells were transfected in triplicate with different concentrations of specific siRNA against *ZNF101* (siZnf101). After 48 h, mRNA levels of *ZNF101*, *APOA1* and *APOB* were quantified (left). Media was used to quantify apoB and apoA1 protein levels (right). **(D)** Huh-7 cells were transfected in triplicate with siZnf101 (21 nM) or miR-541-3p mimics (10 nM), alone or in combination. Changes in *ZNF101*, *APOA1*, and *APOB* mRNA were quantified in triplicate in cell lysates (left), and protein from conditioned media (right). **(E)** Cells were forward transfected (n=3) with plasmids (5 µg) expressing the dual reporter Gaussia luciferase/secreted alkaline phosphatase under control of the Znf101 3′-UTR or a control plasmid without Znf101 3′-UTR. Next day, these cells were reverse transfected in triplicate with miR-541-3p mimics or antimiR-541-3p. After 48 h, media was measured for luciferase and alkaline phosphatase activities in triplicates. **(F)** Cells transfected (n=3) with miR-541-3p mimics (left) or antimiR-541-3p (right) were used for Ago2 immunoprecipitation and measurements of miR-541-3p and Znf101 mRNA. **(G)** Cells were transfected in triplicate with 20 nM miR-541-3p mimics or a control miR in triplicate. After 18 h, cells were washed and treated with actinomycin D (10 µg/mL). At indicated times, Znf101 and apoB mRNA levels were quantified in triplicates. **(H)** Cells were transfected in triplicate with plasmids expressing luciferase under the control of the apoB or cytomegalovirus promoter. Next day, cells were equally distributed in wells and transfected in triplicate with different amounts of miR-541-3p mimics or antimiR-541-3p. After 48 h, luciferase activity was measured in triplicates in conditioned media. **(I)** Cells were transfected in triplicate with plasmids expressing luciferase under control of the wild type or mutated apoB promoter. Next day, cells were equally distributed and transfected with different amounts of siZnf101 in triplicates. After 48 h, conditioned media was used to measure luciferase activity in triplicate. *P < 0.05; **, P < 0.01, ***P < 0.001; ****P < 0.0001. One-way ANOVA non-parametric test.

To test that miR-541-3p interacts with the 3’-UTR of *ZNF101* mRNA and induces its degradation, we used four approaches. First, we identified that the *ZNF101* mRNA 3’-UTR contains a miR-541-3p interacting site conserved in primates (**Extended Data** Fig 3E). Second, we mutated the 3’-UTR of *ZNF101* and found that miR-541-3p mimics were unable to decrease *ZNF101* transcription (**Fig 2E, Extended Data** Fig 3F). Third, expression of miR-541-3p mimics enriched miR-541-3p and *ZNF101* mRNA, while expression of antimiR-541-3p depleted their levels in Ago2 complexes (**Fig 2F**). Fourth, we treated cells with actinomycin D to inhibit gene transcription and studied post-transcriptional mRNA degradations in miR-541-3p mimic treated cells. Post-transcriptional degradation of *ZNF101* mRNA was faster in cells transfected with miR-541-3p mimics compared to controls, while *APOB* mRNA decay was similar in cells transfected with either miR-541-3p mimics or the control miR mimics indicating temporal delay in the regulation of *APOB* mRNA by Znf101 (**Fig 2G**). Thus, miR-541-3p likely interacts with the 3′-UTR of *ZNF101* mRNA in Ago2 complexes to induce post-transcriptional degradation.

A potential Znf101 binding site in the *APOB* promoter exists at +418 bp after the transcription start site (**Extended Data** Fig 3G). Luciferase activity under control of the *APOB*-promoter decreased after the expression of miR-541-3p mimics, but increased with antimiR-541-3p expression (**Fig 2H**). Furthermore, siZnf101 reduced *APOB*-promoter luciferase activity (**Fig 2I**), but not when the Znf101 binding site was mutated, indicating that Znf101 is an enhancer of *APOB*. Thus, miR-541-3p downregulates Znf101, thereby reducing apoB production.

### MiR-541-3p mimics increase apoA1 production by reducing Casz1 expression

To understand regulation of apoA1 by miR-541-3p, we quantified apoA1 transcript levels in Huh-7 cell lysates expressing miR-541-3p mimics and antimiR-541-3p. Transfection of increasing concentrations of miR-541-3p mimics enhanced, while antimiR-541-3p decreased, *APOA1* mRNA levels, respectively (**Fig 3A**). *APOA1* mRNA is not a target of miR-541-3p and BLAST revealed no complementarity between miR-541-3p and *APOA1*. Thus, Watson-Crick base pairing between miR-541-3p and *APOA1* mRNA is unlikely. Subsequently, we identified TFs that might regulate *APOA1*, contain miR-541-3p target sites in their 3’-UTRs, and are expressed in Huh-7 cells (**Extended Data** Fig 4A-B). Our studies found that transcript levels of *CASZ1* were significantly decreased following overexpression of miR-541-3p mimics, while antimiR-541-3p had the opposite effect on *CASZ1* mRNA (**Fig 3B, Extended Data** Fig 4C). Transfection of siCasz1 significantly reduced *CASZ1* and increased *APOA1* mRNA expression and protein secretion, without affecting apoB transcript or protein levels (**Fig 3C**). Thus, miR-541-3p reduces Casz1 repressor to enhance apoA1 expression.

**Fig 3.**
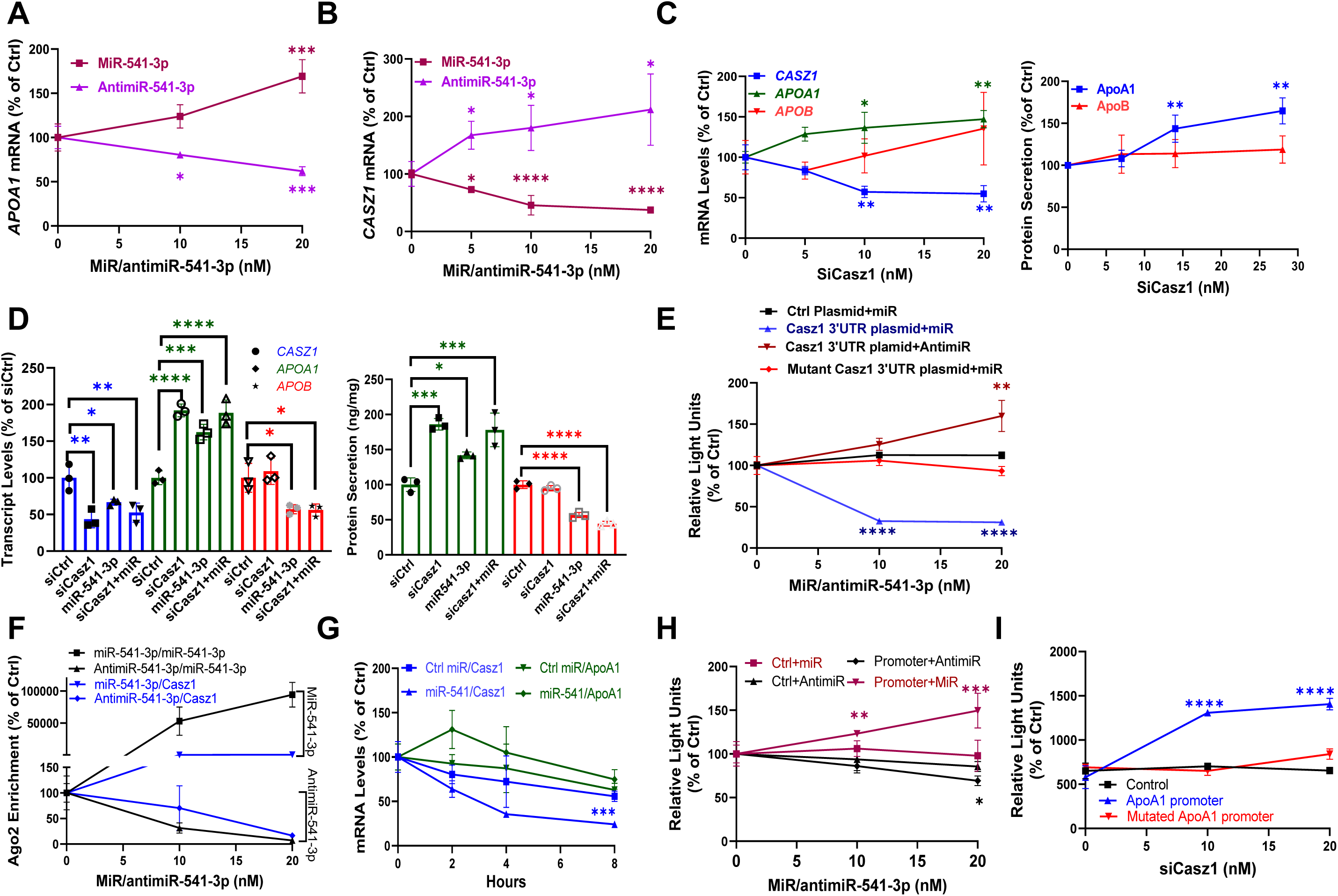
Regulation of apoA1 by miR-541-3p. **(A-B)** Huh-7 cells were forward transfected in triplicate with miR-541-3p mimics or antimiR-541-3p. After 48 h, mRNA levels of *APOA1* and *CASZ1* were quantified in triplicate. **(C)** SiCasz1 transfected cells (n=3) were collected to measure mRNA (left) and media to quantify apoB and apoA1 protein levels (right) in triplicates. **(D)** Cells were transfected in triplicate with 10 nM of siCtrl or siCasz1 or 21 nM miR-541-3p mimimcs, individually or in combination. Cells were used to measure different mRNAs (left) and media to measure apoB and apoA1 protein (right) in triplicate. **(E)** Plasmids for the expression of luciferase with or without the 3′-UTR of Casz1 were transfected (n=3). After 24 h, cells were reversed transfected in triplicate with different amounts of miR-541-3p mimics or antimiR-541-3p. After 24 h, luciferase activity was measured in triplicate in media. **(F)** Cells were transfected (n=3) with increasing amounts of miR-541-3p mimics or antimiR-541-3p. After 48 h, Ago2 immunoprecipitates were used to quantify miR-541-3p and *CASZ1* mRNA levels in triplicates and expressed as a percent of control miR transfected cells. **(G)** Transfected cells were treated with actinomycin D (10 µg/mL) in triplicate and collected at different intervals to measure *CASZ1* and *APOA1* mRNA levels in triplicates. **(H)** Cells were transfected (n=3) with apoA1 promoter luciferase constructs. Next day, they were reverse transfected (n=3) with increasing concentrations of miR-541-3p mimics or antimiR-541-3p. After 48 h, luciferase activity was measured in triplicates. **(I)** Cells were transfected in triplicate with plasmids expressing luciferase under the control of the wild type or mutated apoA1 promoter. Next day, cells were equally distributed and transfected (n=3) with different amounts of siCasz1. After 48 h, luciferase activity was measured in triplicate in conditioned media. *P < 0.05; **, P < 0.01, ***P < 0.001; ****P < 0.0001. One-way ANOVA non-parametric test.

MiR-541-3p mimics and siCasz1, either individually or combined, decreased *CASZ1* and increased *APOA1* mRNA and protein secretion, to similar extents (**Fig 3D, Extended Data** Fig 4D), indicating they are in the same pathway. TargetScan predicted that *CASZ1* mRNA contains a miR-541-3p interacting site in its 3’-UTR that is conserved in primates (**Extended Data** Fig 4E). MiR-541-3p mimics decreased luciferase activity when expressed with the 3’-UTR of *CASZ1* mRNA, whereas antimiR-541-3p increased luciferase activity (**Fig 3E, Extended Data** Fig 4F). These effects were abolished after mutagenesis of the 3’-UTR of *CASZ1* mRNA (**Fig 3E**). Higher miR-541-3p and *CASZ1* mRNA were in Ago2 precipitates from miR-541-3p mimics transfected cells, whereas cells transfected with antimiR-541-3p had lower miR-541-3p and *CASZ1* mRNA (**Fig 3F**). *CASZ1*, but not *APOA1*, mRNA was degraded faster in miR-541-3p mimic expressing cells compared to control cells (**Fig 3G**). These studies indicate that miR-541-3p likely interacts with the 3′-UTR and induces post-transcriptional degradation of *CASZ1* mRNA.

A potential Casz1 binding site is located at –789 bp of the transcription start site in the apoA1 promoter (**Extended Data** Fig 4G). MiR-541-3p mimics increased, while antimiR-541-3p decreased, luciferase activity under the control of the *APOA1* promoter (**Fig 3H**). Furthermore, siCasz1 significantly increased *APOA1* promoter activity, and this increased activity was abolished following mutagenesis of the Casz1 binding site in the *APOA1*-promoter (**Fig 3I**). Thus, miR-541-3p modulates the Casz1 repressor to regulate apoA1 expression.

### Mouse orthologs of human *MIR541*, *ZNF101* and *CASZ1* genes

To evaluate *in vivo* roles of *MIR541*, *ZNF101* and *CASZ1*, we searched for their mouse orthologs. Comparison of human and mouse miR-541-3p sequences revealed three differences each in seed and non-seed sequences (**Extended Data** Fig 5A). Furthermore, predicted mRNA targets for mouse and human miR-541-3p are different. Thus, hsa-miR-541-3p and mmu-miR-541-3p have different seed sequences and are not functional orthologs.

Mouse *Casz1* is an ortholog of human *CASZ1* (GeneCards). Knockdown (KD) of Casz1 in mouse hepatocyte AML12 cells significantly increased *Apoa1* expression (**Extended Data** Fig 5B). Mouse has two orthologs of human *ZNF101*, *Zfp101* and *Zfp961*. The KD of *Zfp101* had no effect on *Apob* mRNA. However, siZfp961 significantly reduced *Zfp961* and *Apob* mRNA levels (**Extended Data** Fig 5B). Thus, mouse *Casz1* and *Zfp961* genes are functional orthologs of human *CASZ1* and *ZNF101* with respect to *Apoa1* and *Apob* gene regulation.

Since high fat diets affect plasma lipids and atherosclerosis, we next asked whether hepatic Casz1 and Zfp961 are regulated *in vivo* by these diets. C57Bl6 mice were fed chow, Western, or obesogenic diets. High fat diets increased *Zfp961* mRNA levels but had no effect on *Casz1* expression (**Extended Data** Fig 5C).

### Hepatic knockdown of Casz1 and Zfp961 alter plasma apoB-Lps and HDL

To test whether Casz1 or Zfp961 regulate hepatic apoB and apoA1 levels, we transduced adult C57BL/6J mice with adenoviruses expressing shRNAs against both TFs, either alone or together. Both shCasz1 and shZfp961 specifically reduced hepatic *Casz1* and *Zfp961* mRNA, respectively, establishing their specificities (**Fig 4A-B**). Hepatic *Apoa1* mRNA levels increased (∼ 2.3-fold) in shCasz1 transduced mice, but remained undisturbed in shZfp961 transduced mice (**Fig 4C**). *Apob* transcript levels did not change in shCasz1 transduced mice, but were significantly reduced (∼ 60%) in shZfp961 transduced mice (**Fig 4D**). Thus, reduction of hepatic *Casz1* increases *Apoa1*, whereas *Zfp961* KD decreases *Apob* expression.

**Fig 4.**
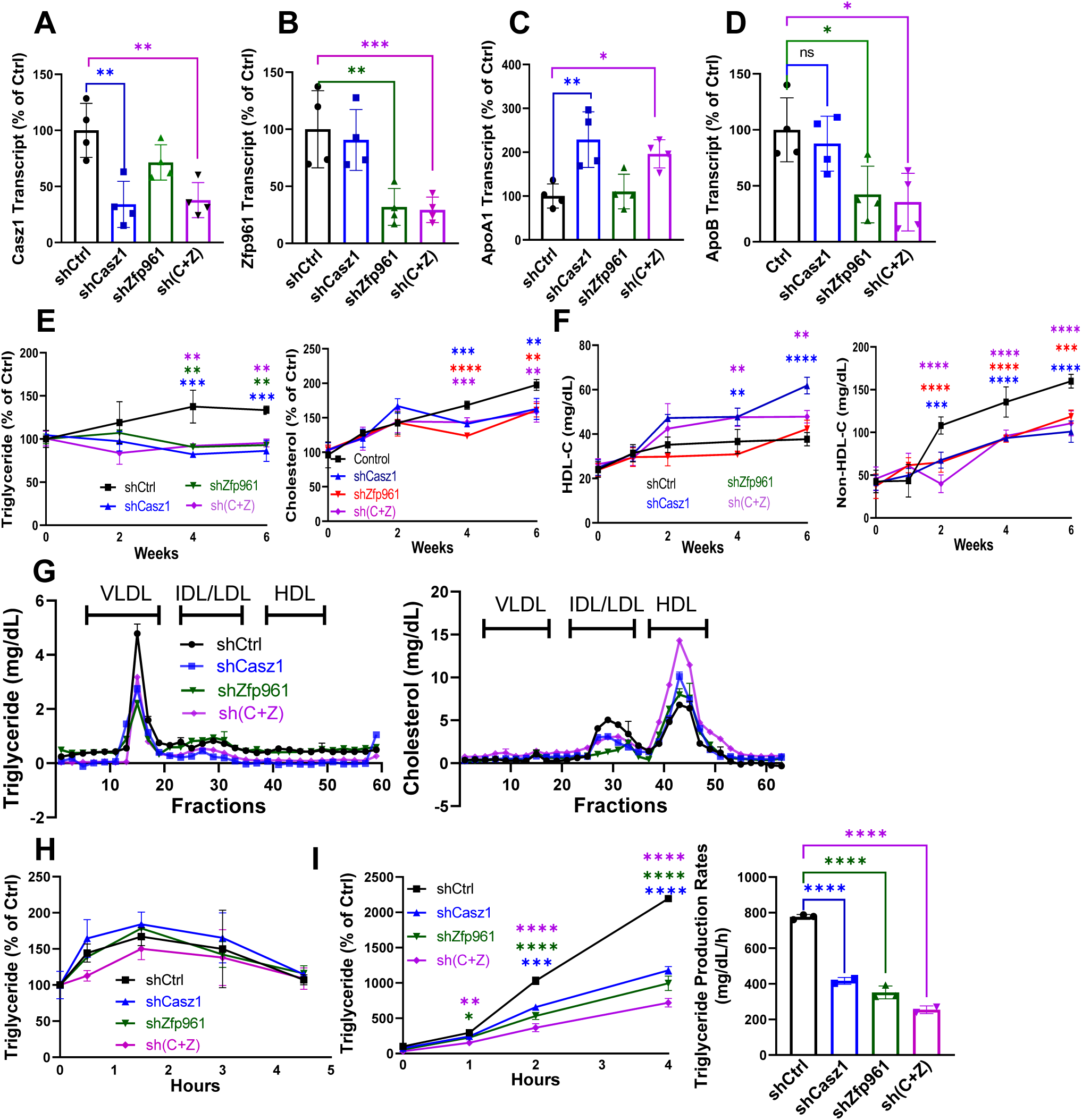
Hepatic knockdowns of Casz1 and Zfp961 alter plasma lipoproteins. Mice (C57Bl6J, female, 2.5-month-old) were transduced with AAV8 (2.5 x 10^11^ gc/mouse) expressing shControl (shCtrl, *n* = 4), shCasz1 (*n* = 4), shZfp961 (*n* = 4), or (sh(C+Z), *n* = 5), and fed a Western diet. **(A-D)** After 6 weeks, livers were collected to measure mRNA levels in triplicates. **(E-F)** Plasma was collected from fasting mice to measure triglyceride and cholesterol (E) in technical duplicates, and subjected to precipitation using polyethylene glycol to measure cholesterol in HDL and non-HDL fractions (F) in triplicate. **(G)** Plasma (200 µL) obtained at the end of the study was subjected to FPLC, and triglyceride and cholesterol were measured in different fractions. *P < 0.05; **, P < 0.01, ***P < 0.001; ****P < 0.0001. One-way ANOVA non-parametric test. **(H)** Mice were gavaged with olive oil (100 µL) after overnight fast. Plasma was collected at different times to measure triglycerides in duplicates. Increases in plasma triglyceride were plotted as % of initial values before gavage. **(I)** Mice were fasted overnight prior to blood collection. Mice were injected intraperitoneally with Poloxamer P407 (1.5 mg/g). Blood was collected to measure triglyceride (left) in triplicate. Data between 2 and 4 h were used to calculate production rates (right).

KD of Casz1 and Zfp961 had no effect on hepatic *Ppara*, *Pgc1a*, *Cebp*, *Mttp*, *Abca1*, and *Ldlr* mRNA, hepatic triglyceride/cholesterol, and plasma ALT/AST levels compared to control mice (**Extended Data** Fig 6A-C). Similarly, these KDs had no effect on total weight gain, or on other physical and physiological parameters, but they reduced fat mass (**Extended Data** Fig 6D-E, **Extended Data Table S2**).

The rise in plasma triglyceride and cholesterol over time was significantly lower in shCasz1 and shZfp961 transduced mice compared to in shCtrl mice (**Fig 4E**). HDL-cholesterol (HDL-C) significantly increased (∼ 63%) in shCasz1 and double shCasz1+shZfp961 transduced mice, but were unaffected by shZfp961 KD alone (**Fig 4F**). KD of either Zfp961 and Casz1, or both together, however, significantly reduced non-HDL-C (**Fig 4F**). Most of the triglycerides were in very low-density lipoprotein (VLDL) fractions in control mice, but the levels were reduced in all KD mice compared to controls (**Fig 4G**). FPLC studies further showed that cholesterol was mainly in HDL of control mice. HDL-C was unaffected by shZfp961 but increased in mice transduced with shCasz1 and shCasz1+shZfp961 (**Fig 4G**), indicating that increases in plasma cholesterol in shCasz1 mice are due to increases in HDL-C. These studies showed that hepatic KDs of Casz1 and Zfp961, either separately or together, reduce triglyceride and non-HDL-C, whereas only Casz1 KD increases HDL-C. Importantly, KD of these TFs had no effect on intestinal lipid absorption (**Fig 4H**); while hepatic triglyceride production rates were significantly reduced (**Fig 4I**). Our findings suggest that hepatic inhibition of both Casz1 and Zfp961 may reduce plasma lipids and apoB-Lps, as a result of impaired hepatic lipoprotein production.

### Atherosclerosis is reduced following hepatic knockdown of Casz1 and Zfp961

To test whether KD of Casz1 and/or Zfp961 could reduce atherosclerosis, we used a mouse atherosclerosis model, in which adult mice are transduced with mutant mouse gain-of-function Pcsk9 (mPcsk9) ^19^. mPcsk9 augments degradation of hepatic LDL receptors, increases plasma cholesterol levels and induces atherosclerosis. ShZfp961 and shCasz1 both specifically reduced their own target mRNAs; while shZfp961 reduced apoB mRNA levels without affecting apoA1, shCasz1 increased apoA1 without affecting apoB mRNA levels (**Extended Data** Fig 7A). Control mice transduced with mPcsk9+shCtrl showed increased plasma triglyceride, cholesterol and non-HDL-C after 5–10 weeks (**Fig 5A**). Mice transduced with shCasz1 and shZfp961, on the other hand, had significantly reduced plasma levels of triglyceride, cholesterol, and non-HDL-C, while the shCasz1 transduced mice also had higher HDL-C levels compared to shCtrl (**Fig 5A**). Triglyceride and cholesterol in VLDL and IDL/LDL fractions were significantly reduced in all KD mice compared to controls (**Fig 5B**). In contrast, KD of Casz1 significantly increased plasma HDL-C levels. Plasma apoA1 protein levels increased in shCasz1 transduced mice but they were not affected by shZfp961 (**Fig 5C, Extended Data** Fig 7B). ApoB100 and apoB48 levels were significantly reduced in all KD mice (**Fig 5C, Extended Data** Fig 7C). ShCasz1 and shZfp961 had no significant effect on other TFs, lipid genes, hepatic lipids and plasma transaminases (**Extended Data** Fig 8A-C). Thus, KDs of Casz1 and Znf961 reduce plasma cholesterol and triglyceride in apoB-Lps without affecting hepatic lipids, whereas Casz1 KD, in addition, increases HDL-C. Plasma and HDL from shCasz1 transduced mice showed increased cholesterol efflux from J774 macrophages compared to shCtrl (**Fig 5D**) indicating increased cholesterol efflux capacity.

**Fig 5.**
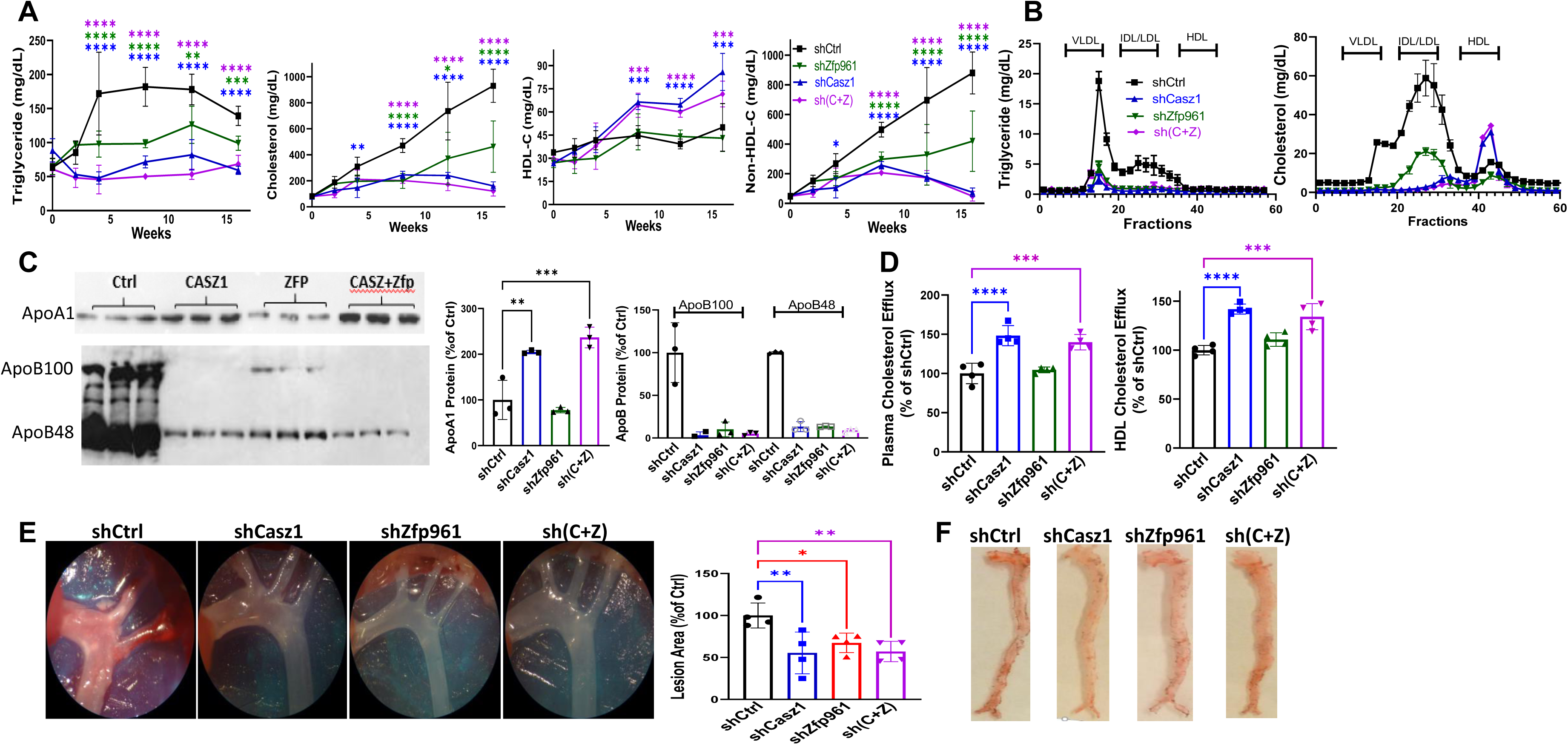
Hepatic knockdown of Casz1 and Zfp961 in mutant PCSK9 transduced mice reduces atherosclerosis. All mice (C57B16J, male, 2.5-month-old) received AAV8 (5 x 10^11^ gc) for the expression of mutant mouse gain-of-function Pcsk9 (mPcsk9). They were divided into four groups and received AAV8 for the expression of either shCtrl (5 x 10^11^, *n* = 4), shCasz1 (shCasz1 2.5 x 10^11^ + 2.5 x 10^11^ shCtrl, *n* = 4) shZnf101 (shZnf101 2.5 x 10^11^ + shCtrl, 2.5 x 10^11^, *n* = 4) or shCasz1+shZnf101 (shCasz1, 2.5 x 10^11^ + shZnf101, 2.5 x 10^11^, *n* = 5) at the same time. Mice were fed an atherogenic Western diet. **(A-B)** Plasma was collected from overnight fasted mice to measure triglyceride, cholesterol, HDL-C, and non-HDL-C (A) in duplicates, and lipoprotein characterization using FPLC (B). **(C)** Plasma (1 µL) was separated on gels and probed for apoA1 (top) and apoB (bottom) using specific antibodies. Images were quantified using ImageJ and plotted as % of Ctrl. **(D)** Total plasma (10 µL, left) and HDL (20 µg/mL, right) were used in triplicates to study efflux of ^3^H-cholesterol from J774 macrophages. **(E-F)** After 4 months, mice were dissected to visualize plaques in the aortae (E). Total aortas were opened and stained with Oil Red O to visualize lipid deposition (F). *, P < 0.05; **, P < 0.01; ***, P < 0.001; ****, P < 0.0001, one way ANOVA, non-parametric. Representative of two experiments.

ShCasz1 and shZfp961 transduced mPcsk9 mice gained less weight and had significantly less fat than control mPcsk9 mice (**Extended Data** Fig 8D, **Extended Data Table S3**). ShCasz1 increased energy expenditure and reduced food intake in the dark, but shZfp961 had no effect (**Extended Data** Fig 8E). Atherosclerotic plaques were significantly reduced by ∼40-50% in all TF KD mPcsk9 mice (**Fig 5E-F, Extended Data** Fig 9A-B).

The above studies showed that shCasz1 and shZfp961 significantly reduced plasma apoB but had no significant effect on hepatic lipids and plasma AST/ALT; markers that are usually increased after reductions in apoB-Lp assembly. To identify reasons for the absence of hepatosteatosis, we quantified mRNA levels and observed no changes in fatty acid oxidation gene expression (**not shown**); however, mRNA levels of the lipogenic proteins Srebp1c, Acc1, Scd1, and glucokinase were significantly reduced (**Extended Data** Fig 9A-B). Furthermore, expression of miR-541-3p reduced, while anti-miR-541-3p enhanced newly synthesized cellular and secreted triglyceride and phospholipids in Huh-7 cells (**Fig 6A-B**). In addition, KD of Znf101 significantly reduced synthesis and secretion of triglycerides and phospholipids (**Fig 6C-D**). SiCasz1 cells tended to reduce lipid synthesis. These studies suggest that reduced lipid synthesis might have prevented hepatosteatosis.

**Fig. 6.**
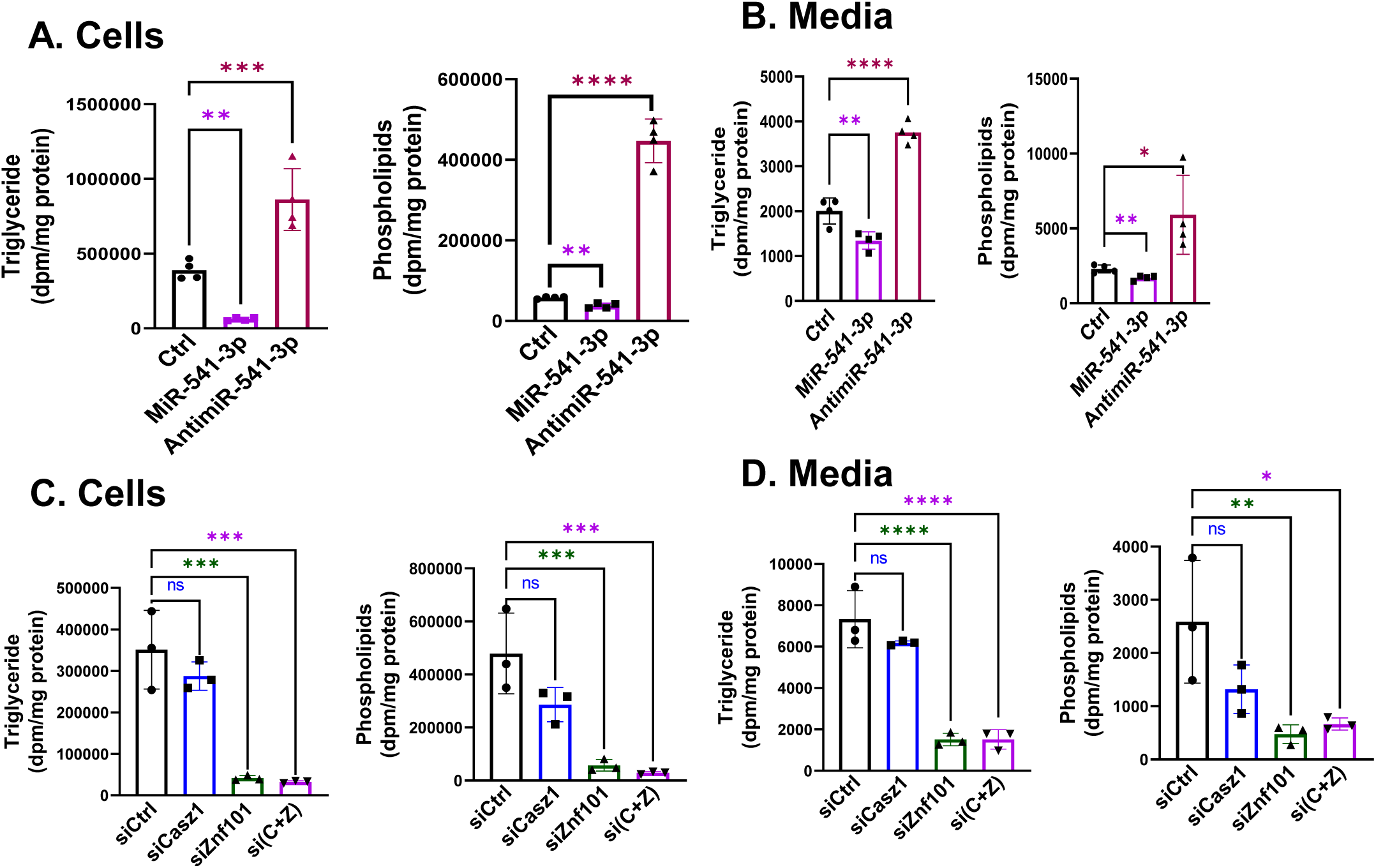
Triglyceride and phospholipid synthesis is reduced in Huh-7 cells overexpressing miR-541-3p and siZnf101. Huh-7 cells were transfected with (A-B) miR-541-p or antimir-541-3p (20 nM) or (C-D) different siRNAs (10 nM) in triplicates. After 24 h, cells were washed and incubated with ^3^H-glycerol (5 µCi) for 16 h. Lipids were extracted from media using CHCl_3_ and CH_3_OH. Cells were washed and incubated with isopropanol for 16 h at 4°C. Lipids were evaporated, resuspended in isopropanol, and separated on thin layer chromatography plates. Bands corresponding to triglyceride and phospholipid were excised and counted.

### Association of genetic variants in *MIR541*, *ZNF101* and *CASZ1* loci with plasma lipids

The above studies identified *MIR541*, *CASZ1* and *ZNF101* genes as modulators of lipid metabolism in human liver cells. To extend the relevance of these genes to human lipid metabolism, we screened for genetic associations of variants in the *MIR541, CASZ1 and ZNF101* loci with plasma lipids in the UK-Biobank. We found significant associations between: rs7161194, located ∼1.8 kb upstream of *MIR541*, and higher plasma HDL-C levels; rs34071855, located in the second intron of *CASZ1*, and reduced plasma cholesterol, LDL-C, and apoB levels; and rs2304130, located in the second intron of *ZNF101*, and reduced plasma cholesterol, LDL-C, apoB and triglyceride levels (**Fig 7, Extended Data Table S4**). These associations were further replicated in the Global Lipid Genetics Consortium (GLGC) dataset ^20^ (**Extended Fig 11A**). The observed significant associations suggest for possible role of these genes in determining plasma lipid and lipoprotein levels. In addition, we studied the associations of these SNPs in *MIR541*, *CASZ1* and *ZNF101* with expression of nearby genes in the human liver using expression quantitative trait loci (eQTL) from the GTEx dataset (V8) ^21^. <u>No significant eQTLs</u> were found for these genes in the liver. Despite no associations, we ran colocalization studies to match GWAS data from the GLGC together with liver eQTL dataset (from GTEx V7). We could not identify SNPs associated with both expression of CASZ1 and with total cholesterol plasma levels using LocusComparer R library^22^ (**Extended Data** Figure 11B) nor with *ZNF101* and *MIR541* (data not shown). These studies show that different SNPs in these loci have significant association with plasma lipids and lipoproteins; however, we could not find evidence showing that these SNPs impact the expression of *CASZ1*, *ZNF101* or *MIR541*. Thus, further studies are warranted to establish the potential link between genetic association signals and the actual causal genes and biological mechanisms.

**Fig. 7.**
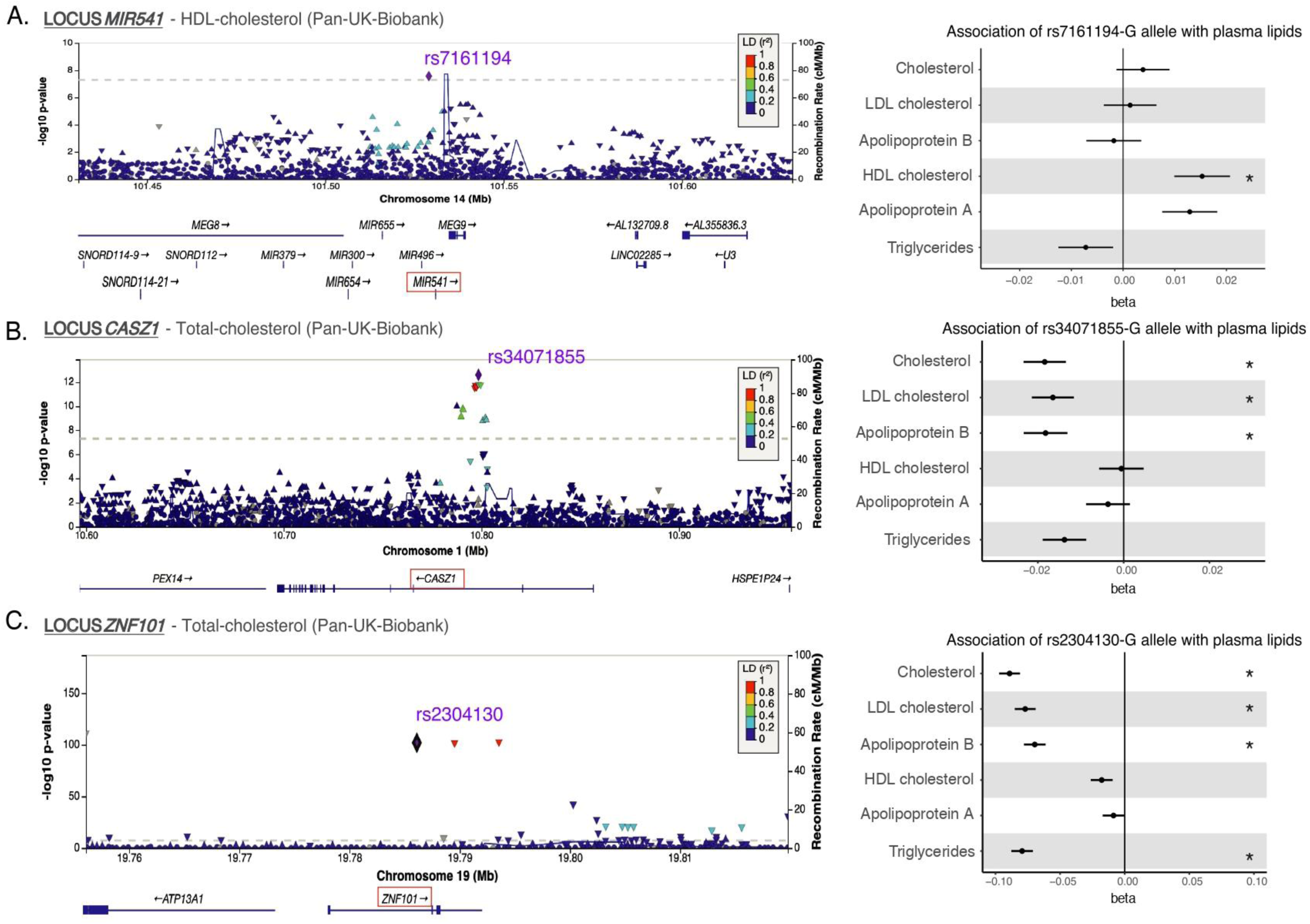
Regional plots showing genome wide associations of variants in the *MIR541*, *CASZ1,* and *ZNF101* loci with plasma lipids and lipoproteins. Association results for SNPs (-log10 P value, y-axis) as a function of genomic coordinates using human gene version 19 (hg19) for (A) *MIR541*, (B) *CASZ1,* and (C) *ZNF101* gene loci with plasma total cholesterol, LDL, apoB, HDL-C, apoA1, and triglycerides in the UK Biobank using the Pan-UKB project summary statistics. The bottom panel shows genes at each locus, as annotated in the UCSC Genome Browser. The most highly associated SNPs are represented as purple diamonds, and the linkage disequilibrium (LD) values (1000 Genomes data) are in the inset. Light blue dotted lines indicate estimated recombination hotspots. Forrest plot on the right panels show the associations (beta ± standard error) of the top associated SNP with plasma lipids and apolipoproteins levels. (*) Correspond to genome wide significant p-values (p<5.0E-08). Raw data are presented in Supplemental Table 3.

## Discussion

We provide evidence that miR-541-3p mimics decrease human apoB and increase apoA1 expression by degrading mRNA levels of two transcription factors, Znf101 and Casz1, in human liver cells. In mouse liver cells, Zfp961 enhances the expression of apoB and plasma LDL, whereas Casz1 reduces the expression of apoA1 and plasma HDL levels. KD of these TFs decreases plasma apoB-Lps, lipogenesis and atherosclerosis, without causing hepatosteatosis or increasing AST/ALT in mice. We further find that plasma lipid levels are associated with human genetic variants in the *MIR541*, *ZNF101* and *CASZ1* loci.

Post-translational mechanisms regulate apoB-Lps synthesis^23^. Here, we describe that Znf101 KD reduces apoB mRNA and protein levels in cultured cells and in mice, indicating that Znf101 regulates plasma apoB-Lps at the transcriptional level. In contrast to Znf101, KD of Casz1 has no significant effect on apoB mRNA levels, but significantly reduces hepatic triglyceride production and plasma apoB levels in mice. These studies suggest that Casz1 might regulate production of apoB-Lps in hepatocytes, involving as of yet unknown post-transcriptional mechanisms.

We show that miR-541-3p interacts with the 3′-UTR of Znf101 and Casz1 to increase mRNA degradation. We, furthermore, find that Znf101 interacts with the *APOB* promoter at +418 to enhance transcription. This site is within the DNAse 1 hypersensitive promoter region responsible for apoB expression in liver cells, but not in Hela cells^24^. Moreover, we show that Casz1 interacts with the *APOA1* promoter to repress its expression. Thus, we have explained molecular mechanisms involved in the regulation of apoB and secretion by miR-541-3p.

Our data provide evidence that Znf101/Zfp961 and Casz1 affect production of apoB and apoA1 in mice. We show that KD of Znf101 decreases apoB mRNA levels in mouse livers (**Fig 4D**). We observed that hepatic TG production was less in Zfp961 KD mice than control mice (**Fig 4I**). These data indicate that Zfp961 regulates apoB-containing lipoprotein production. Furthermore, we provided evidence in Huh-7 cells that KD of Znf101 reduces apoB mRNA levels and its secretion (**Fig 2C-D**). We have also provided evidence that Znf101 interacts with apoB promoter and that these interactions are abolished after site-directed mutagenesis (**Fig 2H**). Similarly, we have shown that KD of Casz1 increases mRNA levels and media apoA1 in Huh-7 cells (**Fig 3**). Furthermore, KD of Casz1 in mouse liver increased apoA1 mRNA levels (**Fig 4C**). Both, Casz1 and Zfp961/Znf101 are TFs. Hence, their effect is mainly anticipated to be at the level of transcription and protein production. In short, our data suggest that these TFs modulate apoB and apoA1 production by regulating their transcription. Besides these observations, we also observed that Casz1 reduced apoB levels in mice but not in cell culture systems. Therefore, it is possible that at organismal levels, Casz1 can modulate plasma lipoproteins by affecting additional pathways, such as intracellular apoB degradation, that require further investigation.

We attempted to correlate plasma miR-541-3p levels measured by Taqman with plasma triglyceride and cholesterol in healthy and obese subjects. The Ct values for miR-541-3p were 32 in both healthy and obese subjects. More sensitive measurements, such as droplet digital PCR, are needed to ascertain whether miR-541-3p can be used as a disease biomarker.

A limitation of the study is that we could not directly assess the physiologic role of miR-541-3p in mice as this is a primate specific miR. However, we did identify intermediary TFs through which miR-541-3p regulates apoB and apoA1 secretion in human liver cells. Furthermore, we identified orthologous TFs in mice with preserved functions in the regulation of apoB and apoA1 production. After identifying the intermediary TFs, we were able to demonstrate the preservation of plasma LDL and HDL regulatory mechanisms in both species. Thus, it is likely that miR-541-3p evolved in primates to regulate conserved regulatory mechanisms that control plasma lipoproteins levels.

Another caveat of our studies is that changes in Znf101 and Casz1 levels were documented via measuring mRNA levels. We have not been able to document changes in their protein levels as commercially procured have not yielded good bands.

In short, we identified a primate specific miR-541-3p that concurrently reduces apoB and increases apoA1 in human hepatoma cells. Furthermore, we identified that miR-541-3p regulates apoB and apoA1 by regulating two different TFs, Casz1 and Znf101. Next, we studied the *in vivo* roles of these TFs in the livers of mice on plasma lipids and atherosclerosis. Hepatic Casz1 KD increases apoA1, HDL, and augments cholesterol efflux capacity and could be a contributing factor for reduced atherosclerosis. KD of Zfp961 and Casz1 reduces plasma apoB-Lps, hepatic triglyceride production, and, most importantly, atherosclerosis. These studies suggest that miR-541-3p could be a therapeutic agent to simultaneously lower plasma apoB-Lps while increasing plasma HDL and reduce atherosclerosis. *ZNF101* and *CASZ1* are also potential therapeutic targets to modulate plasma LDL and HDL for beneficial outcomes.

## Acknowledgments

This work was supported in part by U.S. National Institutes of Health (NIH) grants DK121490, HL137202, HL158054, and HD094778, as well as VA Merit Award BX4113 and David R. Doucette Research Innovation Award to MMH. The content is the responsibility of the authors and does not represent the views of the NIH and VA.

Genetic studies were conducted using the UK Biobank resource under the application number 49823 and 31063. We thank the UK Biobank participants and the Pan-UK Biobank team as well as the Global Lipids Genetics Consortium for making the GWAS data publicly available. We are grateful to the Bioinformatics Core Facility of Nantes BiRD, member of Biogenouest, Institut Français de Bioinformatique (IFB) (ANR-11-INBS-0013) for the use of its resources and for its technical support in these analyses.

## Author contributions

MMH conceived the ideas, supervised experiments, and secured funding. LZ performed the initial screening of the microRNA library. BP and BG performed Oil Red O and Trichrome staining of the aortic arch cross sections as well as western blots for MTP and ABCA1. LE and AC performed several secondary screenings. PKY focused work on miR-541-3p. AA took over the project and performed all the cell culture and mechanistic studies and brought the studies to a fruitful conclusion. PKY and AA performed all the mice experiments. SA worked with AA in quantifying mRNA levels. AR performed genetic analyses. AA made the figures and wrote the first draft of the manuscript, which was extensively modified by MMH. All authors read the drafts and provided their suggestions on the manuscript.

## Methods

### Cell culture

Human hepatoma Huh-7, mouse hepatocyte AML12, and mouse macrophage J774 cells were cultured in Dulbecco’s modified Eagle’s medium (DMEM) containing 10% fetal bovine serum (FBS), and 1% L-glutamine in T75 flasks at 37°C in humidified 5% CO_2_ incubators^25,26^.

**Screening:** MiR mimics were suspended in RNase free water to obtain 2 μM stocks and 3 μL of each miR stock were added in duplicate wells to obtain a final concentration of 50 nM. To each well, we added 7 μL of Opti-MEM and 10 μL of lipofectamine RNAiMAX (Life technologies) diluted 1:20 in Serum Reduced Opti-MEM. After 20 to 30 minutes, 25,000 cells in 100 μL of Opti-MEM were added to each well. After additional 24 h, culture media were changed with fresh DMEM containing 10% fetal bovine serum. Media were changed 24 hours later and cells were incubated with DMEM containing oleic acid/BSA complex ((oleic acid (0.4mM)/BSA (1.5%)) for 2 h. ApoB and apoAI concentrations in medium were measured by ELISA (60). Cells were used for protein measurements.

### Transfection in hepatoma cells

Huh-7 or AML12 cells (2.5 x 10^5^) were reverse transfected in triplicate in 6 well plates using Endofectin (Genecopoeia) with the indicated doses of miRIDIAN miRNA mimics (miR-541-3p), miRIDIAN miRNA inhibitors (antimiR-541-3p), siRNAs (Origene), scramble control mimic (20 nM, Thermo Fisher Scientific), or siControl as noted in the figure legends. Media was replaced with fresh DMEM containing 10% FBS and 1% L-glutamine after 16 h of transfection. For long-term cultures, media was changed every 24 h.

### ApoA1 and apoB protein quantifications in Huh-7 cells by ELISA

One day after transfections, media was replaced with 1 mL of fresh DMEM containing 10% FBS. Cells and media (last 24 h) were collected 70 h post transfection. Cells were lysed with 0.1 N NaOH to quantify total protein by BCA method. Total cell protein was used to normalize amounts of the apoA1 and apoB protein in media. To measure apoA1 and apoB protein levels in Huh-7 cells, transfected cells were lysed with 1 x RIPA buffer containing protease inhibitor mixture (Sigma-Aldrich). Cell lysates were centrifuged and the supernatants were added to ELISA plates (Costar, Corning) to measure apoA1 (R&D Systems® ELISA Kits) and apoB (Mabtech ELISA Kits) in triplicate. In some samples, sandwich ELISA using 1D1 as capture antibodies was used for apoB ELISA in triplicate as previously described^25^.

### Quantification of mRNAs by quantitative RT-PCR

Huh-7 cells were washed with PBS, 1 mL of TRIzol was added, mixed vigorously for 30 s, and incubated for 10 min at room temperature. Similarly, 10 mg of mouse liver tissue was lysed in 1 mL of TRIzol and homogenized using a tissue homogenizer. Chloroform (200 µL) was added and mixed vigorously, left at room temperature for 5 min, centrifuged (12,000 rpm, 15 min, 4 °C), and the top aqueous phase was transferred to a new tube. Subsequently, samples were mixed with equal volumes of ice chilled isopropanol and kept at room temperature for 5 min. Samples were centrifuged (12,000 rpm, 15 min, 4°C) and supernatants were discarded, pellets were re-suspended in 1 mL of 70% ethanol, centrifuged (12,000 rpm, 15 min, 4 °C), and supernatants were discarded. Pellets were dried in 37 °C incubator for 10 min and dissolved in molecular biology grade water.

First strand cDNA was synthesized using 1 µg of RNA with the Applied Biosystems™ High-Capacity cDNA Reverse Transcription Kit was used for quantitative RT-PCR (PowerTrack SYBR Green Master Mix) in triplicate and the Ct values for each mRNA were normalized to 18S. For miR quantification, 100 ng of RNA and miR specific primers were used for cDNA synthesis using the TaqMan MicroRNA Reverse Transcription kit (Applied Biosystems, 4366597). MiR-541-3p levels were quantified using the Ct method after normalization with RNU44 or U6 and are presented as arbitrary units.

### Immunoblotting

Huh-7 cells (from 1 well of a 6 well plate) were mixed with 200 µL of cold 1 x RIPA buffer and vortexed. After complete cell lysis, lysates were centrifuged (1000 rpm for 10 min at 4°C). Total protein (25 µg) was resolved on 6% and 12% acrylamide gels for apoB and apoA1, respectively. Liver tissue (50 mg) was homogenized in 1 mL of buffer K (1 mM Tris-Cl, 1 mM EGTA, and 1 mM MgCl_2_, pH 7.6) containing protease inhibitor mixture. Proteins (∼25 μg) were resolved on SDS-PAGE gels, transferred to nitrocellulose blotting membranes by semi-dry transfer method and membranes were blocked with 5% dry milk powder in Tris-buffered saline (TBS) for 1 h. After washing with TBS containing 0.1% Tween® 20 detergent (TBST), membranes were incubated with 1:1000 dilution of primary polyclonal antibodies against mouse/human apoA1 and apoB or a rabbit antibody to human/mouse β-actin overnight at 4°C. The next day, membranes were washed and incubated with alkaline phosphatase conjugated secondary antibody (1:10,000 dilution) for 2 h, washed, substrate was added and bands were visualized using Storm 860 Molecular Imager.

### Luciferase assay

Plasmids expressing *Renilla* luciferase with Znf101-3’-UTR, Casz1-3’-UTR, apoA1-promoter, or apoB-promoter (GLuc/SEAP dual-reporter vector system) were obtained from Genecopiea. Huh-7 cells were first forward transfected with 5 µg of these plasmids. After 16 h of transfection, cells were trypsinized and equally distributed into wells and reverse transfected in triplicate with miR-541-3p, antimiR-541-3p, or different indicated siRNAs. Luciferase activity was measured in media after 8 h of reverse transfection using a kit. Luciferase activity was normalized with SEAP (secreted alkaline phosphatase) activity.

### Ago2 precipitation

Huh-7 cells transfected (n=3) with miR-541-3p or antimiR-541-3p were collected in buffer K and sonicated to lyse the cells. Cell lysates were centrifuged (10000 rpm, 10 min, 4°C), supernatants were collected and incubated for 1 h with IgG for preclearing. After centrifugation, supernatants were mixed with protein A/G plus agarose beads for 1 h at 4 °C. This mixture was centrifuged (800 rpm, 1 min) and supernatants were incubated with 1:100 dilution of human Ago2 antibody overnight at 4°C. Next day, mixtures were incubated with agarose beads for 1 h, and centrifuged (800 rpm, 1 min). Supernatants were discarded and pellets were processed for RNA isolation using TRIzol method.

### Actinomycin D study

After 16 h of miR-541-3p (20 nM) transfection, Huh-7 cells were treated with 10 μg ml^−1^ of actinomycin D in triplicate to analyze post-transcriptional degradation of different mRNAs. Cells were collected every 2 h and RNA was isolated using TRIzol. Isolated RNAs were used to quantify expression levels of apoA1, apoB, Casz1, and Znf101 transcript levels.

### De novo lipogenesis

Huh-7 cells (*n* = 4) were reverse transfected with control miR, miR/antimiR-541-3p (20 nM) in 12-well plates. After 8 h of transfection, cells were washed three times with PBS and incubated with 500 µL of serum free media containing ^3^[H]-glycerol (5 µCi/mL) for 16 h. Media was collected and processed for lipid isolation using the chloroform:methanol method^27^. Cells were washed three times with PBS and incubated overnight with 1 mL of isopropanol at 4°C. Isopropanol was collected and dried by vacuum centrifugation. Dried lipids were dissolved in 25 µL of chloroform and applied to thin layer chromatography (TLC) plates (Whatmann, aluminum plates coated with silica, 4420221) and separated using petroleum ether:diethyl ether:acetic acid (70:30:1) solvent mixture. TLC plates were exposed to iodine vapors for 2-5 min, triglyceride and phospholipid (PL) bands were marked with pencil, excised, placed in scintillation vial and 5 mL of ScintiSafe™ Econo 2 Cocktail was added. Each sample was counted in liquid scintillation counter (Wallac 1409) for 5 min to measure disintegrations per minute (dpm).

### Knockdown of TFs in mice

Female C57Bl/6 mice (*n* = 17, 2.5-month-old) were divided into 4 groups and were intravenously transduced with different viruses expressing specific shRNA (2.5 x 10^11^ genome copies) as follows: shControl (4 mice), shCasz1 (4 mice), shZnf101 (4 mice), and shCasz1+shZnf101 (5 mice). These mice were fed a Western diet (Envigo TD.88137 contains 15.2% protein, 42.7% carbohydrate, and 42% fat kcal as well as cholesterol 1.5 g/kg) for 2 months. In a separate experiment, male mice were transduced with these shRNAs along with adeno-associated virus expressing a gain-of-function mutant of mouse PCSK9 (AAV8-D377Y-mPCSK9, 1.0 x 10^11^ genome copies) and fed a Western diet for 4 months. Plasma was collected at different times to document changes in plasma cholesterol and triglyceride levels. At the end, mice were fasted overnight, euthanized, and tissues were collected. In these mice, we also evaluated atherosclerosis, by dissecting and examining aortas, as previously described^28^.

### Plasma and tissue lipid measurements

Mice were fasted overnight (15 h) and blood was collected in EDTA-rinsed tubes from retro-orbital venous plexus. Blood was centrifuged (7,000 rpm, 10 min), plasma was collected and stored at –80 °C. Total plasma cholesterol and triglyceride concentrations were enzymatically measured in triplicates using kits. HDL-C was measured after precipitating apoB-lipoproteins using equal volumes of HDL cholesterol reagent (Pointe Scientific). LDL-C values were determined by subtracting the concentrations of HDL-C from total lipids. For hepatic lipid measurements, liver pieces (∼50 mg) were homogenized in 1 mL of buffer K, and 200 µL was subjected to lipid extraction^27,29^. Plasma ALT (Cayman) and AST (Sigma) were measured in triplicates using kits according to the manufacturer’s instructions.

### Fast performance liquid chromatography (FPLC)

Plasma samples (200 μL) were loaded onto AKTA pure FPLC column [Superose™ 6 Increase 10/300 GL FPLC column (GE Healthcare). Lipoproteins were eluted with PBS and 0.25 mL fractions were collected at a flow rate of 0.4 mL/min.

### Cholesterol efflux

Mouse macrophage J774 cells were seeded in 12-well plates at a density of 75 x 10^3^ cells/well and cultured in humidified incubator (37 °C, 5% CO2). After 15 h, 250 µL of DMEM complete media containing 0.25 μCi [^3^H]-cholesterol + 5 μg acetylated LDL (Ac-LDL) were added in each well and incubated. After 48 h, cells were examined under the microscope to ensure healthy appearance and about 80% confluency, media was removed, and cells were washed three times with PBS. Next, 500 μL of Opti-MEM media containing 2 μM LXR agonist T0-901317 was added into each well. After 18 h, media was removed and cells were gently washed with PBS. Next, Opti-MEM (500 µL) containing acceptor (HDL-C, 5 μg) or (plasma, 10 μL) in triplicate were added into each well. One set of wells received Opti-MEM only for background. After 4 h, different media was collected and centrifuged at 13,000 rpm for 10 min. Plates were kept in the freezer for 1 h, prior to adding 500 μL dH_2_O to each well, and incubated on a shaker at 4 °C for 1 h to detach cells. Cells were pipetted up and down to break up cell clumps and obtain homogeneous suspensions. Media (100 μL) and cells (100 μL) were transferred to 7 mL scintillation vials, 5 mL of Insta-gel Plus (PerkinElmer) was added, mixed well, and [^3^H] dpm was measured in a scintillation counter. The following formula was used to measure cholesterol efflux from cells to the acceptor: % cholesterol efflux = (media count – background)/(media count – background) + (cell counts – background) x 100.

### Hepatic triglyceride production

Mice were fasted for 16 h and then 1.5 mg/gm (in 500 µL PBS) of poloxamer 407 (P407) was injected intraperitoneally. Plasma was collected at the indicated time points. Triglycerides were measured using a kit. Rates of triglyceride production were measured between 2 and 4 h.

### Intestinal lipid absorption studies

Mice were fasted overnight, then 100 µL of olive oil was administered by intragastric gavage. Blood samples were collected from retro-orbital venous plexus at different times. Plasma triglyceride levels were measured using kits.

### Aortic plaque analyses

Aortic arches were dissected and exposed for photography and quantified with ImageJ. Aortic fatty streaks were visualized with Oil Red O staining.

### Statistics

All data are presented as mean ± SD. Statistical significance (P < 0.05) was determined using Student’s *t*-test, one-way ANOVA, or two-way ANOVA (GraphPad Prism), and significant differences are represented as * P < 0.05; **P < 0.01; ***P < 0.001; ****P < 0.0001.

### Study Population

The UK Biobank study, described in detail previously^30^, is a population-based prospective cohort in the United Kingdom in which > 500,000 individuals aged between 40 and 69 years were included from 2006 to 2010. The study has been approved by the North West Multi-Centre Research Ethics Committee for the United Kingdom, the National Information Governance Board for Health and Social Care for England and Wales, and the Community Health Index Advisory Group for Scotland (https://www.ukbiobank.ac.uk/media/0xsbmfmw/egf.pdf). All participants gave informed consent (https://www.ukbiobank.ac.uk/media/gnkeyh2q/study-rationale.pdf).

### Blood biomarker measurement

Blood biomarker measurement details are described in the UK-biobank showcase (https://biobank.ndph.ox.ac.uk/showcase/label.cgi?id=17518). In brief, plasma cholesterol and triglyceride levels were measured using enzymatic assays (https://biobank.ndph.ox.ac.uk/showcase/showcase/docs/serum_biochemistry.pdf). HDL-C was measured using an enzyme immuno-inhibition method while LDL-C was measured using an enzymatic selective protection method. Apolipoprotein A1 (apoA1), apolipoprotein B (apoB) were measured using immuno-turbidimetric assays (Beckman Coulter (UK), Ltd). Alkaline phosphatase (ALP), alanine aminotransferase (ALT), aspartate aminotransferase (AST), and gamma-glutamyltransferase (GGT) were measured using enzymatic rate methods. All biomarkers were measured on a Beckman Coulter AU5800 (Beckman Coulter (UK), Ltd).

### Genetic association of common variants with biomarkers in the UK Biobank

The genetic associations with biomarkers were extracted from the Pan-ancestry genetic analysis of the UK Biobank, by the Pan-UK Biobank team [released June 15, 2020]. In short, genetic and phenotypic data from ∼500,000 participants in the UK Biobank (https://www.ukbiob4ank.ac.uk) were used to conduct a Genome-Wide Association Study (GWAS). Genotypes were imputed from the Haplotype Reference Consortium plus UK10K & 1000 Genomes reference panels as released by UK-Biobank in March 2018. Plotting of associations within genetic loci has been performed using LocusZoom^31^. This research was conducted using the UK Biobank Resource (project ID 31063), and the use of these data is bound by all terms of usage of the UK Biobank. P-values below 5.0E–8 were considered as genome wide significant.

### Genetic analysis, data processing and plotting

The combination of genetic and phenotypic data, as well as data plotting was performed using RStudio (v.2022.02.1) with R-packages ggplot2 and ggforest.

## Methods

### Quantification of mRNAs by quantitative RT-PCR

Huh-7 cells were washed with PBS, 1 mL of TRIzol was added, mixed vigorously for 30 s, and incubated for 10 min at room temperature. Similarly, 10 mg of mouse liver tissue was lysed in 1 mL of TRIzol and homogenized using a tissue homogenizer. Chloroform (200 µL) was added and mixed vigorously, left at room temperature for 5 min, centrifuged (12,000 rpm, 15 min, 4 °C), and the top aqueous phase was transferred to a new tube. Subsequently, samples were mixed with equal volumes of ice chilled isopropanol and kept at room temperature for 5 min. Samples were centrifuged (12,000 rpm, 15 min, 4 °C) and supernatants were discarded, pellets were re-suspended in 1 mL of 70% ethanol, centrifuged (12,000 rpm, 15 min, 4 °C), and supernatants were discarded. Pellets were dried in 37 °C incubator for 10 min and dissolved in molecular biology grade water.

### Knockdown of TFs in mice

Female mice (*n* = 17, 2.5-month-old) were divided into 4 groups and were intravenously transduced with different viruses expressing specific shRNA (2.5 x 10^11^ genome copies) as follows: shControl (4 mice), shCasz1 (4 mice), shZnf101 (4 mice), and shCasz1+shZnf101 (5 mice). These mice were fed a Western diet (Envigo TD.88137 contains 15.2% protein, 42.7% carbohydrate, and 42% fat kcal as well as cholesterol 1.5 g/kg) for 2 months. In a separate experiment, male mice were transduced with these shRNAs along with adeno-associated virus expressing a gain-of-function mutant of mouse PCSK9 (AAV8-D377Y-mPCSK9, 1.0 x 10^11^ genome copies) and fed a Western diet for 4 months. Plasma was collected at different times to document changes in plasma cholesterol and triglyceride levels. At the end, mice were fasted overnight, euthanized, and tissues were collected. In these mice, we also evaluated atherosclerosis, by dissecting and examining aortas, as previously described^28^.

### Cholesterol efflux

Mouse macrophage J774 cells were seeded in 12-well plates at a density of 75 x 10^3^ cells/well and cultured in humidified incubator (37 °C, 5% CO_2_). After 15 h, 250 µL of DMEM complete media containing 0.25 μCi [^3^H]-cholesterol + 5 μg acetylated LDL (Ac-LDL) were added in each well and incubated. After 48 h, cells were examined under the microscope to ensure healthy appearance and about 80% confluency, media was removed, and cells were washed three times with PBS. Next, 500 μL of Opti-MEM media containing 2 μM LXR agonist T0-901317 was added into each well. After 18 h, media was removed and cells were gently washed with PBS. Next, Opti-MEM (500 µL) containing acceptor (HDL-C, 5 μg) or (plasma, 10 μL) in triplicate were added into each well. One set of wells received Opti-MEM only for background. After 4 h, different media was collected and centrifuged at 13,000 rpm for 10 min. Plates were kept in the freezer for 1 h, prior to adding 500 μL dH_2_O to each well, and incubated on a shaker at 4 °C for 1 h to detach cells. Cells were pipetted up and down to break up cell clumps and obtain homogeneous suspensions. Media (100 μL) and cells (100 μL) were transferred to 7 mL scintillation vials, 5 mL of Insta-gel Plus (PerkinElmer) was added, mixed well, and [^3^H] dpm was measured in a scintillation counter. The following formula was used to measure cholesterol efflux from cells to the acceptor: % cholesterol efflux = (media count – background)/(media count – background) + (cell counts – background) x 100.

### Genetic association of common variants with plasma lipoproteins from the Global Lipids Genetics Consortium

The Global Lipids Genetics Consortium aggregated GWAS results from 1,654,960 individuals from 201 primary studies representing five genetic ancestry groups^20^. Summary statistics from the meta-analysis results were downloaded via the following link http://csg.sph.umich.edu/willer/public/glgc-lipids2021.

### Association between genetic variants and expression of nearby genes (cis-eQTL)

Expression quantitative trait loci (eQTL) analyses aim to connect genetic variants with the expression of genes, or specifically of nearby genes, in case of cis-eQTL. We studied the effects of top associated genes on the expression of nearby genes in the human liver using the eQTL dataset from GTEx (V7)^21^.

### Genetic analysis, data processing and plotting

The combination of genetic, phenotypic and eQTL data, as well as data plotting was performed using RStudio (v.2022.02.1) with R-packages ggplot2, ggforest and locus comparer^22^.

## Abbreviations

ApoA1: apolipoprotein A1
ApoB: apolipoprotein B
ApoB-Lps: apoB-containing lipoproteins
GWAS: genome wide association studies
HDL: high density lipoproteins
KD: knockdown
LDL: low density lipoproteins
miRs: microRNAs
TF: transcription factor
UTR: untranslated region

## MATERIALS

| Diets | % Fat | Company, ID |
| --- | --- | --- |
| Chow | 13% | Lab Diet, 5053 |
| Western | 45% | Envigo, TD.88137 |
| Obesogenic | 60% | Research Diets, D12492 |

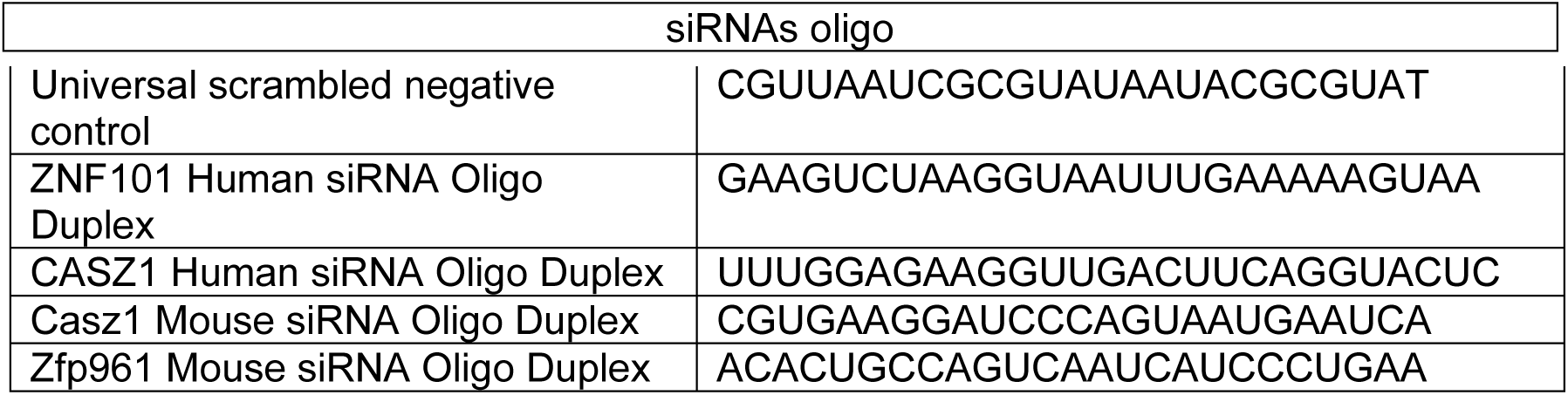

| Reagents |  |  |
| --- | --- | --- |
| Items | Vendor | Catalog # |
| Hsa-miR-541-3p | Thermo Fisher Scientific | 4464066, Assay ID#MC13119 |
| Hsa-AntimiR-541-3p | Thermo Fisher Scientific | 4464084, Assay ID# MH13119 |
| Hsa-miR-541-5p | Thermo Fisher Scientific | 4464066, Assay ID#MC12516 |
| miRNA Negative Control | Thermo Fisher Scientific | 4464058 |
| TaqMan™ miR-541-3p Assay | Thermo Fisher Scientific | 4427975, Assay ID#002201 |
| TaqMan™ miR-U6 Assay | Thermo Fisher Scientific | 4427975, Assay ID#001973 |
| TaqMan™ MicroRNA Reverse Transcription Kit | Thermo Fisher Scientific | 4366596 |
| TaqMan™ Universal Master Mix II, no UNG | Thermo Fisher Scientific | 4440043 |
| PowerTrack SYBR Green Master Mix | Thermo Fisher Scientific | A46012 |
| High-Capacity cDNA Reverse Transcription Kit | Thermo Fisher Scientific | 4374967 |
| TRIzol™ Reagent | Thermo Fisher Scientific | 15596026 |
| ELISA Pro: Human apoB | MABTECH | 3715-1HP-20 |
| Human Apolipoprotein A-I/ApoA1 DuoSet ELISA | R&D system | DY3664-05 |
| miRNA Target clone control vector | Genecopoeia | CmiT000001-MT05 |
| miRNA 3'UTR target expression clone for Human ZNF101 | Genecopoeia | HmiT071405-MT05 |
| miRNA 3'UTR target expression clone for Human CASZ1 | Genecopoeia | HmiT105330-MT05 |
| Custom promoter reporter clone for Human APOB | Genecopoeia | CS-HPRM30361-PG04-02 |
| Promoter reporter clone for Human APOA1 | Genecopoeia | HPRM25646-PG04 |
| Zfp961-AAVPrime™ Purified AAV Particles | Genecopoeia | AA08-MSH074798-AV03 |
| Casz1-AAVPrime™ Purified AAV Particles | Genecopoeia | AA08-MSH034915-AV03 |
| AAVPrime™ Purified AAV Particles for shRNA scrambled control | Genecopoeia | AA08-CSHCTR001-AV03 |
| AAV8-D377y-mPCSK9 (5e12 gc) | Vector biosystems Inc | 7600 |
| Secrete-Pair Dual Luminescence Assay |  | LF031 |
| Apolipoprotein A-I/ApoA1 Antibody | NOVUS | NBP2-15429 |
| Apolipoprotein B (APOB), Polyclonal Antibody | MyBiosource | MBS2006107 |
| Anti-Argonaute-2 antibody | Abcam | ab186733 |
| Pointe Scientific Triglycerides Liquid Reagent | Pointe Scientific | 23-666-411 |
| Pointe Scientific Cholesterol Liquid Reagents | Pointe Scientific | 23-666-200 |
| EndoFectin™ Max transfection Reagent | Genecopoeia | EF013 |
| Protease Inhibitor Cocktail | Sigma Aldrich | P1860 |
| RIPA buffer | Thermo Fisher Scientific | J63306 |
| Poloxamer 40716758 | Sigma Aldrich | 40716758 |
| Olive oil | Sigma Aldrich | 01514 |
| T0901317, LXR agonist | Abcam | ab142808 |
| ScintiSafe™ Econo 2 Cocktail | Thermo Fisher Scientific | FISSX21-5 |
| Cholesterol, [1,2-3H(N)] | Perkin Elmer | NET139001MC |

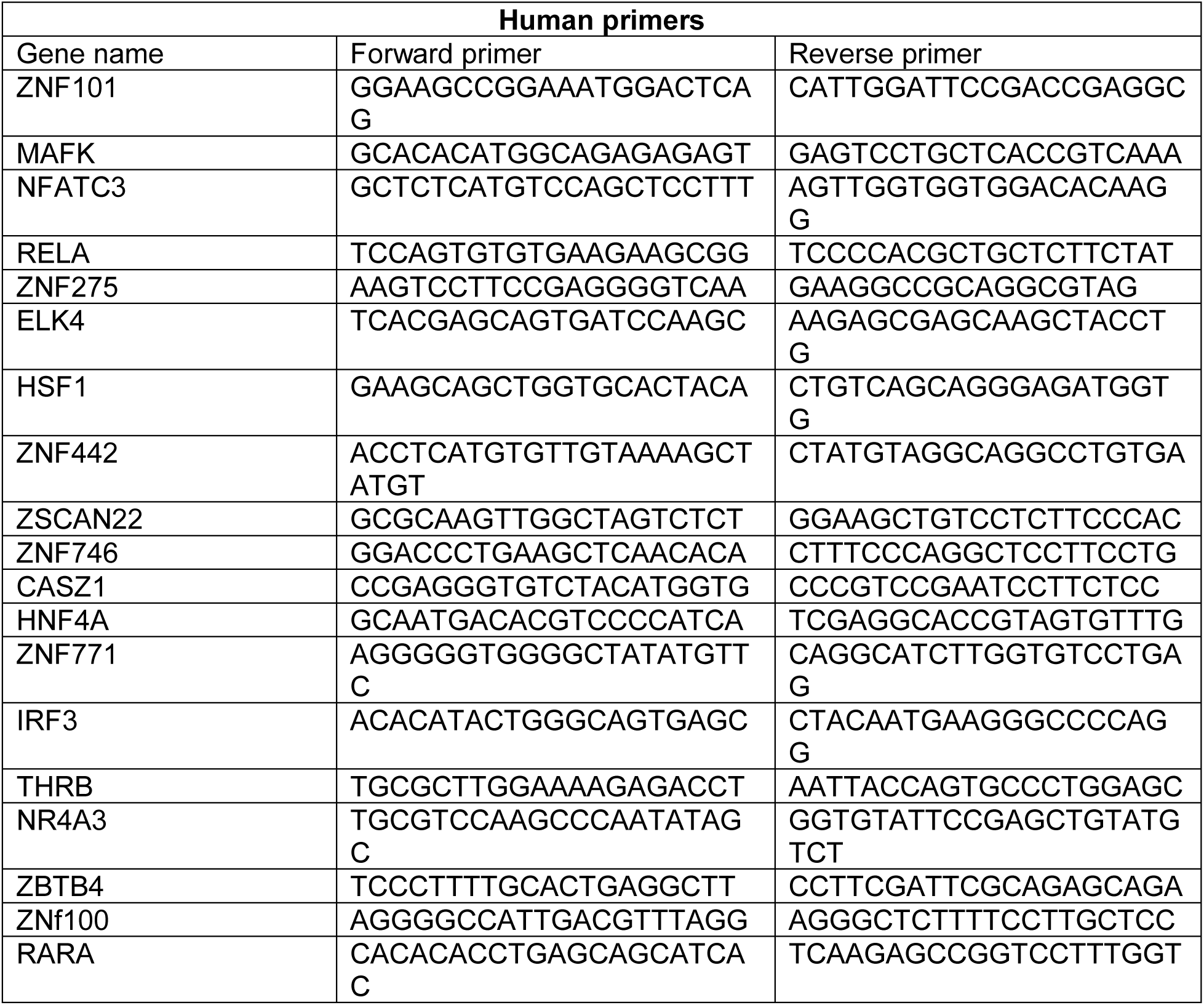
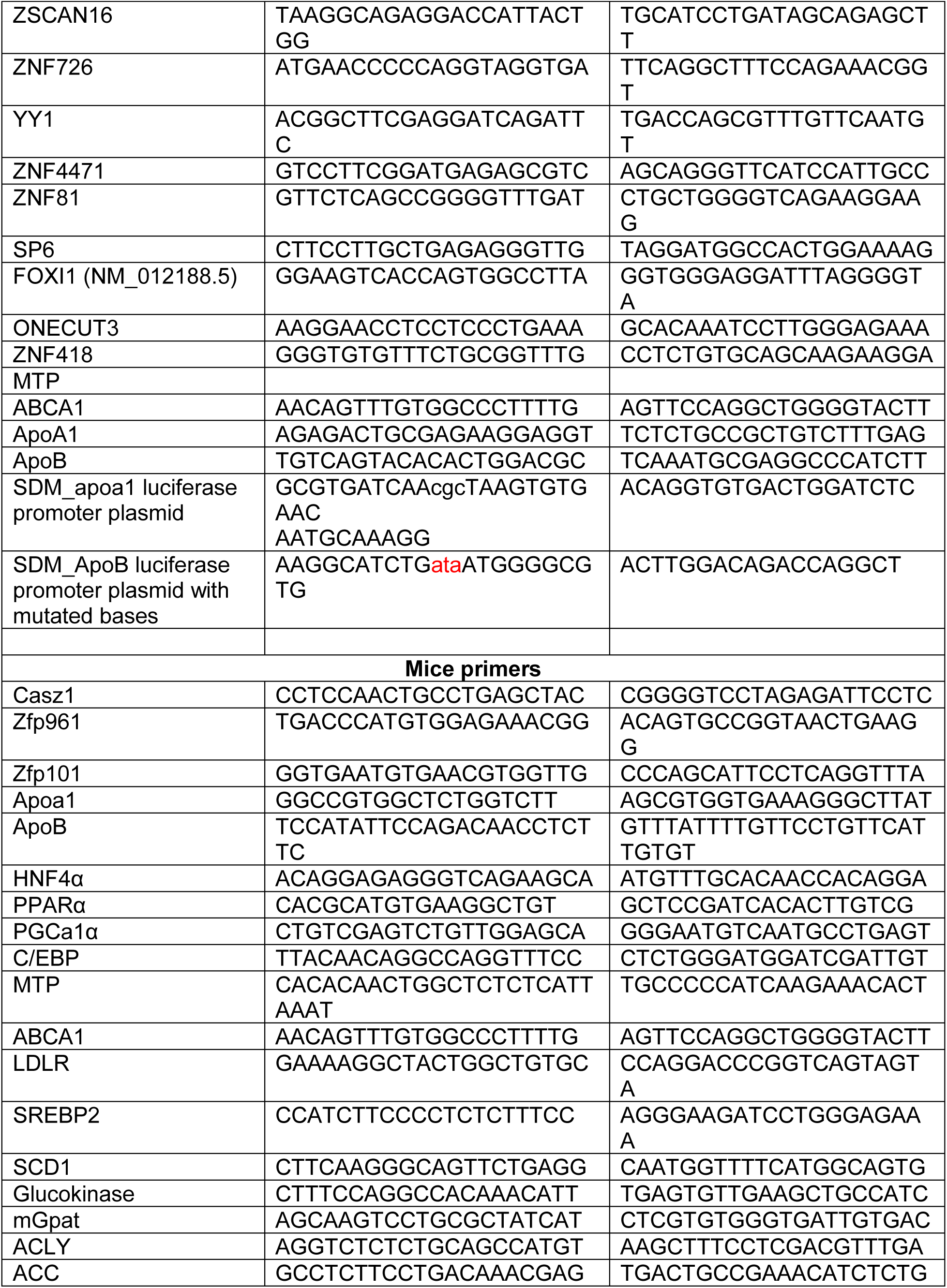
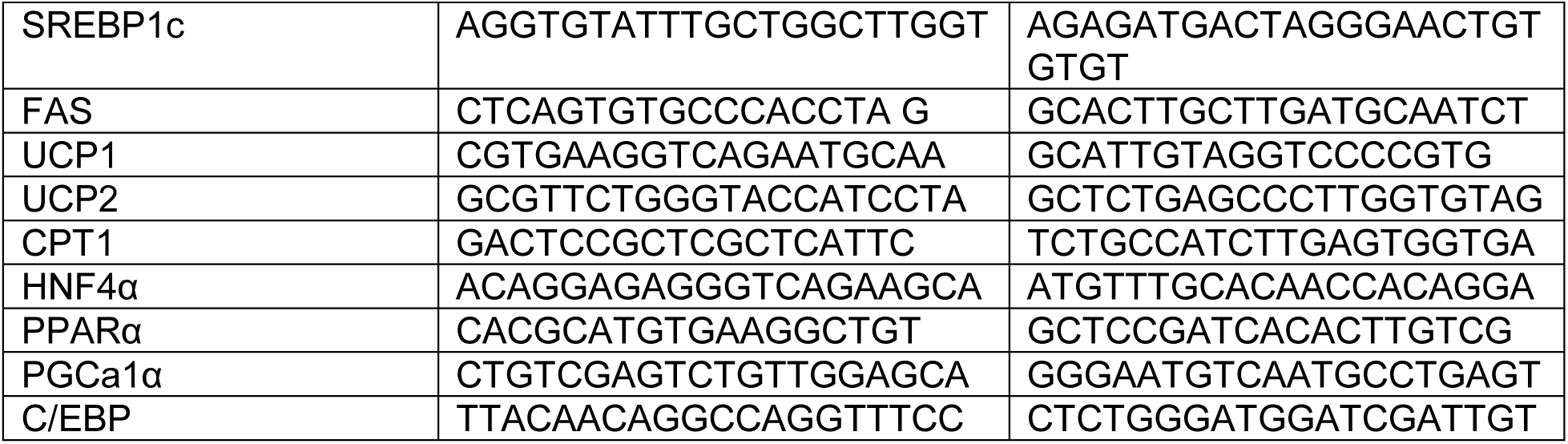

