## Supplementary Figures for "MicroRNA-541-3p alters lipoproteins to reduce atherosclerosis by degrading Znf101 and Casz1 transcription factors"

### Extended Data Fig 1

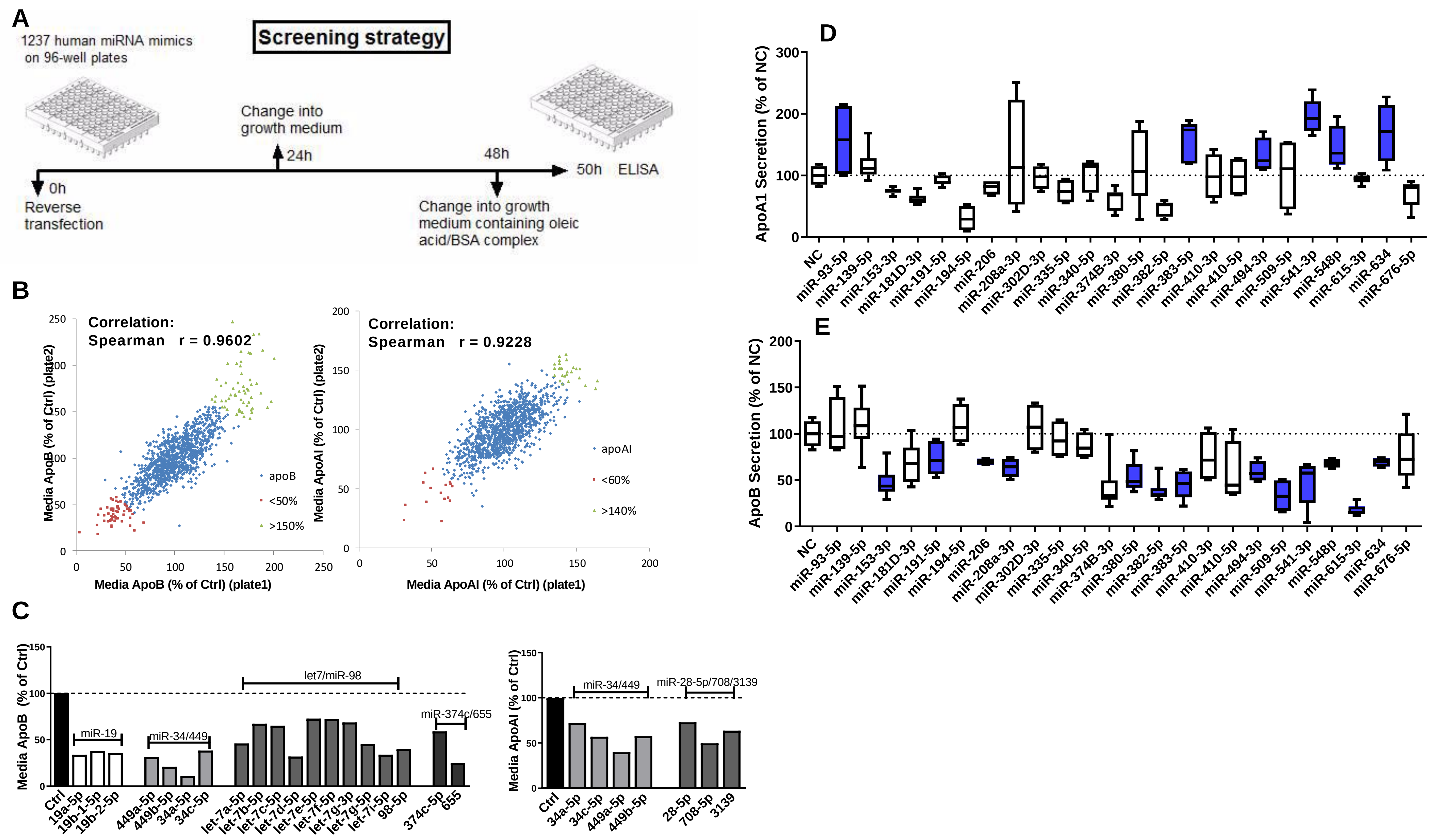

**Extended Data Fig 1. Screening a library to identify microRNAs regulating apoB and apoA1 secretion in human hepatoma Huh-7 cells.**

**(A)** Huh-7 cells were reverse transfected in duplicate plates with a human miRDIAN mimic 16.0 library (Dharmacon) of 1237 miRs at 50 nM. After 24 h, cells received complete media with 10% FBS. After another 24 h, cells were incubated with complete media containing 10% FBS and oleic acid/BSA complexes (0.4 mM/1.5%) for 2 h to avoid identification of miRs that affect posttranslational degradation of apoB. Media were used to quantify apoB and apoA1 levels by ELISA. Few wells in each plate were simultaneously transfected with negative control (Ctrl) or miR-30c (positive control, reduces apoB).

**(B)** Percentage change in media apoB and apoA1 in two plates exposed to the same miRs were compared with the negative control. Changes (%) in plate 1 are plotted against plate 2. Correlation between two plates with respect to changes in media apoB and apoA1 was determined.

**(C)** Different miR family members with the same seed sequence showed similar reductions in media apoB and apoA1 indicating internal consistency in the regulation of apoB and apoA1 secretion by family members.

**(D-E)** Second screening to identify miRs that simultaneously regulate ApoB and ApoA1. Huh-7 cells transfected in triplicate with 24 different miR mimics (100 nM). Media was changed after 48 h and collected after overnight incubation to quantify ApoB and ApoA1 by ELISA. (D) Compared to negative control (NC), six miRs significantly increased ApoA1 secretion, while (E) thirteen miRs downregulated ApoB secretion. Four miRs simultaneously regulated both ApoB and ApoA1.

Extended Data Fig 2

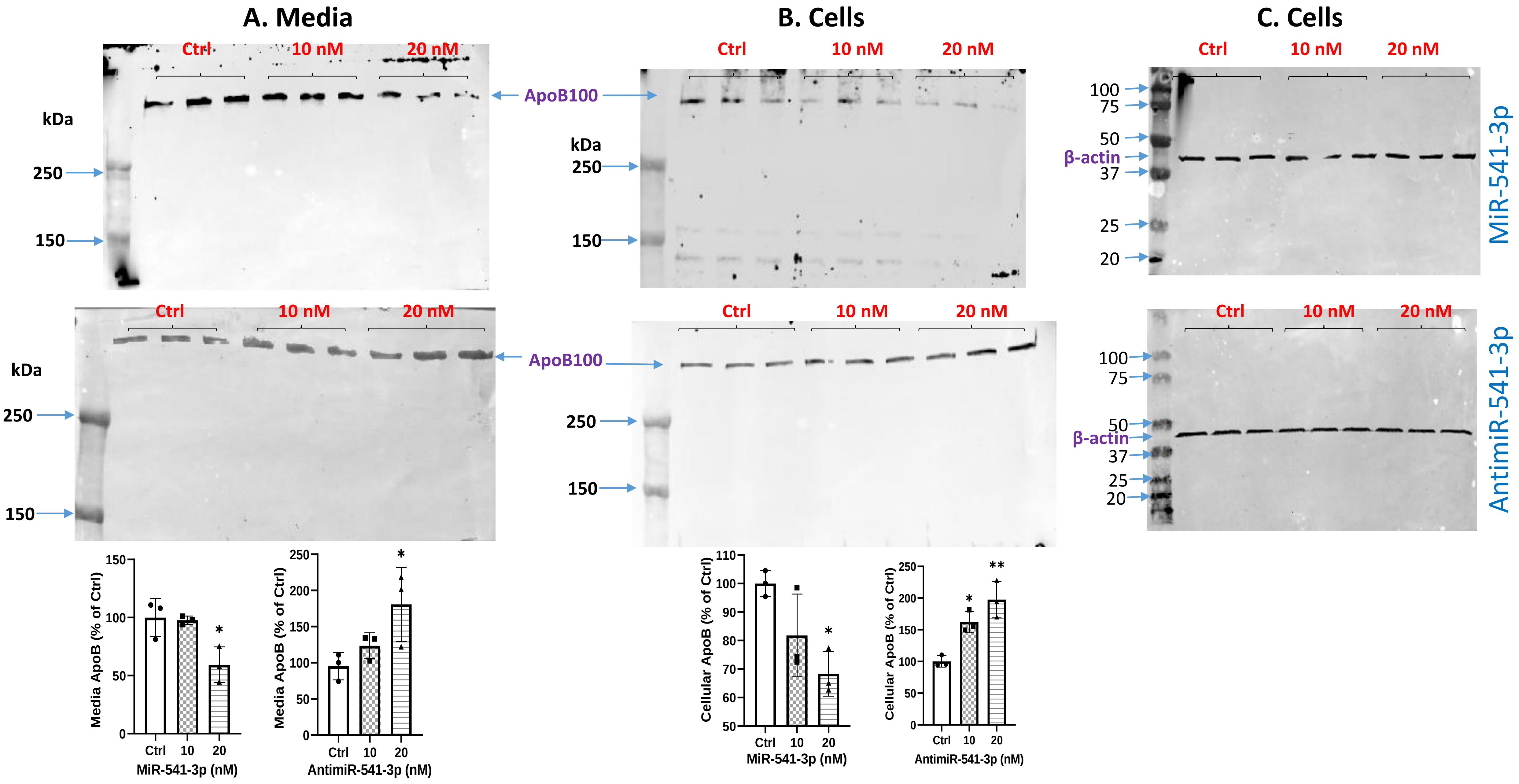

#### Extended Data Fig 2

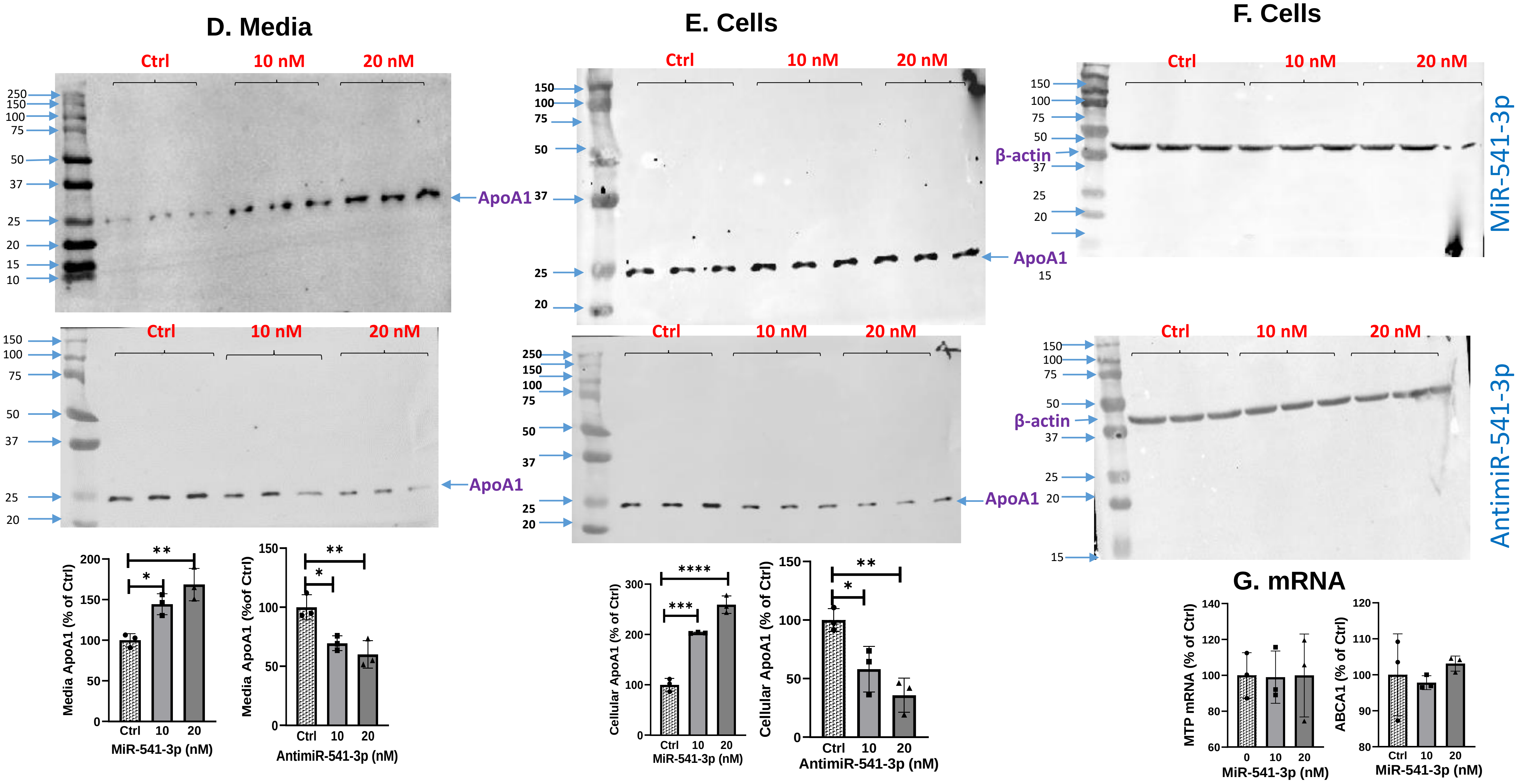

**Extended Data Fig 2. Regulation of apoB and apoA1 by miR-541-3p.**

(**A-C**) Full gels probing apoB100 in the (A) media and (B-C) cell lysates of Huh-7 cells treated with miR-541-3p mimics (top) or antimiR-541-3p (bottom), respectively. Bands were quantified by ImageJ and plotted as % of control (insets). The densitometric analysis showed significant reductions in apoB protein levels in cells and media of Huh-7 cells overexpressing miR-541-3p mimics and significant increases in cells overexpressing antimiR-541-3p. MiR-541-3p mimics and antimiR-541-3p had no effect on β-actin protein levels.

(**D-F**) Full gels and quantification of apoA1 bands in media (A) and cells (B) transfected with miR-541-3p mimics and antimiR-541-3p. Protein bands were quantified using ImageJ and normalized to β-actin (C) and plotted as % of control (bottom). MiR-541-3p mimics increase whereas antimiR-541-3p decreases apoA1 expression.

(**G**) Quantification of *MTP* and *ABCA1* mRNA levels in Huh-7 cells transfected with different amounts of miR-541-3p mimics.

#### Extended Data Fig 3

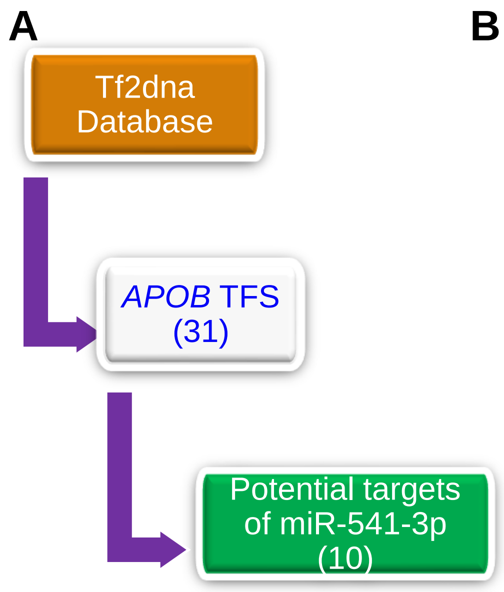


B

| Expression of TFs Regulating ApoB in Huh-7 cells | |
|---|---|
| Expressed | Not expressed |
| ZNF101 | ZNF442 |
| MAFK | ZSCAN22 |
| NFATC3 | ZNF746 |
| RELA | |
| ZNF275 | |
| ELK4 | |
| HSF1 | |

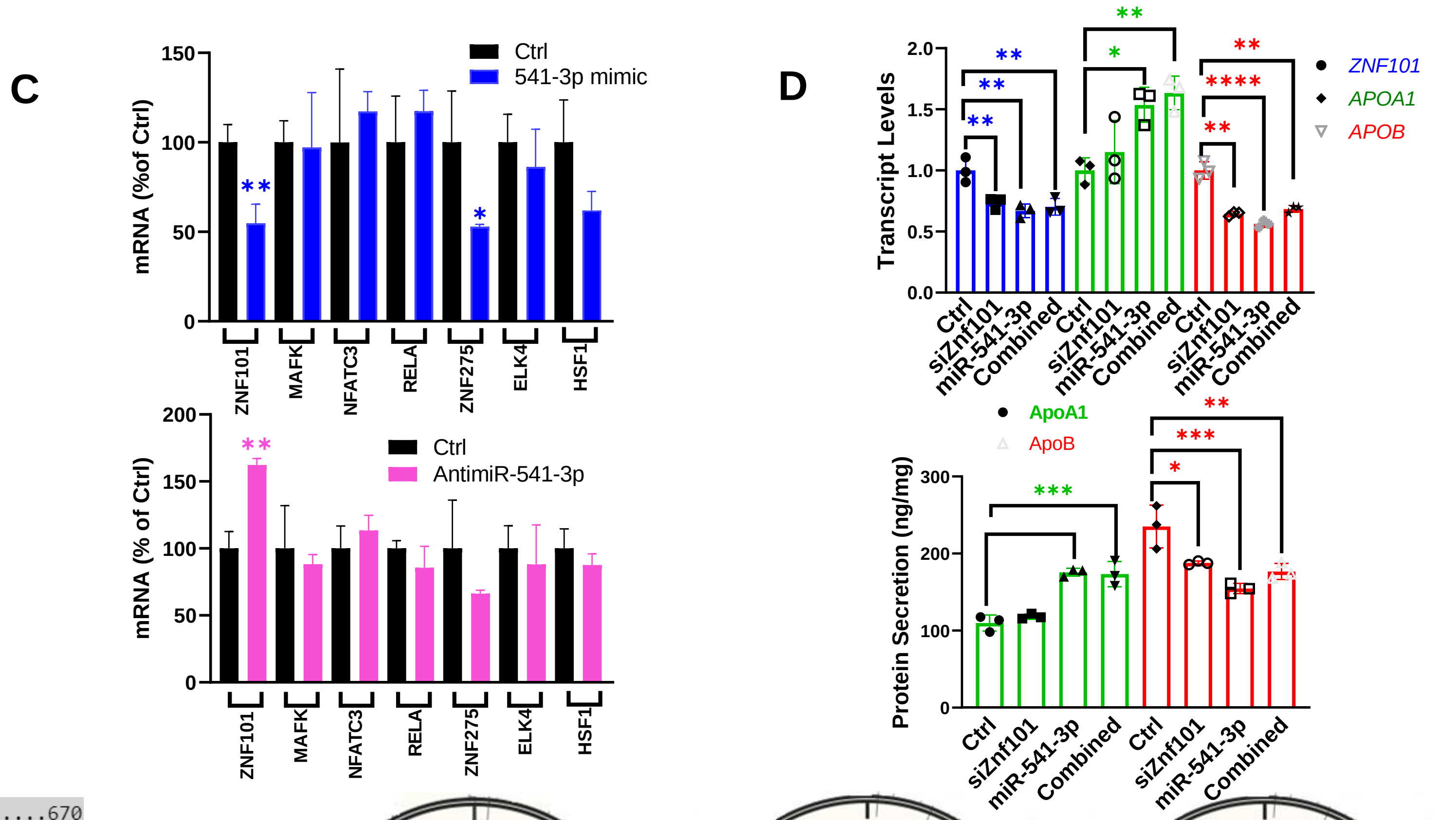


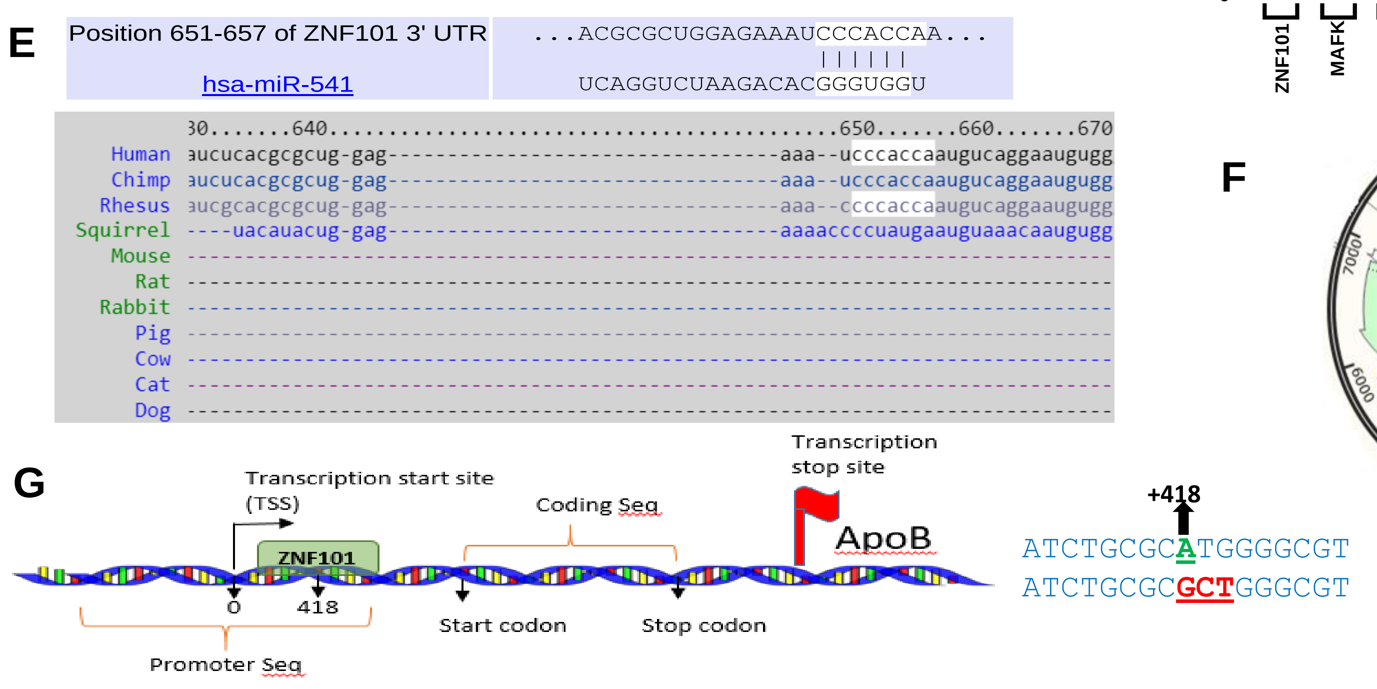


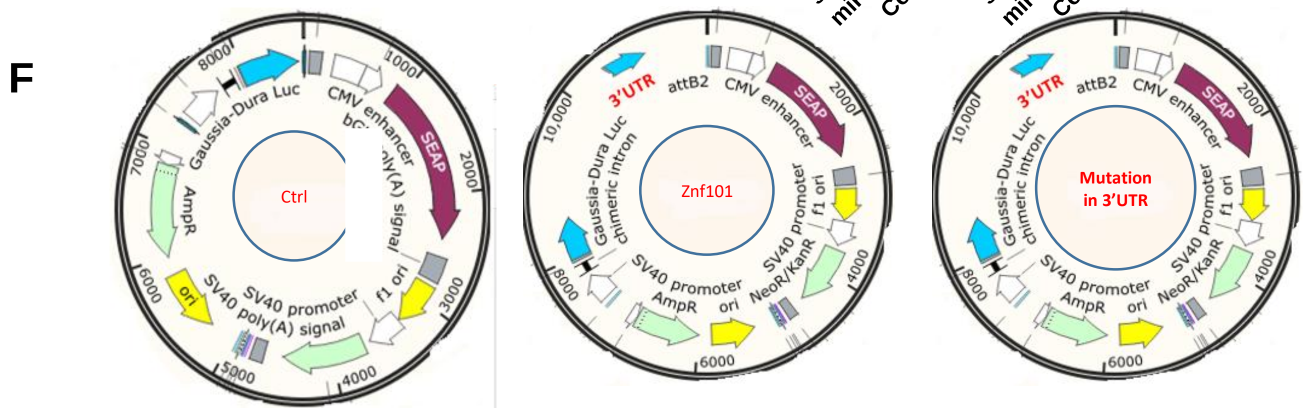


**Mutations in the seed complementary sequence of 3'-UTR of ZNF101**

5' CCCACCA 3'- ZNF101 sequence

5' CCCACAT 3'- Mutated (red) ZNF101 sequence

**Extended Data Fig 3. Identification of transcription factors (TFs) regulating apoB.**

(**A**) Tf2dna database was queried for TFs that could potentially regulate *APOB* gene expression. This identified 31 potential TFs that could regulate *APOB* expression. Ten of these TFs were identified as targets of miR-541-3p using Target Scan.

(**B**) Transcript levels of these 10 TFs were quantified in triplicate using specific primers. Seven TFs could be measured in Huh-7 cells.

(**C**) Huh-7 cells were transfected in triplicate with 20 nM miR-541-3p or antimiR-541-3p. After 48 h, changes in mRNA levels were quantified. Znf101 significantly reduced or increased, respectively, in cells transfected with miR-541-3p mimics or antimiR-541-3p.

(**D**) Huh-7 cells were transfected with miR-541-3p mimics (20 nM) or siZnf101 (28 nM), alone or in combination. After 48 h, cell lysates were used to measure different mRNA levels (left). Conditioned media was used to quantify apoB and apoA1 levels by ELISA (right).

(**E**) Target Scan predicted that the 3′-UTR of ZNF101 mRNA contains complementary bases that could pair with miR-541-3p seed sequence (top). Target Scan was used to determine the conservation of Znf101 3′-UTR in different species. Conserved sequences were found in the rhesus monkey and chimpanzee (bottom).

(**F**) Schematic diagrams of plasmids expressing Gaussia-Dura luciferase under the control of Znf101 3′-UTR obtained from Genecopoeia. The plasmid also constitutively expresses alkaline phosphatase under the control of cytomegalovirus (CMV) promoter and is used as a control. The highlighted bases were mutated in the wild-type plasmid.

(**G**) Schematic diagram showing potential Znf101 binding site in the *APOB* promoter. Transcription start site (TSS) and stop sites are identified. *APOB* promoter and coding sequences (seq) are shown (not to scale). Potential Znf101 binding site was mutated as shown in red.

### Extended Data Fig 4

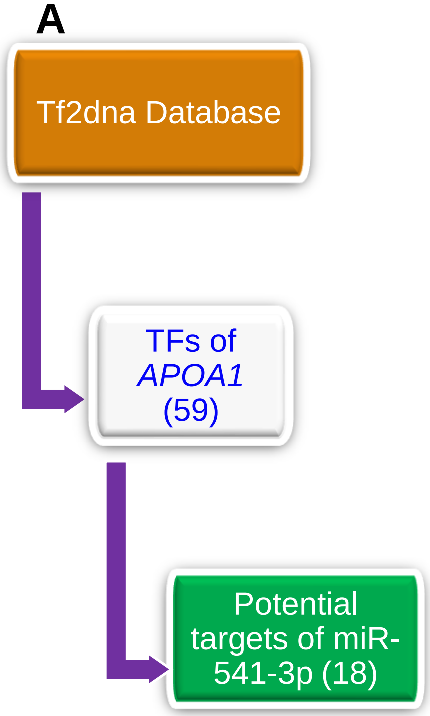


B

| Expression of TFs regulating ApoA1 in Huh-7 cells | |
|---|---|
| Expressed | Not expressed |
| HNF4A | SP6 |
| ZBTB4 | ZNF81 |
| IRF3 | FOXI1 |
| ZNF100 | ONECUT3 |
| THRB | ZNF418 |
| RARA | |
| NR4A3 | |
| ZSCAN16 | |
| ZNF726 | |
| ZNF471 | |
| YY1 | |
| CASZ1 | |
| ZNF771 | |

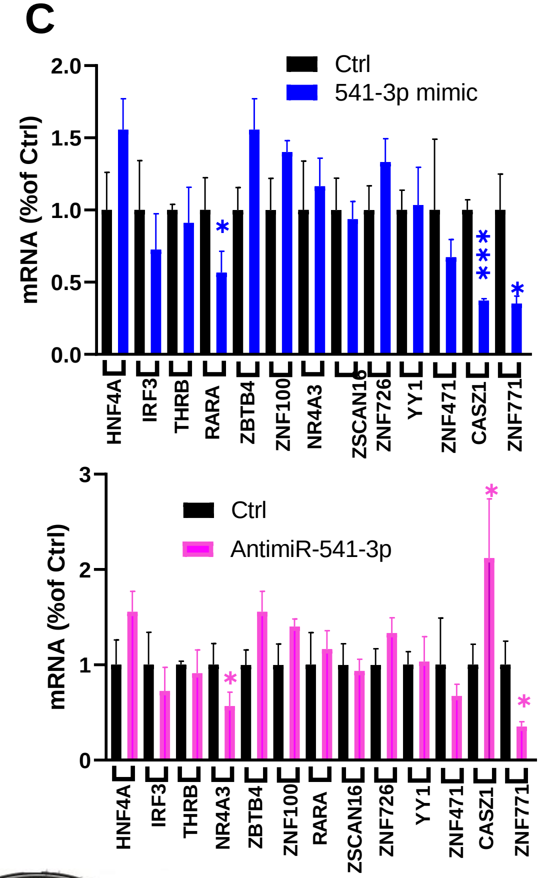


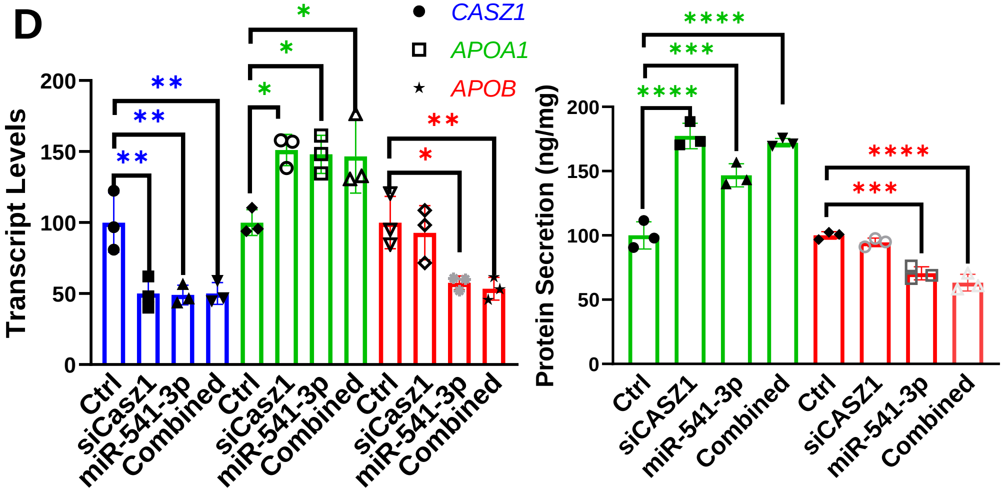


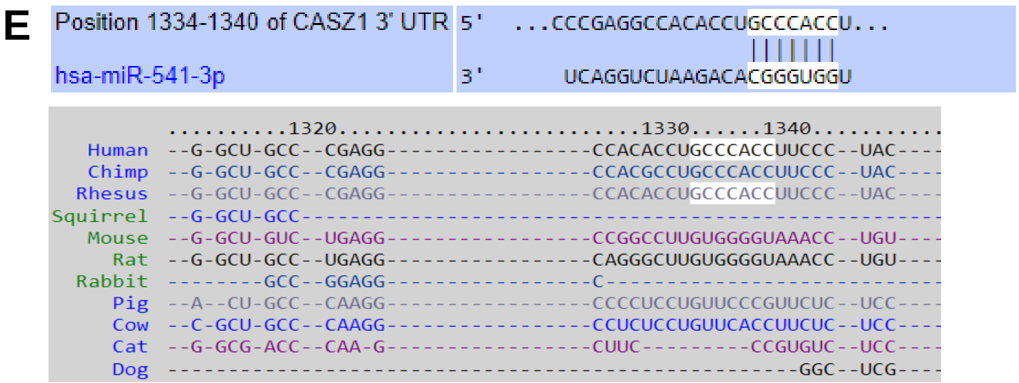


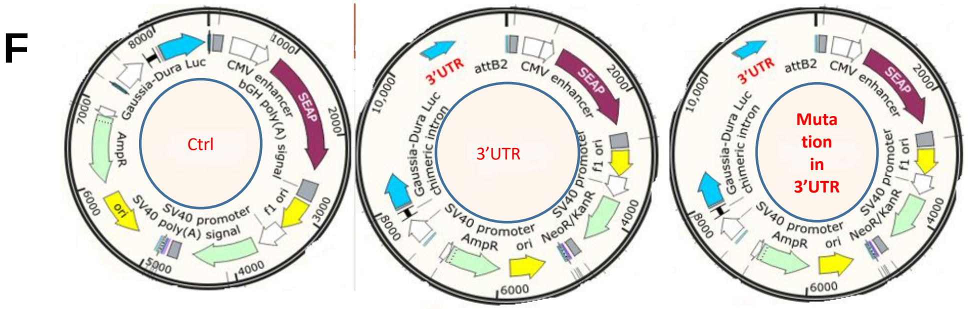


**Mutations in the seed complementary sequence 3'-UTR of Casz1:**

5' GCCCACC 3'- CASZ1 sequence

5' GCAATCC 3' Mutated (red) CASZ1 sequence

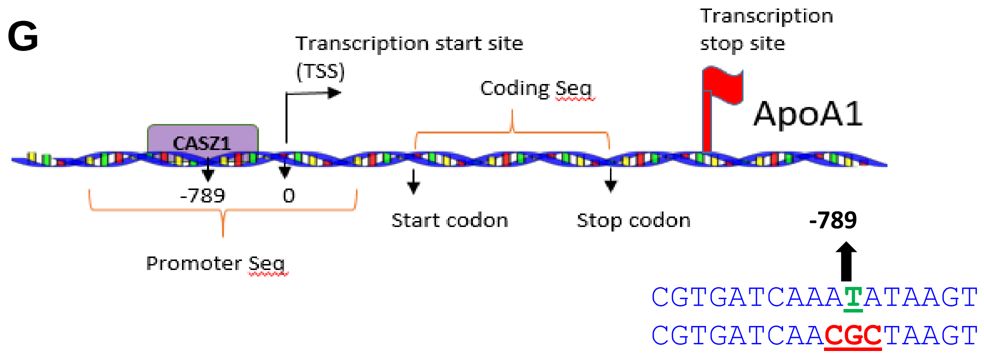

**Extended Data Fig 4. Identification of TFs regulating apoA1.**

(**A**) Tf2dna database was searched for TFs that could regulate *APOA1* gene expression. This identified 59 potential TFs. Target Scan predicted 18 of these TFs to be targets of miR-541-3p.

(**B**) mRNA levels of these TFs were quantified in triplicate by qRT-PCR. Huh-7 cells express 13 of these TFs.

(**C**) Huh-7 cells were transfected with 20 nM miR-541-3p mimics or antimiR-541-3p. After 48 h, changes in mRNA levels of different TFs were quantified in triplicate. Casz1 was significantly decreased and increased, respectively, in cells expressing miR-541-3p mimics or antimiR-541-3p.

(**D**) Huh-7 cells were transfected in triplicate with miR-541-3p mimics (20 nM) or siCasz1 (28 nM), alone or in combination. After 48 h, mRNA levels were quantified in cells (left), and protein levels in conditioned media (right) in triplicate. MiR-541-3p mimics and siCasz1, individually and combined, increased apoA1 levels to a similar extent indicating that they are in the same pathway.

(**E**) Target Scan 7.2 showed that Casz1 3′-UTR contains a complementary sequence that could base pair with miR-541-3p seed sequence (top). Clustal W alignment algorithms indicated conservation of the 3′-UTR of Casz1 in primates (bottom).

(**F**) Plasmids expressing luciferase with Casz1 3′-UTR were obtained from Genecopoeia. Casz1 3′-UTR was mutated as shown below in red.

(**G**) Schematic diagram showing potential Casz1 binding site in the *APOA1* promoter (not to scale). Potential binding site sequence was mutated as shown in red.

### Extended Data Fig 5

**A**

Hsa-miR-541-3p: 3′ UCAGGUCUAAGACACGGGUGGU 5′

Mmu-miR-541-3p: 3′ UCAUACCUAAGACACAAGCGGU 5′

**B**

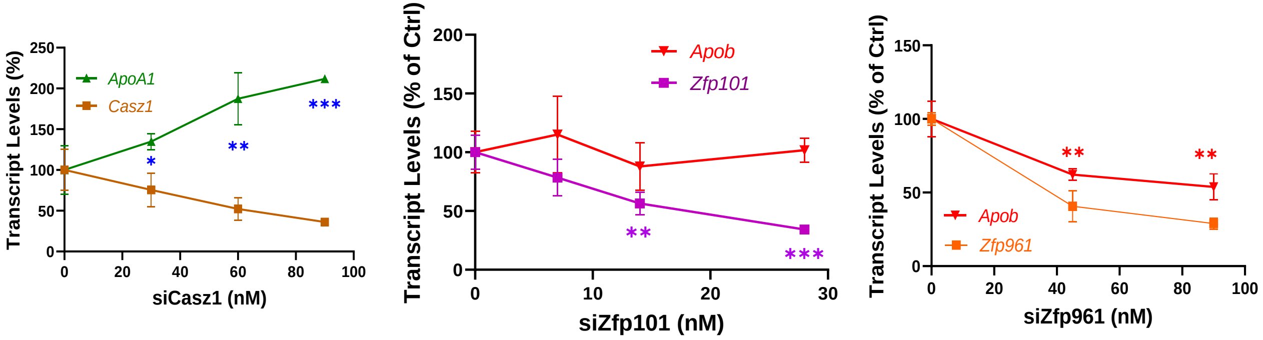


**C**

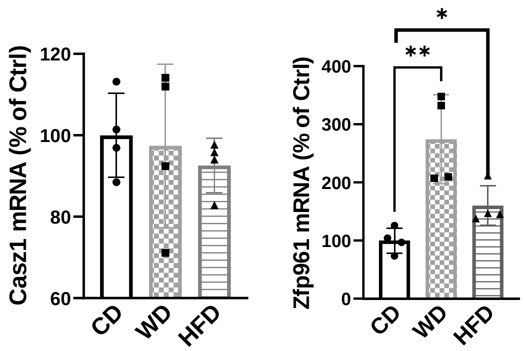

**Extended Data Fig 5. Mouse orthologs of human genes.**

(**A**) Human and mouse miR-541-3p sequences are compared to highlight sequence differences (red). Hsa-miR-541-3p and mmu-miR-541-3p are not identical. They contain three different bases in seed and non-seed sequences each.

(**B**) Identification of mouse orthologs of human Znf101 and Casz1 and their role in the regulation of mouse apoB and apoA1 expression. Mouse liver AML12 cells were transfected in triplicate with different concentrations of mouse siCasz1 (left), siZfp101 (middle) or siZfp961 (right). After 48 h, the indicated mouse mRNAs were quantified in triplicate. These studies identified *Casz1* and *Zfp961* as functional mouse orthologs of human *CASZ1* and *ZNF101* genes. * $P < 0.05$; ** $P < 0.01$; ***$P < 0.001$; ****$P < 0.0001$.

(**C**) Male C57Bl6 mice (*n* = 4, 5 months old) were fed chow (CD), Western (WD) or obesogenic (HFD) diets for 13 weeks. Livers were collected to measure mRNA levels. Zfp961 expression increased in the livers of high fat diet fed mice. One-way ANOVA, * $P < 0.05$, **$P < 0.001$.

### Extended Data Fig 6

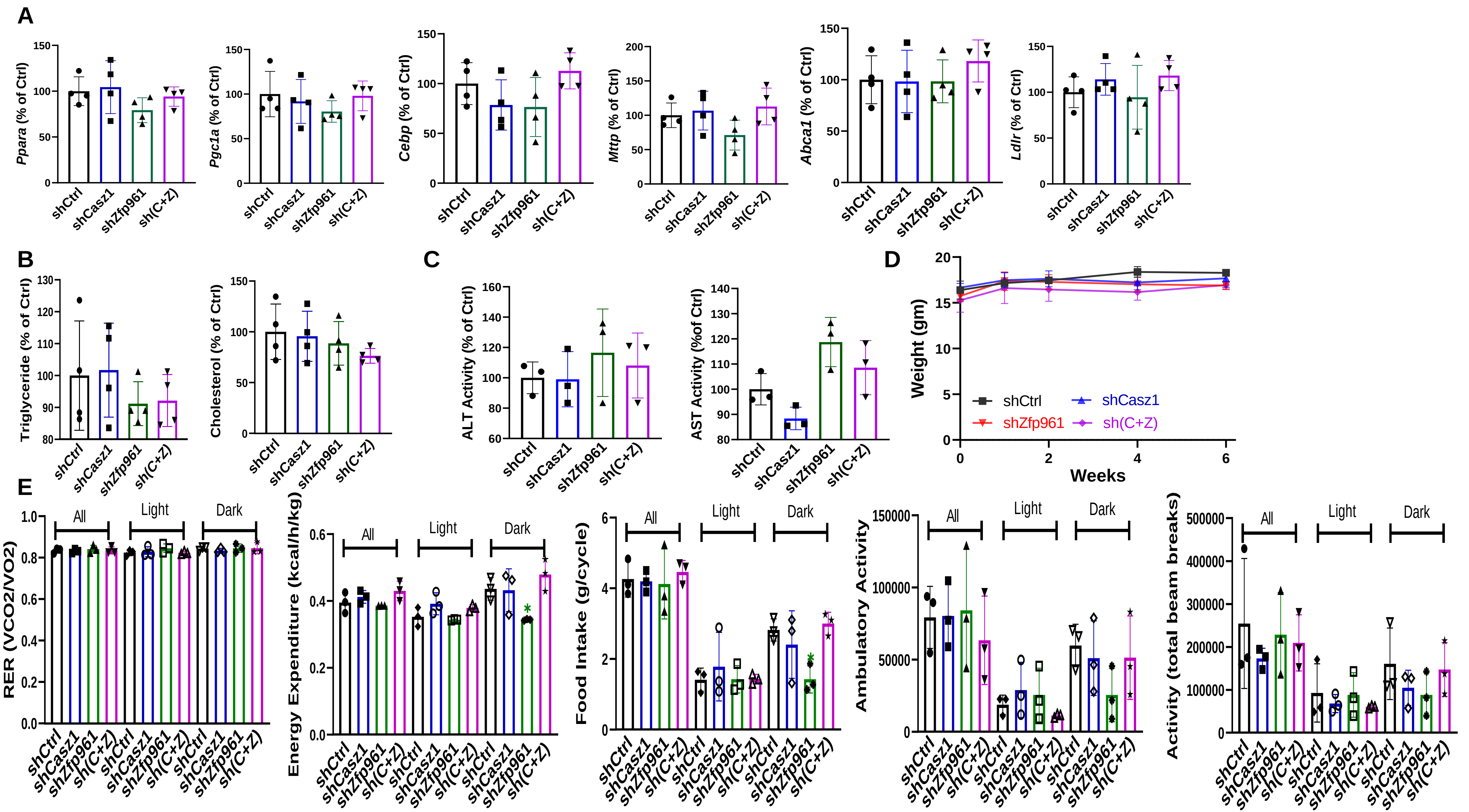

**Extended Data Fig 6. Effect of hepatic Casz1 and Zfp961 knockdown on different parameters in mice.**

Mice (C57Bl6J, female, 2.5-month-old) were transduced with AAV8 ($2.5 \times 10^{11}$ gc/mouse) expressing shControl (shCtrl, *n* = 4), shCasz1 (*n* = 4) or shZfp961 (*n* = 4), alone or combined (sh(C+Z), *n* = 5), and started on a Western diet.

(**A**) After 6 weeks, livers were collected to measure in triplicate different TFs and lipid metabolism genes. KD of different genes had no effect on their expression.

(**B**) After 6 weeks, lipids were extracted from liver slices and triglyceride and cholesterol levels were measured in triplicate and normalized with protein levels. KD of different genes had no effect on hepatic lipids.

(**C**) Plasma was used to measure ALT and AST activities in triplicate. These enzyme activities were unaffected by KD of Casz1 or Zfp961.

(**D**) Weight gain in mice was monitored over the course of  6 weeks. No significant differences were observed.

(**E**) After 5 week of injections, mice were placed in CLAMS to measure different physiological parameters. No significant differences were noted.

**Extended Data Fig 7**

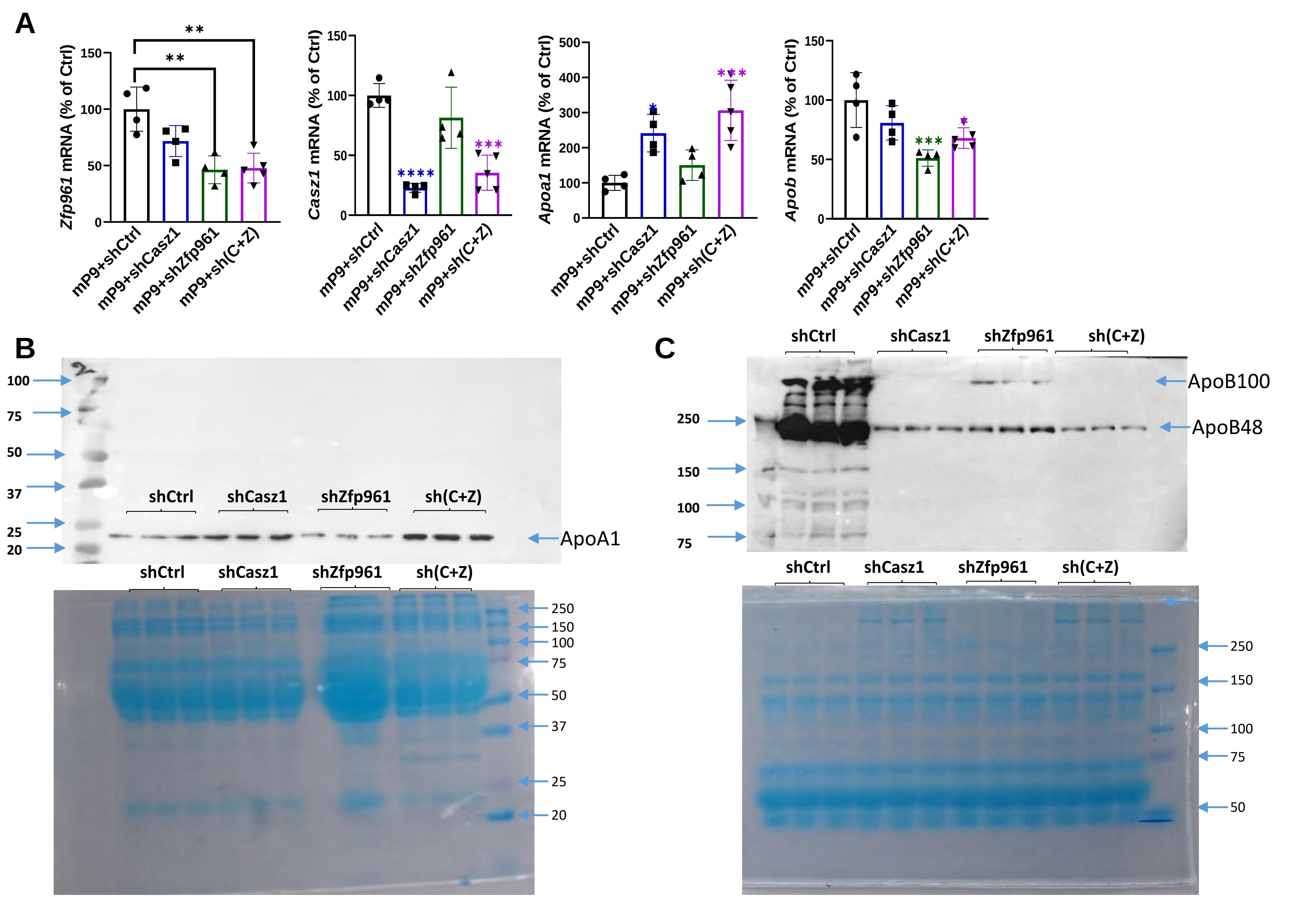

**Extended Data Fig 7. Effect of knockdown of different TFs on atherosclerosis.**

All mice received AA8 expressing mutant mouse PCSK9. In addition, mice were transduced with shCtrl, shCasz1, shZnf101, or shCasz1+shZnf101, and started on a Western diet.

(**A**) After 4 months, livers were collected to measure mRNA levels in triplicates.

(**B**) Detection of plasma apoA1 (top) by western blotting. Plasma (1 µL) was separated on a 10% gel, transferred, and probed with anti-apoA1 antibodies. Plasma (1 µL) was separated and stained with Coomassie blue (bottom) for control.

(**C**) Detection of apoB by western blotting (top) after separating plasma (1 µL) on a 6% gel. Total plasma was separated and stained with Coomassie blue (bottom) for control.

### Extended Data Fig 8

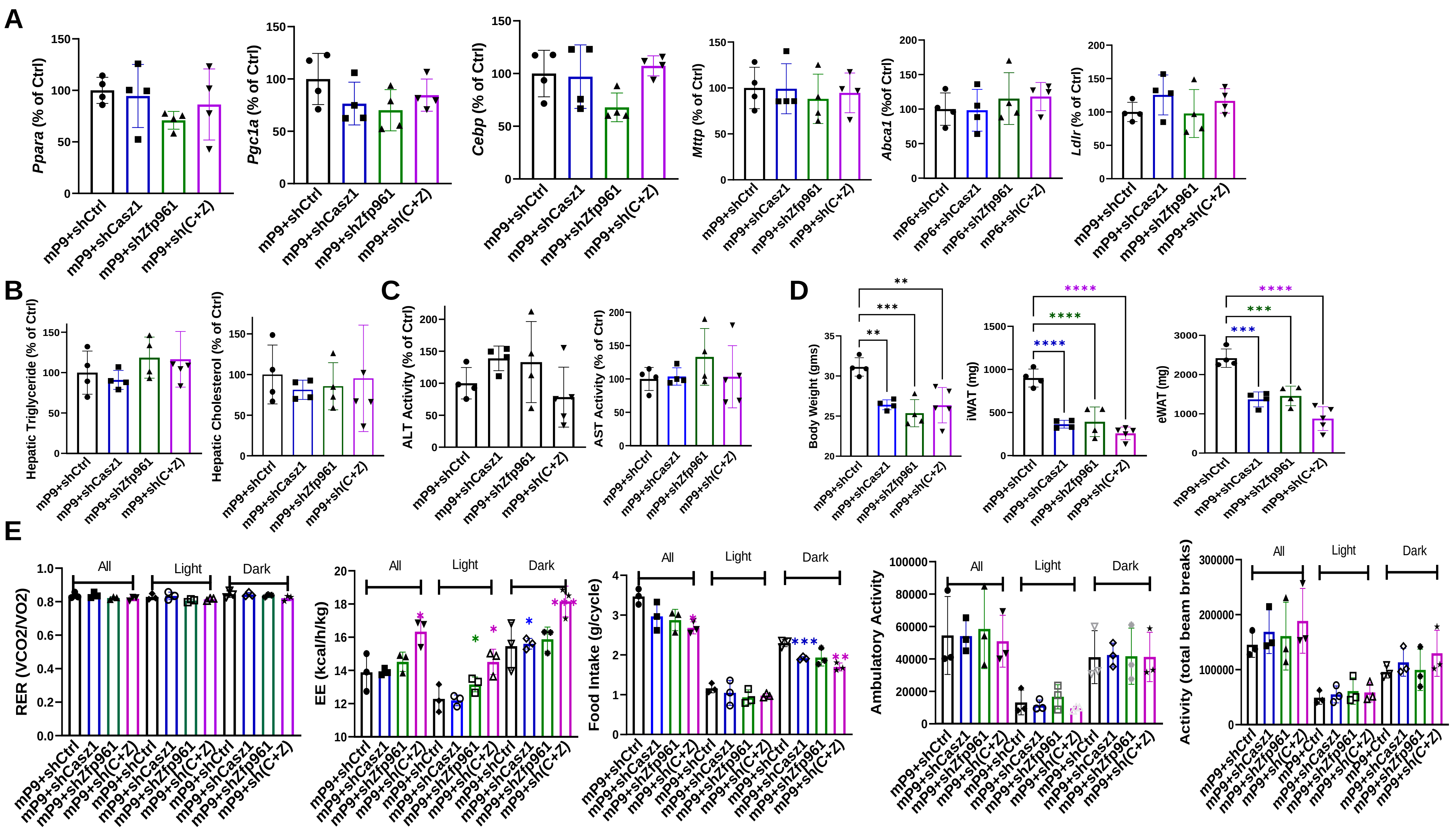

**Extended Data Fig 8. Effect of knockdown of different TFs on physiological parameters.** Mice were transduced with adenoviruses expressing gain-of-function mutant Pcsk9 and those expressing different shRNAs as in Fig 5.

(**A**) After 4 months, livers were collected to measure TFs and lipid metabolism genes in triplicates.

(**B**) Lipids were measured in triplicates in livers and normalized to protein levels. Knockdown of different TFs had no effect on hepatic lipids.

(**C**) Plasma was used to measure AST/ALT activities in triplicates.

(**D**) At the end, different tissues were collected. Total body and adipose tissue weights in different knockdown mice were lower than in control mice (right). No differences in other organ weights were observed.

(**E**) After 3 months, mice were placed in CLAMS (comprehensive laboratory animal monitoring system) to monitor physiological indices.

**Extended Data Fig 9**

A

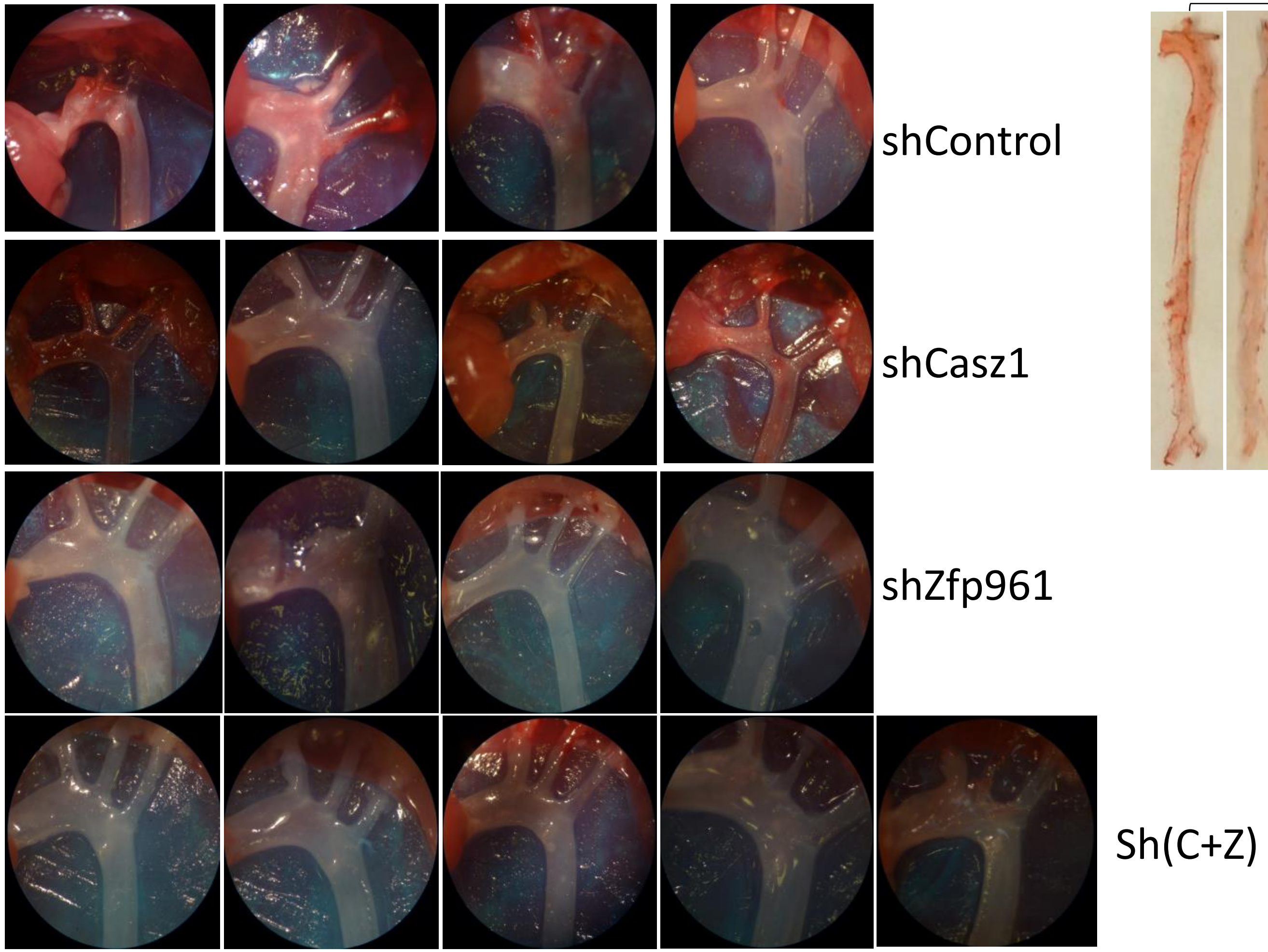


B

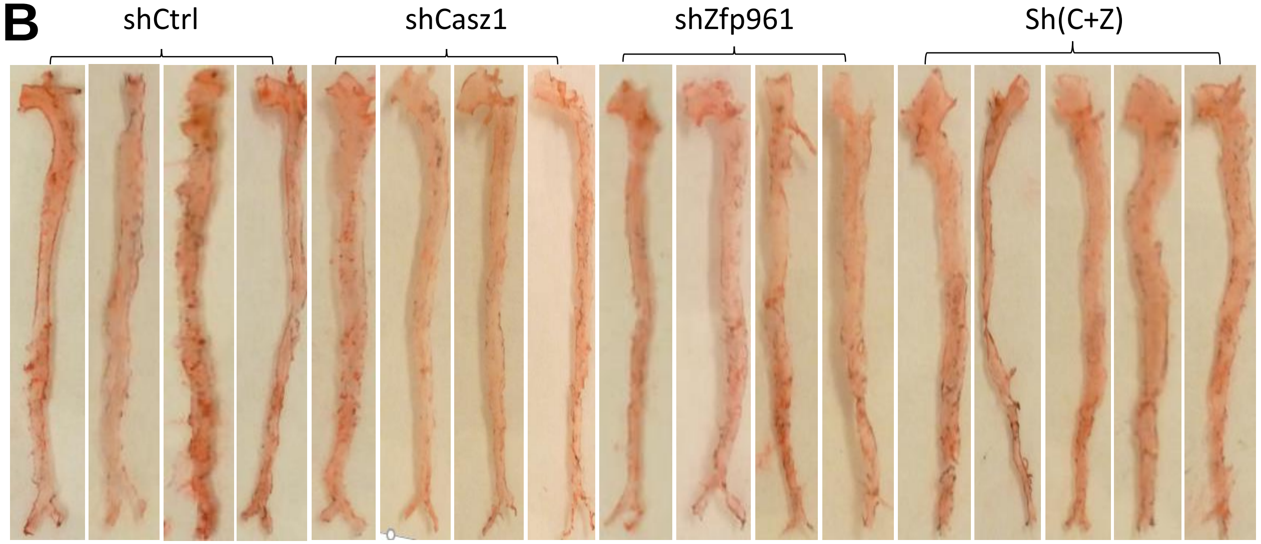

**Extended Data Fig 9. Effect of knockdown of different TFs on atherosclerosis.**

Mice were transduced with adenoviruses expressing gain-of-function mutant Pcsk9 and those expressing different shRNAs as in Fig 5.

(**A**) Aortas with major arteries were dissected from all mice, photographed, and presented as a collage.

(**B**) Whole aortas were stained with Oil Red O and all images were compiled.

**Extended Data Fig 10**

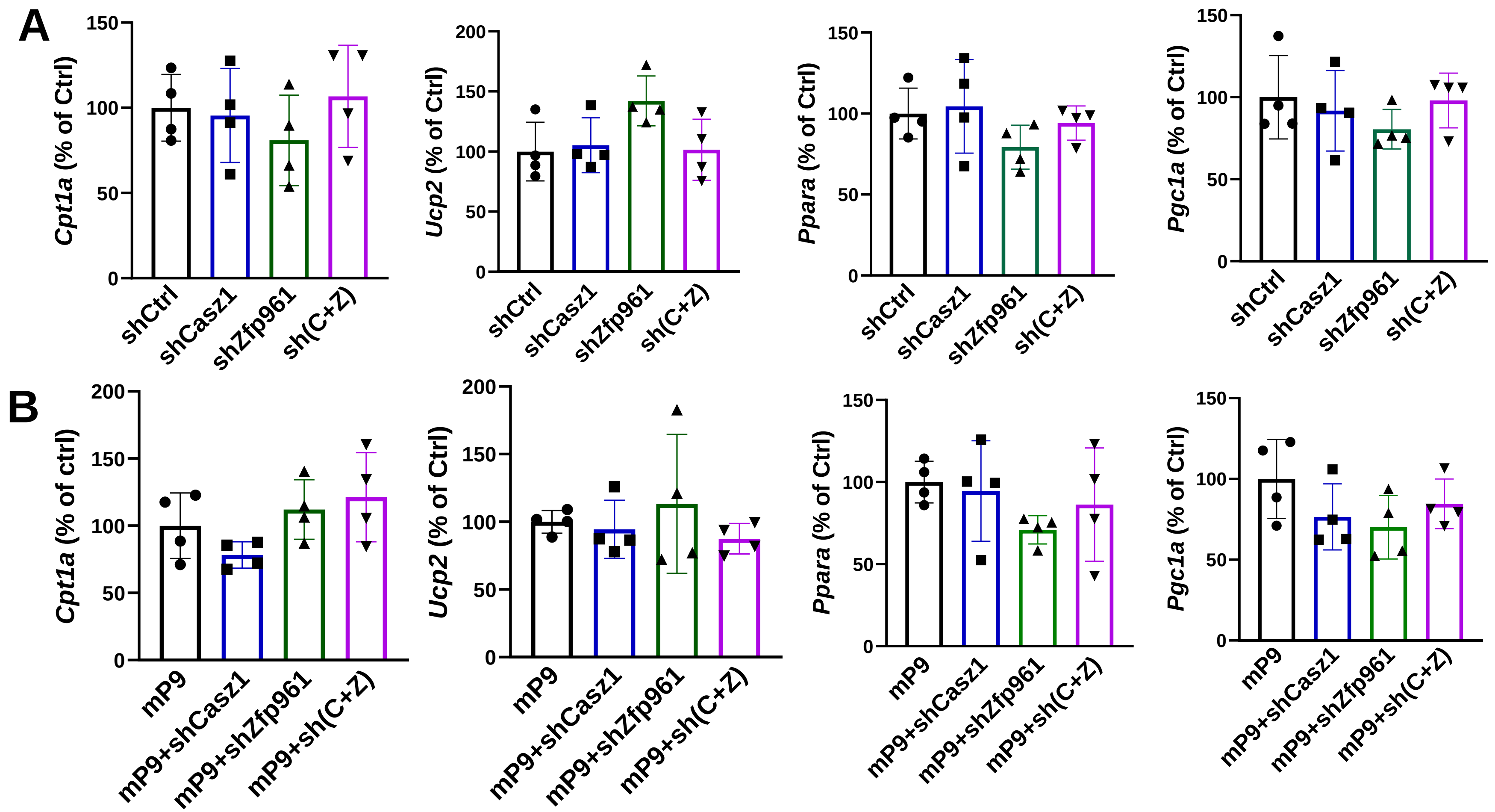

### Extended Data Fig 10

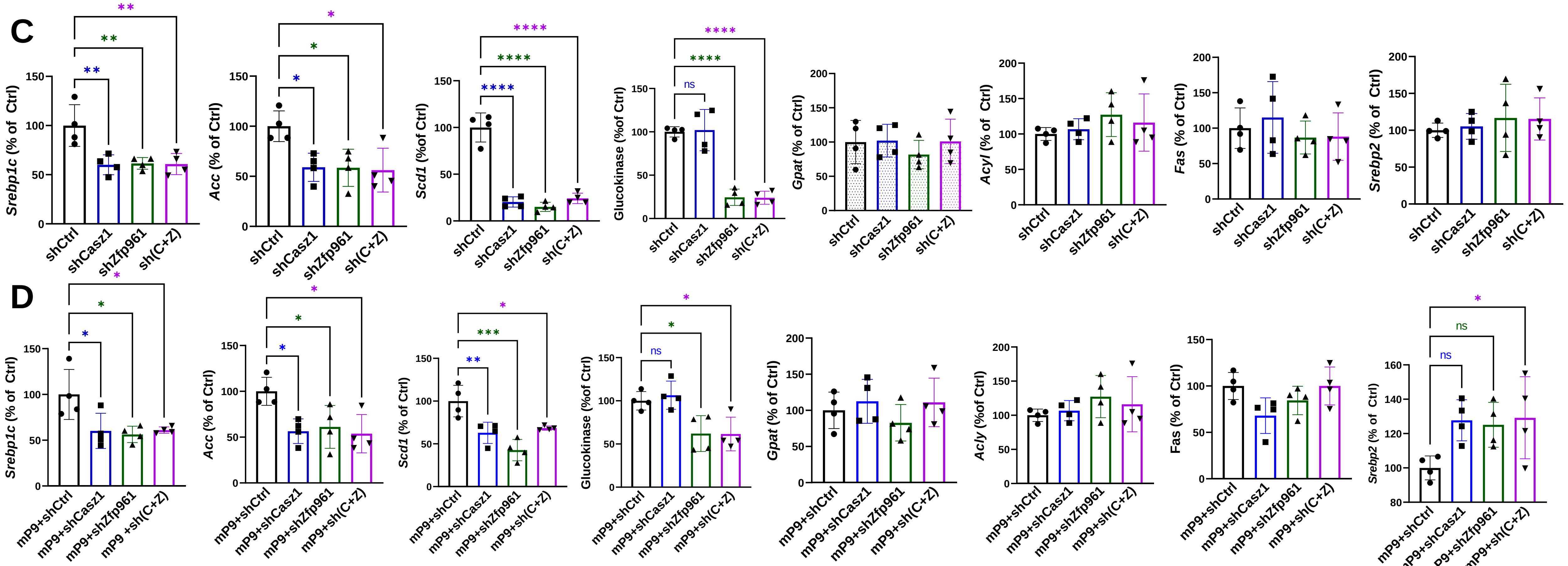

**Extended Data Fig 10. Hepatic knockdown of Casz1 and Znf961 has no effect on the mRNA levels of genes involved in β-oxidation, but reduces the expression of genes in lipogenesis.**

Mice were transduced with viruses as in Fig 4 (A, C) and Fig 5 (B, D). Livers from these mice were used to quantify in triplicate different mRNAs involved in β-oxidation (A, B) and lipogenesis (C, D).

**Extended Data Fig 11**

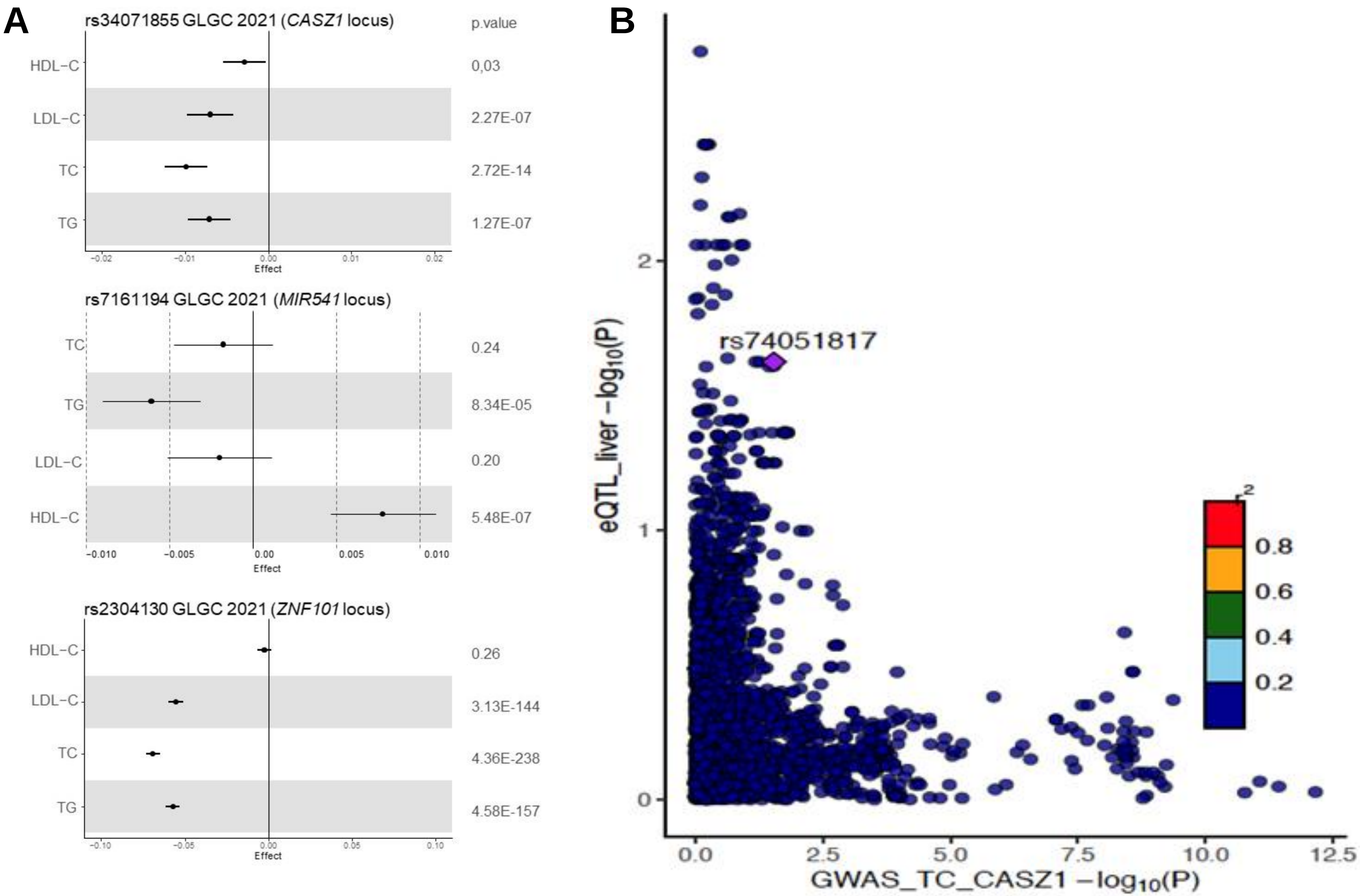

**Extended Data Fig 11. Association of variants in the *MIR541*, *CASZ1,* and *ZNF101* loci with plasma lipids and lipoproteins in GLGC.**

(A) Forrest plots show the associations (beta ± standard error) of the top associated SNP with plasma lipids and apolipoproteins levels in GLGC.

(B) LocusCompare plot shows the relationship between SNPs in the *CASZ1* locus (±100kb) and their association with total cholesterol plasma levels in the x-axis with the gene expression of *CASZ1* in the liver (eQTL_liver-log10(P) from GTEx dataset V7) in the y-axis.
