## Supplementary Table 1 for "MicroRNA-541-3p alters lipoproteins to reduce atherosclerosis by degrading Znf101 and Casz1 transcription factors"

**Supplementary Table I. The effect of 1237 miRs on apoB and apoAI secretion in Huh7 cells.**
**Medium apoB and apoAI were normalized against protein levels and calculated to percentages of Scr control.**

| | | | apoB | | apoAI | |
|---|---|---|---|---|---|---|
| Mature Name | Mature Accession | Mature Sequence | plate1 | plate2 | plate1 | plate2 |
| hsa-let-7a-2-3p | MIMAT0010195 | CUGUACAGCCUCCUAGCUUUCC | 110.79 | 94.94 | 114.44 | 102.87 |
| hsa-let-7a-3p | MIMAT0004481 | CUAUACAAUCUACUGUCUUUC | 112.31 | 86.33 | 73.30 | 59.51 |
| hsa-let-7a-5p | MIMAT0000062 | UGAGGUAGUAGGUUGUAUAGUU | 35.73 | 56.36 | 72.34 | 83.45 |
| hsa-let-7b-3p | MIMAT0004482 | CUAUACAACCUACUGCCUUCCC | 105.54 | 109.86 | 90.36 | 91.48 |
| hsa-let-7b-5p | MIMAT0000063 | UGAGGUAGUAGGUUGUGUGGUU | 77.29 | 57.43 | 107.87 | 115.59 |
| hsa-let-7c | MIMAT0000064 | UGAGGUAGUAGGUUGUAUGGUU | 67.69 | 62.76 | 78.38 | 70.12 |
| hsa-let-7c* | MIMAT0004483 | UAGAGUUACACCCUGGGAGUUA | 89.15 | 94.94 | 114.44 | 107.62 |
| hsa-let-7d-3p | MIMAT0004484 | CUAUACGACCUGCUGCCUUUCU | 158.43 | 246.59 | 91.70 | 95.20 |
| hsa-let-7d-5p | MIMAT0000065 | AGAGGUAGUAGGUUGCAUAGUU | 32.34 | 31.45 | 62.74 | 39.59 |
| hsa-let-7e-3p | MIMAT0004485 | CUAUACGGCCUCCUAGCUUUCC | 125.04 | 144.66 | 96.92 | 102.60 |
| hsa-let-7e-5p | MIMAT0000066 | UGAGGUAGGAGGUUGUAUAGUU | 78.48 | 67.11 | 89.71 | 77.22 |
| hsa-let-7f-1-3p | MIMAT0004486 | CUAUACAAUCUAUUGCCUUCCC | 122.09 | 92.95 | 100.81 | 95.83 |
| hsa-let-7f-2-3p | MIMAT0004487 | CUAUACAGUCUACUGUCUUUCC | 138.71 | 148.89 | 93.54 | 97.85 |
| hsa-let-7f-5p | MIMAT0000067 | UGAGGUAGUAGAUUGUAUAGUU | 72.94 | 71.62 | 60.78 | 71.53 |
| hsa-let-7g-3p | MIMAT0004584 | CUGUACAGGCCACUGCCUUGC | 69.96 | 67.38 | 110.83 | 99.48 |
| hsa-let-7g-5p | MIMAT0000414 | UGAGGUAGUAGUUUGUACAGUU | 55.08 | 35.56 | 91.47 | 83.59 |
| hsa-let-7i-3p | MIMAT0004585 | CUGCGCAAGCUACUGCCUUGCU | 86.46 | 92.51 | 108.30 | 108.19 |
| hsa-let-7i-5p | MIMAT0000415 | UGAGGUAGUAGUUUGUGCUGUU | 35.04 | 32.74 | 64.23 | 51.90 |
| hsa-miR-1 | MIMAT0000416 | UGGAAUGUAAAGAAGUAUGUAU | 144.14 | 142.57 | 125.78 | 120.96 |
| hsa-miR-100-3p | MIMAT0004512 | CAAGCUUGUAUCUAUAGGUAUG | 102.52 | 82.20 | 100.81 | 113.61 |
| hsa-miR-100-5p | MIMAT0000098 | AACCCGUAGAUCCGAACUUGUG | 107.18 | 111.45 | 102.86 | 80.38 |
| hsa-miR-101-3p | MIMAT0000099 | UACAGUACUGUGAUAACUGAA | 107.83 | 106.29 | 94.22 | 81.17 |
| hsa-miR-101-5p | MIMAT0004513 | CAGUUAUCACAGUGCUGAUGCU | 47.51 | 60.73 | 67.47 | 70.52 |
| hsa-miR-103a-2-5p | MIMAT0009196 | AGCUUCUUUACAGUGCUGCCUUG | 87.04 | 114.00 | 94.41 | 92.61 |
| hsa-miR-103a-3p | MIMAT0000101 | AGCAGCAUUGUACAGGGCUAUGA | 93.20 | 105.24 | 79.17 | 81.24 |
| hsa-miR-103b | MIMAT0007402 | UCAUAGCCCUGUACAAUGCUGCU | 91.33 | 92.95 | 125.87 | 122.49 |
| hsa-miR-105-3p | MIMAT0004516 | ACGGAUGUUUGAGCAUGUGCUA | 162.70 | 201.90 | 107.31 | 122.44 |
| hsa-miR-105-5p | MIMAT0000102 | UCAAAUGCUCAGACUCCUGUGGU | 89.81 | 80.09 | 82.59 | 115.36 |
| hsa-miR-106a-3p | MIMAT0004517 | CUGCAAUGUAAGCACUUCUUAC | 100.65 | 91.42 | 71.76 | 71.72 |
| hsa-miR-106a-5p | MIMAT0000103 | AAAAGUGCUUACAGUGCAGGUAG | 130.76 | 120.29 | 91.17 | 123.07 |
| hsa-miR-106b-3p | MIMAT0004672 | CCGCACUGUGGGUACUUGCUGC | 42.28 | 41.93 | 101.74 | 71.50 |
| hsa-miR-106b-5p | MIMAT0000680 | UAAAGUGCUGACAGUGCAGAU | 105.69 | 78.40 | 119.00 | 137.63 |
| hsa-miR-107 | MIMAT0000104 | AGCAGCAUUGUACAGGGCUAUCA | 152.95 | 152.07 | 93.21 | 120.04 |
| hsa-miR-10a-3p | MIMAT0004555 | CAAAUUCGUAUCUAGGGGAAUA | 112.50 | 89.12 | 107.03 | 85.44 |
| hsa-miR-10a-5p | MIMAT0000253 | UACCCUGUAGAUCCGAAUUUGUG | 111.97 | 124.33 | 90.01 | 79.79 |
| hsa-miR-10b-3p | MIMAT0004556 | ACAGAUUCGAUUCUAGGGGAAU | 138.06 | 125.51 | 96.13 | 76.86 |
| hsa-miR-10b-5p | MIMAT0000254 | UACCCUGUAGAACCGAAUUUGUG | 110.05 | 109.81 | 111.83 | 81.95 |
| hsa-miR-1178-3p | MIMAT0005823 | UUGCUCACUGUUCUUCCCUAG | 116.70 | 96.58 | 101.86 | 91.31 |
| hsa-miR-1179 | MIMAT0005824 | AAGCAUUCUUUCAUUGGUUGG | 113.16 | 124.38 | 102.85 | 112.60 |

| | | | | | | |
|---|---|---|---|---|---|---|
| hsa-miR-1180 | MIMAT0005825 | UUUCCGGCUCGCGUGGGUGUGU | 95.23 | 86.33 | 78.32 | 84.34 |
| hsa-miR-1181 | MIMAT0005826 | CCGUCGCCGCCACCCGAGCCG | 122.09 | 106.78 | 105.94 | 82.51 |
| hsa-miR-1182 | MIMAT0005827 | GAGGGUCUUGGGAGGGAUGUGAC | 138.79 | 128.61 | 126.82 | 128.87 |
| hsa-miR-1183 | MIMAT0005828 | CACUGUAGGUGAUGGUGAGAGUGGGCA | 52.19 | 45.32 | 64.36 | 59.02 |
| hsa-miR-1184 | MIMAT0005829 | CCUGCAGCGACUUGAUGGCUUCC | 144.95 | 126.62 | 95.92 | 122.56 |
| hsa-miR-1184 | MIMAT0005829 | CCUGCAGCGACUUGAUGGCUUCC | 129.95 | 123.14 | 91.66 | 112.83 |
| hsa-miR-1185-5p | MIMAT0005798 | AGAGGAUACCCUUUGUAUGUU | 94.13 | 79.12 | 104.80 | 114.24 |
| hsa-miR-1193 | MIMAT0015049 | GGGAUGGUAGACCGGUGACGUGC | 118.45 | 81.94 | 95.19 | 81.56 |
| hsa-miR-1197 | MIMAT0005955 | UAGGACACAUGGUCUACUUCU | 113.14 | 88.51 | 119.20 | 99.16 |
| hsa-miR-1200 | MIMAT0005863 | CUCCUGAGCCAUUCUGAGCCUC | 28.99 | 31.74 | 145.08 | 155.96 |
| hsa-miR-1201 | MIMAT0005864 | AGCCUGAUUAAACACAUGCUCUGA | 82.47 | 111.85 | 83.60 | 78.97 |
| hsa-miR-1202 | MIMAT0005865 | GUGCCAGCUGCAGUGGGGGAG | 75.58 | 65.19 | 112.88 | 98.47 |
| hsa-miR-1203 | MIMAT0005866 | CCCGGAGCCAGGAUGCAGCUC | 135.30 | 129.60 | 134.57 | 130.80 |
| hsa-miR-1204 | MIMAT0005868 | UCGUGGCCUGGUCUCCAUUAU | 85.73 | 109.22 | 106.35 | 102.34 |
| hsa-miR-1205 | MIMAT0005869 | UCUGCAGGGUUUGCUUUGAG | 83.53 | 96.30 | 108.09 | 106.54 |
| hsa-miR-1206 | MIMAT0005870 | UGUUCAUGUAGAUGUUUAAGC | 62.56 | 77.76 | 63.54 | 58.24 |
| hsa-miR-1207-3p | MIMAT0005872 | UCAGCUGGCCCUCAUUUC | 61.40 | 60.18 | 113.17 | 95.95 |
| hsa-miR-1207-5p | MIMAT0005871 | UGGCAGGGAGGCUGGGAGGGG | 112.87 | 117.90 | 151.94 | 123.51 |
| hsa-miR-1208 | MIMAT0005873 | UCACUGUUCAGACAGGCGGA | 114.87 | 109.58 | 151.91 | 161.83 |
| hsa-miR-122-3p | MIMAT0004590 | AACGCCAUUAUCACACUAAAUA | 114.53 | 99.16 | 87.63 | 104.83 |
| hsa-miR-1224-3p | MIMAT0005459 | CCCCACCUCCUCUCUCCUCAG | 84.09 | 100.83 | 93.28 | 100.31 |
| hsa-miR-1224-5p | MIMAT0005458 | GUGAGGACUCGGGAGGUGG | 86.15 | 78.45 | 104.32 | 102.15 |
| hsa-miR-1225-3p | MIMAT0005573 | UGAGCCCCUGUGCCGCCCCCAG | 96.94 | 99.01 | 112.33 | 107.03 |
| hsa-miR-1225-5p | MIMAT0005572 | GUGGGUACGGCCCAGUGGGGGG | 131.99 | 140.53 | 102.55 | 92.98 |
| hsa-miR-122-5p | MIMAT0000421 | UGGAGUGUGACAAUGGUGUUUG | 123.66 | 141.89 | 94.10 | 115.04 |
| hsa-miR-1226-3p | MIMAT0005577 | UCACCAGCCCUGUGUUCCCUAG | 96.52 | 85.57 | 98.76 | 104.73 |
| hsa-miR-1226-5p | MIMAT0005576 | GUGAGGGCAUGCAGGCCUGGAUGGGG | 136.29 | 122.54 | 111.73 | 135.94 |
| hsa-miR-1227-3p | MIMAT0005580 | CGUGCCACCCUUUUCCCCAG | 102.71 | 126.71 | 126.08 | 149.56 |
| hsa-miR-1228-3p | MIMAT0005583 | UCACACCUGCCUCGCCCCCC | 166.12 | 150.46 | 140.76 | 137.86 |
| hsa-miR-1228-5p | MIMAT0005582 | GUGGGCGGGGGCAGGUGUGUG | 101.82 | 104.19 | 103.67 | 155.07 |
| hsa-miR-1229-3p | MIMAT0005584 | CUCUCACCACUGCCCUCCCACAG | 83.85 | 99.11 | 105.37 | 102.92 |
| hsa-miR-1231 | MIMAT0005586 | GUGUCUGGGCGGACAGCUGC | 140.49 | 134.25 | 75.53 | 78.78 |
| hsa-miR-1233-3p | MIMAT0005588 | UGAGCCCUGUCCUCCCGCAG | 66.51 | 70.96 | 99.71 | 102.70 |
| hsa-miR-1233-3p | MIMAT0005588 | UGAGCCCUGUCCUCCCGCAG | 76.74 | 64.70 | 109.18 | 101.08 |
| hsa-miR-1234-3p | MIMAT0005589 | UCGGCCUGACCACCCACCCCAC | 101.30 | 96.12 | 113.06 | 121.44 |
| hsa-miR-1236-3p | MIMAT0005591 | CCUCUUCCCCUUGUCUCUCCAG | 134.26 | 123.35 | 122.18 | 115.82 |
| hsa-miR-1237-3p | MIMAT0005592 | UCCUUCUGCUCCGUCCCCCAG | 63.63 | 72.23 | 98.39 | 128.44 |
| hsa-miR-1238-3p | MIMAT0005593 | CUUCCUCGUCUGUCUGCCCC | 107.18 | 97.60 | 85.01 | 35.11 |
| hsa-miR-1243 | MIMAT0005894 | AACUGGAUCAAUUAUAGGAGUG | 119.29 | 108.32 | 56.96 | 22.21 |
| hsa-miR-124-3p | MIMAT0000422 | UAAGGCACGCGGUGAAUGCC | 85.48 | 81.06 | 100.65 | 109.74 |
| hsa-miR-1244 | MIMAT0005896 | AAGUAGUUGGUUUGUAUGAGAUGGUU | 99.83 | 80.53 | 85.69 | 101.62 |

| | | | | | | |
|---|---|---|---|---|---|---|
| hsa-miR-1244 | MIMAT0005896 | AAGUAGUUGGUUUGUAUGAGAUGGUU | 73.40 | 91.43 | 91.12 | 106.18 |
| hsa-miR-1245a | MIMAT0005897 | AAGUGAUCUAAAGGCCUACAU | 141.91 | 145.35 | 113.00 | 129.59 |
| hsa-miR-124-5p | MIMAT0004591 | CGUGUUCACAGCGGACCUUGAU | 39.80 | 37.58 | 93.96 | 99.34 |
| hsa-miR-1246 | MIMAT0005898 | AAUGGAUUUUUGGAGCAGG | 86.87 | 106.17 | 103.77 | 110.50 |
| hsa-miR-1247-5p | MIMAT0005899 | ACCCGUCCCGUUCGUCCCCGGA | 95.76 | 130.95 | 105.47 | 87.89 |
| hsa-miR-1248 | MIMAT0005900 | ACCUUCUUGUAUAAGCACUGUGCUAAA | 110.60 | 90.56 | 114.00 | 80.92 |
| hsa-miR-1249 | MIMAT0005901 | ACGCCCUUCCCCCCCUUCUUCA | 80.02 | 96.24 | 93.06 | 108.84 |
| hsa-miR-1250 | MIMAT0005902 | ACGGUGCUGGAUGUGGCCUUU | 71.49 | 61.09 | 110.40 | 94.68 |
| hsa-miR-1251 | MIMAT0005903 | ACUCUAGCUGCCAAAGGCGCU | 121.16 | 116.00 | 107.65 | 106.62 |
| hsa-miR-1252 | MIMAT0005944 | AGAAGGAAAUUGAAUUCAUUUA | 126.57 | 104.36 | 127.72 | 113.14 |
| hsa-miR-1253 | MIMAT0005904 | AGAGAAGAAGAUCAGCCUGCA | 102.13 | 80.01 | 72.53 | 84.02 |
| hsa-miR-1254 | MIMAT0005905 | AGCCUGGAAGCUGGAGCCUGCAGU | 49.17 | 59.73 | 83.65 | 99.30 |
| hsa-miR-1255a | MIMAT0005906 | AGGAUGAGCAAAGAAAGUAGAUU | 88.54 | 80.66 | 107.65 | 118.68 |
| hsa-miR-1255b-5p | MIMAT0005945 | CGGAUGAGCAAAGAAAGUGGUU | 74.20 | 67.39 | 109.32 | 98.34 |
| hsa-miR-1256 | MIMAT0005907 | AGGCAUUGACUUCUCACUAGCU | 38.12 | 42.15 | 72.95 | 79.25 |
| hsa-miR-1257 | MIMAT0005908 | AGUGAAUGAUGGGUUCUGACC | 197.57 | 160.70 | 135.85 | 123.64 |
| hsa-miR-1258 | MIMAT0005909 | AGUUAGGAUUAGGUCGUGGAA | 179.87 | 232.76 | 108.22 | 116.78 |
| hsa-miR-1259 | MIMAT0005910 | AUAUAUGAUGACUUAGCUUUU | 76.42 | 94.49 | 83.16 | 92.03 |
| hsa-miR-125a-3p | MIMAT0004602 | ACAGGUGAGGUUCUUGGGAGCC | 66.15 | 67.12 | 104.32 | 117.84 |
| hsa-miR-125a-5p | MIMAT0000443 | UCCCUGAGACCCUUUAACCUGUGA | 88.45 | 101.98 | 97.01 | 95.28 |
| hsa-miR-125b-1-3p | MIMAT0004592 | ACGGGUUAGGCUCUUGGGAGCU | 110.90 | 129.83 | 87.71 | 102.82 |
| hsa-miR-125b-2-3p | MIMAT0004603 | UCACAAGUCAGGCUCUUGGGAC | 65.00 | 51.04 | 62.15 | 88.87 |
| hsa-miR-125b-5p | MIMAT0000423 | UCCCUGAGACCCUAACUUGUGA | 62.52 | 83.48 | 73.26 | 115.05 |
| hsa-miR-1260a | MIMAT0005911 | AUCCCACCUCUGCCACCA | 171.48 | 169.77 | 102.52 | 111.07 |
| hsa-miR-1260b | MIMAT0015041 | AUCCCACCACUGCCACCAU | 93.11 | 103.66 | 115.34 | 121.75 |
| hsa-miR-1261 | MIMAT0005913 | AUGGAUAAGGCUUUGGCUU | 76.42 | 92.95 | 100.24 | 102.82 |
| hsa-miR-1262 | MIMAT0005914 | AUGGGUGAAUUUGUAGAAGGAU | 66.49 | 63.99 | 96.22 | 79.39 |
| hsa-miR-1263 | MIMAT0005915 | AUGGUACCCUGGCAUACUGAGU | 102.52 | 103.71 | 129.86 | 137.72 |
| hsa-miR-126-3p | MIMAT0000445 | UCGUACCGUGAGUAAUAAUGCG | 112.30 | 124.65 | 118.57 | 101.17 |
| hsa-miR-1264 | MIMAT0005791 | CAAGUCUUAUUUGAGCACCUGUU | 138.86 | 129.83 | 59.80 | 52.68 |
| hsa-miR-1265 | MIMAT0005918 | CAGGAUGUGGUCAAGUGUUGUU | 134.20 | 146.73 | 105.37 | 107.89 |
| hsa-miR-126-5p | MIMAT0000444 | CAUUAUUACUUUUGGUACGCG | 113.16 | 90.56 | 103.41 | 78.35 |
| hsa-miR-1266 | MIMAT0005920 | CCUCAGGGCUGUAGAACAGGGCU | 134.15 | 118.21 | 95.23 | 85.17 |
| hsa-miR-1267 | MIMAT0005921 | CCUGUUGAAGUGUAAUCCCCA | 110.76 | 101.11 | 107.61 | 108.03 |
| hsa-miR-1268a | MIMAT0005922 | CGGGCGUGGUGGUGGGGG | 94.84 | 100.76 | 73.66 | 78.84 |
| hsa-miR-1269a | MIMAT0005923 | CUGGACUGAGCCGUGCUACUGG | 59.36 | 21.49 | 80.55 | 48.38 |
| hsa-miR-1270 | MIMAT0005924 | CUGGAGAUAUGGAAGAGCUGUGU | 125.32 | 138.58 | 88.91 | 81.83 |
| hsa-miR-1270 | MIMAT0005924 | CUGGAGAUAUGGAAGAGCUGUGU | 118.58 | 138.58 | 81.45 | 81.40 |
| hsa-miR-1271-5p | MIMAT0005796 | CUUGGCACCUAGCAAGCACUCA | 99.97 | 110.28 | 82.76 | 115.02 |
| hsa-miR-1272 | MIMAT0005925 | GAUGAUGAUGGCAGCAAAUUCUGAAA | 141.70 | 139.97 | 163.23 | 134.14 |
| hsa-miR-1273a | MIMAT0005926 | GGGCGACAAAGCAAGACUCUUUCUU | 135.71 | 143.44 | 83.52 | 110.63 |

| | | | | | | |
|---|---|---|---|---|---|---|
| hsa-miR-1273c | MIMAT0015017 | GGCGACAAAACGAGACCCUGUC | 130.63 | 118.27 | 106.82 | 117.69 |
| hsa-miR-1273d | MIMAT0015090 | GAACCCAUGAGGUUGAGGCUGCAGU | 97.89 | 94.62 | 97.34 | 86.90 |
| hsa-miR-1273e | MIMAT0018079 | UUGCUUGAACCCAGGAAGUGGA | 70.09 | 81.33 | 79.73 | 77.49 |
| hsa-miR-127-3p | MIMAT0000446 | UCGGAUCCGUCUGAGCUUGGCU | 67.87 | 73.75 | 68.92 | 79.61 |
| hsa-miR-1274a | MIMAT0005927 | GUCCCUGUUCAGGCGCCA | 89.33 | 96.24 | 94.77 | 111.23 |
| hsa-miR-1274b | MIMAT0005938 | UCCCUGUUCGGGCGCCA | 109.93 | 126.62 | 111.55 | 104.07 |
| hsa-miR-1275 | MIMAT0005929 | GUGGGGGAGAGGCUGUC | 25.09 | 44.86 | 79.58 | 120.32 |
| hsa-miR-127-5p | MIMAT0004604 | CUGAAGCUCAGAGGGCUCUGAU | 88.18 | 130.82 | 70.19 | 70.88 |
| hsa-miR-1276 | MIMAT0005930 | UAAAGAGCCCUGUGGAGACA | 159.21 | 175.34 | 102.31 | 120.83 |
| hsa-miR-1277-3p | MIMAT0005933 | UACGUAGAUAUAUAUGUAUUUU | 112.99 | 115.43 | 106.71 | 100.13 |
| hsa-miR-1278 | MIMAT0005936 | UAGUACUGUGCAUAUCAUCUAU | 98.64 | 97.59 | 110.61 | 119.82 |
| hsa-miR-1279 | MIMAT0005937 | UCAUAUUGCUUCUUUCU | 169.64 | 159.80 | 117.45 | 112.66 |
| hsa-miR-128 | MIMAT0000424 | UCACAGUGAACCGGUCUCUUU | 119.68 | 102.74 | 94.52 | 90.63 |
| hsa-miR-1280 | MIMAT0005946 | UCCCACCGCUGCCACCC | 88.40 | 80.08 | 108.55 | 113.85 |
| hsa-miR-1281 | MIMAT0005939 | UCGCCUCCUCCUCUCCC | 90.29 | 83.83 | 107.03 | 121.36 |
| hsa-miR-1282 | MIMAT0005940 | UCGUUUGCCUUUUUCUGCUU | 118.45 | 134.23 | 110.53 | 99.21 |
| hsa-miR-1283 | MIMAT0005799 | UCUACAAAGGAAAGCGCUUUCU | 140.73 | 129.83 | 103.66 | 114.24 |
| hsa-miR-1284 | MIMAT0005941 | UCUAUACAGACCCUGGCUUUUC | 109.30 | 90.73 | 101.22 | 99.81 |
| hsa-miR-1285-3p | MIMAT0005876 | UCUGGGCAACAAAGUGAGACCU | 89.47 | 97.56 | 80.31 | 101.55 |
| hsa-miR-1286 | MIMAT0005877 | UGCAGGACCAAGAUGAGCCCU | 88.16 | 82.27 | 88.36 | 106.93 |
| hsa-miR-1287 | MIMAT0005878 | UGCUGGAUCAGUGGUUCGAGUC | 100.35 | 99.72 | 100.06 | 123.73 |
| hsa-miR-1288 | MIMAT0005942 | UGGACUGCCCUGAUCUGGAGA | 125.82 | 105.24 | 84.86 | 78.06 |
| hsa-miR-1289 | MIMAT0005879 | UGGAGUCCAGGAAUCUGCAUUUU | 102.96 | 79.43 | 92.91 | 81.85 |
| hsa-miR-1290 | MIMAT0005880 | UGGAUUUUUGGAUCAGGGA | 129.92 | 136.08 | 100.39 | 110.04 |
| hsa-miR-1291 | MIMAT0005881 | UGGCCCUGACUGAAGACCAGCAGU | 125.63 | 98.77 | 126.67 | 110.42 |
| hsa-miR-129-1-3p | MIMAT0004548 | AAGCCCUUACCCCAAAAAGUAU | 147.09 | 133.17 | 127.98 | 123.87 |
| hsa-miR-129-2-3p | MIMAT0004605 | AAGCCCUUACCCCAAAAAGCAU | 130.20 | 145.87 | 116.12 | 118.43 |
| hsa-miR-1292-5p | MIMAT0005943 | UGGGAACGGGUUCCGGCAGACGCUG | 114.12 | 114.84 | 156.17 | 136.96 |
| hsa-miR-1293 | MIMAT0005883 | UGGGUGGUCUGGAGAUUUGUGC | 42.19 | 34.36 | 119.24 | 138.84 |
| hsa-miR-1294 | MIMAT0005884 | UGUGAGGUUGGCAUUGUUGUCU | 36.74 | 33.06 | 62.19 | 72.23 |
| hsa-miR-1295a | MIMAT0005885 | UUAGGCCGCAGAUCUGGGUGA | 116.97 | 105.37 | 108.17 | 91.69 |
| hsa-miR-129-5p | MIMAT0000242 | CUUUUUGCGGUCUGGGCUUGC | 127.86 | 101.94 | 107.82 | 115.57 |
| hsa-miR-1296 | MIMAT0005794 | UUAGGGCCCUGGCUCCAUCUCC | 135.30 | 130.50 | 99.05 | 87.84 |
| hsa-miR-1297 | MIMAT0005886 | UUCAAGUAAUUCAGGUG | 67.92 | 63.87 | 80.62 | 83.49 |
| hsa-miR-1298 | MIMAT0005800 | UUCAUUCGGCUGUCCAGAUGUA | 131.10 | 120.16 | 115.11 | 128.44 |
| hsa-miR-1299 | MIMAT0005887 | UUCUGGAAUUCUGUGUGAGGGA | 177.44 | 151.54 | 96.24 | 102.34 |
| hsa-miR-1300 | MIMAT0005888 | UUGAGAAGGAGGCUGCUG | 130.16 | 110.81 | 72.64 | 78.04 |
| hsa-miR-1301 | MIMAT0005797 | UUGCAGCUGCCUGGGAGUGACUUC | 99.24 | 97.89 | 106.40 | 82.57 |
| hsa-miR-1302 | MIMAT0005890 | UUGGGACAUACUUAUGCUAAA | 98.92 | 120.50 | 76.87 | 108.47 |
| hsa-miR-1302 | MIMAT0005890 | UUGGGACAUACUUAUGCUAAA | 109.87 | 107.42 | 94.54 | 110.95 |
| hsa-miR-1303 | MIMAT0005891 | UUUAGAGACGGGGUCUUGCUCU | 130.73 | 124.03 | 112.08 | 128.49 |

| | | | | | | |
|---|---|---|---|---|---|---|
| hsa-miR-1304-5p | MIMAT0005892 | UUUGAGGCUACAGUGAGAUGUG | 67.90 | 64.48 | 93.93 | 95.04 |
| hsa-miR-1305 | MIMAT0005893 | UUUUCAACUCUAAUGGGAGAGA | 127.16 | 113.37 | 84.08 | 106.46 |
| hsa-miR-1306-3p | MIMAT0005950 | ACGUUGGCUCUGGUGGUG | 84.02 | 78.02 | 110.74 | 122.94 |
| hsa-miR-1307-3p | MIMAT0005951 | ACUCGGCGUGGCGUCGGUCGUG | 84.09 | 98.50 | 120.45 | 87.36 |
| hsa-miR-1308 | MIMAT0005947 | GCAUGGGUGGUUCAGUGG | 122.50 | 119.68 | 100.34 | 112.19 |
| hsa-miR-130a-3p | MIMAT0000425 | CAGUGCAAUGUUAAAAGGGCAU | 175.16 | 213.42 | 115.12 | 111.32 |
| hsa-miR-130a-5p | MIMAT0004593 | UUCACAUUGUGCUACUGUCUGC | 101.58 | 74.52 | 89.42 | 86.31 |
| hsa-miR-130b-3p | MIMAT0000691 | CAGUGCAAUGAUGAAAGGGCAU | 121.07 | 141.58 | 120.69 | 86.88 |
| hsa-miR-130b-5p | MIMAT0004680 | ACUCUUUCCCUGUUGCACUAC | 115.73 | 115.22 | 51.01 | 66.79 |
| hsa-miR-1321 | MIMAT0005952 | CAGGGAGGUGAAUGUGAU | 51.50 | 68.29 | 86.22 | 89.75 |
| hsa-miR-1322 | MIMAT0005953 | GAUGAUGCUGCUGAUGCUG | 81.08 | 76.05 | 88.85 | 76.16 |
| hsa-miR-1323 | MIMAT0005795 | UCAAAACUGAGGGGCAUUUUCU | 79.22 | 66.83 | 113.34 | 125.66 |
| hsa-miR-132-3p | MIMAT0000426 | UAACAGUCUACAGCCAUGGUCG | 138.93 | 154.68 | 84.13 | 97.21 |
| hsa-miR-1324 | MIMAT0005956 | CCAGACAGAAUUCUAUGCACUUUC | 120.17 | 134.11 | 117.88 | 115.53 |
| hsa-miR-132-5p | MIMAT0004594 | ACCGUGGCUUUCGAUUGUUACU | 75.49 | 89.88 | 105.94 | 116.14 |
| hsa-miR-133a | MIMAT0000427 | UUUGGUCCCCUUCAACCAGCUG | 99.46 | 112.98 | 79.75 | 69.91 |
| hsa-miR-133b | MIMAT0000770 | UUUGGUCCCCUUCAACCAGCUA | 142.29 | 185.05 | 118.31 | 104.55 |
| hsa-miR-134 | MIMAT0000447 | UGUGACUGGUUGACCAGAGGGG | 91.39 | 97.33 | 102.40 | 93.98 |
| hsa-miR-135a-3p | MIMAT0004595 | UAUAGGGAUUGGAGCCGUGGCG | 124.92 | 128.04 | 96.87 | 84.85 |
| hsa-miR-135a-5p | MIMAT0000428 | UAUGGCUUUUUAUUCCUAUGUGA | 84.81 | 71.44 | 46.70 | 38.71 |
| hsa-miR-135b-3p | MIMAT0004698 | AUGUAGGGCUAAAAGCCAUGGG | 102.13 | 103.45 | 96.49 | 112.19 |
| hsa-miR-135b-5p | MIMAT0000758 | UAUGGCUUUUCAUUCCUAUGUGA | 69.21 | 80.66 | 90.85 | 77.25 |
| hsa-miR-136-3p | MIMAT0004606 | CAUCAUCGUCUCAAAUGAGUCU | 109.04 | 97.56 | 74.61 | 62.83 |
| hsa-miR-136-5p | MIMAT0000448 | ACUCCAUUUGUUUUGAUGAUGGA | 109.99 | 122.90 | 97.74 | 97.25 |
| hsa-miR-137 | MIMAT0000429 | UUAUUGCUUAAGAAUACGCGUAG | 116.59 | 99.52 | 80.64 | 64.36 |
| hsa-miR-138-1-3p | MIMAT0004607 | GCUACUUCACAACACCAGGGCC | 95.41 | 99.17 | 111.75 | 103.93 |
| hsa-miR-138-2-3p | MIMAT0004596 | GCUAUUUCACGACACCAGGGUU | 100.39 | 84.97 | 93.49 | 108.37 |
| hsa-miR-138-5p | MIMAT0000430 | AGCUGGUGUUGUGAAUCAGGCCG | 75.53 | 66.62 | 76.18 | 88.27 |
| hsa-miR-139-3p | MIMAT0004552 | GGAGACGCGGCCCUGUUGGAGU | 78.33 | 86.48 | 124.22 | 117.39 |
| hsa-miR-139-5p | MIMAT0000250 | UCUACAGUGCACGUGUCUCCAG | 68.75 | 29.95 | 77.76 | 46.24 |
| hsa-miR-140-3p | MIMAT0004597 | UACCACAGGGUAGAACCACGG | 161.52 | 168.23 | 90.07 | 112.61 |
| hsa-miR-140-5p | MIMAT0000431 | CAGUGGUUUUACCCUAUGGUAG | 106.14 | 122.03 | 93.72 | 79.92 |
| hsa-miR-141-3p | MIMAT0000432 | UAACACUGUCUGGUAAAGAUGG | 96.55 | 97.06 | 99.61 | 114.49 |
| hsa-miR-141-5p | MIMAT0004598 | CAUCUUCCAGUACAGUGUUGGA | 87.00 | 86.32 | 60.12 | 57.76 |
| hsa-miR-142-3p | MIMAT0000434 | UGUAGUGUUUCCUACUUUAUGGA | 135.31 | 126.32 | 88.46 | 84.83 |
| hsa-miR-142-5p | MIMAT0000433 | CAUAAAGUAGAAAGCACUACU | 114.35 | 119.68 | 93.92 | 83.06 |
| hsa-miR-143-3p | MIMAT0000435 | UGAGAUGAAGCACUGUAGCUC | 97.71 | 121.47 | 83.17 | 103.36 |
| hsa-miR-143-5p | MIMAT0004599 | GGUGCAGUGCUGCAUCUCUGGU | 123.05 | 106.37 | 80.10 | 89.82 |
| hsa-miR-144-3p | MIMAT0000436 | UACAGUAUAGAUGAUGUACU | 161.18 | 167.61 | 82.85 | 83.90 |
| hsa-miR-144-5p | MIMAT0004600 | GGAUAUCAUCAUAUACUGUAAG | 112.14 | 121.62 | 130.25 | 112.25 |
| hsa-miR-145-3p | MIMAT0004601 | GGAUUCCUGGAAAUACUGUUCU | 53.83 | 42.58 | 110.06 | 106.57 |

| | | | | | | |
|---|---|---|---|---|---|---|
| hsa-miR-145-5p | MIMAT0000437 | GUCCAGUUUUCCCAGGAAUCCCU | 112.60 | 116.38 | 121.53 | 135.30 |
| hsa-miR-1468 | MIMAT0006789 | CUCCGUUUGCCUGUUUCGCUG | 73.03 | 65.59 | 84.08 | 100.25 |
| hsa-miR-1469 | MIMAT0007347 | CUCGGCGCGGGGCGCGGGCUCC | 93.26 | 101.48 | 104.82 | 103.18 |
| hsa-miR-146a-3p | MIMAT0004608 | CCUCUGAAAUUCAGUUCUUCAG | 116.68 | 102.55 | 71.67 | 87.84 |
| hsa-miR-146a-5p | MIMAT0000449 | UGAGAACUGAAUUCCAUGGGUU | 149.05 | 142.62 | 98.29 | 118.51 |
| hsa-miR-146b-3p | MIMAT0004766 | UGCCCUGUGGACUCAGUUCUGG | 103.50 | 113.79 | 125.67 | 112.21 |
| hsa-miR-146b-5p | MIMAT0002809 | UGAGAACUGAAUUCCAUAGGCU | 107.18 | 92.81 | 77.26 | 94.72 |
| hsa-miR-1470 | MIMAT0007348 | GCCCUCCGCCCGUGCACCCCG | 71.76 | 76.05 | 117.90 | 123.76 |
| hsa-miR-1471 | MIMAT0007349 | GCCCGCGUGUGGAGCCAGGUGU | 81.18 | 89.48 | 81.94 | 83.54 |
| hsa-miR-147a | MIMAT0000251 | GUGUGUGGAAAUGCUUCUGC | 116.76 | 96.30 | 102.04 | 96.89 |
| hsa-miR-147b | MIMAT0004928 | GUGUGCGGAAAUGCUUCUGCUA | 41.11 | 57.15 | 107.15 | 104.58 |
| hsa-miR-148a-3p | MIMAT0000243 | UCAGUGCACUACAGAACUUUGU | 109.32 | 122.35 | 94.43 | 103.16 |
| hsa-miR-148a-5p | MIMAT0004549 | AAAGUUCUGAGACACUCCGACU | 109.66 | 84.33 | 75.58 | 80.63 |
| hsa-miR-148b-3p | MIMAT0000759 | UCAGUGCAUCACAGAACUUUGU | 123.55 | 90.26 | 125.26 | 126.32 |
| hsa-miR-148b-5p | MIMAT0004699 | AAGUUCUGUUAUACACUCAGGC | 89.47 | 96.02 | 63.22 | 53.95 |
| hsa-miR-149-3p | MIMAT0004609 | AGGGAGGGACGGGGGCUGUGC | 88.79 | 126.88 | 95.20 | 79.10 |
| hsa-miR-149-5p | MIMAT0000450 | UCUGGCUCCGUGUCUUCACUCCC | 97.27 | 103.45 | 98.92 | 105.58 |
| hsa-miR-150-3p | MIMAT0004610 | CUGGUACAGGCCUGGGGGACAG | 133.80 | 127.73 | 126.67 | 113.99 |
| hsa-miR-150-5p | MIMAT0000451 | UCUCCCAACCCUUGUACCAGUG | 97.47 | 106.53 | 104.04 | 95.51 |
| hsa-miR-151a-3p | MIMAT0000757 | CUAGACUGAAGCUCCUUGAGG | 85.74 | 63.76 | 113.91 | 123.13 |
| hsa-miR-151a-5p | MIMAT0004697 | UCGAGGAGCUCACAGUCUAGU | 153.49 | 146.50 | 133.77 | 120.32 |
| hsa-miR-152 | MIMAT0000438 | UCAGUGCAUGACAGAACUUGG | 177.78 | 149.43 | 82.79 | 108.37 |
| hsa-miR-153 | MIMAT0000439 | UUGCAUAGUCACAAAAGUGAUC | 57.63 | 39.04 | 104.80 | 118.78 |
| hsa-miR-1537 | MIMAT0007399 | AAAACCGUCUAGUUACAGUUGU | 135.14 | 123.68 | 140.11 | 145.34 |
| hsa-miR-1538 | MIMAT0007400 | CGGCCCGGGCUGCUGCUGUUCCU | 90.29 | 75.90 | 112.08 | 111.85 |
| hsa-miR-1539 | MIMAT0007401 | UCCUGCGCGUCCCAGAUGCCC | 194.06 | 154.64 | 114.88 | 115.90 |
| hsa-miR-154-3p | MIMAT0000453 | AAUCAUACACGGUUGACCUAUU | 113.95 | 74.32 | 84.36 | 87.13 |
| hsa-miR-154-5p | MIMAT0000452 | UAGGUUAUCCGUGUUGCCUUCG | 84.17 | 100.90 | 89.06 | 87.46 |
| hsa-miR-155-3p | MIMAT0004658 | CUCCUACAUAUUAGCAUUAACA | 128.61 | 139.04 | 104.23 | 111.07 |
| hsa-miR-155-5p | MIMAT0000646 | UUAAUGCUAAUCGUGAUAGGGGU | 148.36 | 130.95 | 84.38 | 115.16 |
| hsa-miR-15a-3p | MIMAT0004488 | CAGGCCAUAUUGUGCUGCCUCA | 110.85 | 116.18 | 77.92 | 63.41 |
| hsa-miR-15a-5p | MIMAT0000068 | UAGCAGCACAUAAUGGUUUGUG | 104.58 | 85.91 | 73.92 | 87.59 |
| hsa-miR-15b-3p | MIMAT0004586 | CGAAUCAUUAUUUGCUGCUCUA | 177.07 | 165.16 | 129.29 | 135.82 |
| hsa-miR-15b-5p | MIMAT0000417 | UAGCAGCACAUCAUGGUUUACA | 69.16 | 88.37 | 86.77 | 82.99 |
| hsa-miR-16-1-3p | MIMAT0004489 | CCAGUAUUAACUGUGCUGCUGA | 116.50 | 89.88 | 101.95 | 93.30 |
| hsa-miR-16-2-3p | MIMAT0004518 | CCAAUAUUACUGUGCUGCUUUA | 80.60 | 67.44 | 85.79 | 76.46 |
| hsa-miR-16-5p | MIMAT0000069 | UAGCAGCACGUAAAUAUUGGCG | 68.47 | 70.77 | 75.07 | 90.15 |
| hsa-miR-17-3p | MIMAT0000071 | ACUGCAGUGAAGGCACUUGUAG | 73.85 | 80.73 | 109.66 | 103.18 |
| hsa-miR-17-5p | MIMAT0000070 | CAAAGUGCUUACAGUGCAGGUAG | 95.94 | 87.30 | 85.46 | 102.44 |
| hsa-miR-181a-2-3p | MIMAT0004558 | ACCACUGACCGUUGACUGUACC | 176.77 | 212.80 | 151.87 | 143.32 |
| hsa-miR-181a-3p | MIMAT0000270 | ACCAUCGACCGUUGAUUGUACC | 68.79 | 74.76 | 98.54 | 105.59 |

| | | | | | | |
|---|---|---|---|---|---|---|
| hsa-miR-181a-5p | MIMAT0000256 | AACAUUCAACGCUGUCGGUGAGU | 114.26 | 113.40 | 82.66 | 84.47 |
| hsa-miR-181b-5p | MIMAT0000257 | AACAUUCAUUGCUGUCGGUGGGU | 119.01 | 94.44 | 108.04 | 89.75 |
| hsa-miR-181c-3p | MIMAT0004559 | AACCAUCGACCGUUGAGUGGAC | 124.58 | 106.81 | 99.90 | 95.48 |
| hsa-miR-181c-5p | MIMAT0000258 | AACAUUCAACCUGUCGGUGAGU | 95.95 | 91.75 | 74.66 | 85.13 |
| hsa-miR-181d | MIMAT0002821 | AACAUUCAUUGUUGUCGGUGGGU | 173.03 | 156.59 | 83.97 | 92.51 |
| hsa-miR-182-3p | MIMAT0000260 | UGGUUCUAGACUUGCCAACUA | 75.49 | 76.05 | 82.02 | 93.93 |
| hsa-miR-1825 | MIMAT0006765 | UCCAGUGCCCUCCUCUCC | 174.60 | 160.38 | 94.82 | 89.07 |
| hsa-miR-182-5p | MIMAT0000259 | UUUGGCAAUGGUAGAACUCACACU | 92.30 | 106.34 | 117.84 | 108.36 |
| hsa-miR-1826 | MIMAT0006766 | AUUGAUCAUCGACACUUCGAACGCAAU | 93.40 | 102.55 | 124.30 | 111.23 |
| hsa-miR-1827 | MIMAT0006767 | UGAGGCAGUAGAUUGAAU | 76.05 | 78.02 | 92.87 | 102.87 |
| hsa-miR-183-3p | MIMAT0004560 | GUGAAUUACCGAAGGGCCAUAA | 116.30 | 86.44 | 103.67 | 74.59 |
| hsa-miR-183-5p | MIMAT0000261 | UAUGGCACUGGUAGAAUUCACU | 106.08 | 90.86 | 112.04 | 93.19 |
| hsa-miR-184 | MIMAT0000454 | UGGACGGAGAACUGAUAAGGGU | 85.71 | 72.95 | 110.41 | 102.36 |
| hsa-miR-185-3p | MIMAT0004611 | AGGGGCUGGCUUUCCUCUGGUC | 43.80 | 45.32 | 87.14 | 101.55 |
| hsa-miR-185-5p | MIMAT0000455 | UGGAGAGAAAGGCAGUUCCUGA | 143.83 | 131.44 | 94.56 | 92.57 |
| hsa-miR-186-3p | MIMAT0004612 | GCCCAAAGGUGAAUUUUUUGGG | 140.49 | 130.73 | 103.96 | 120.30 |
| hsa-miR-186-5p | MIMAT0000456 | CAAAGAAUUCUCCUUUUGGGCU | 128.58 | 107.84 | 115.34 | 86.50 |
| hsa-miR-187-3p | MIMAT0000262 | UCGUGUCUUGUGUUGCAGCCGG | 96.93 | 90.32 | 79.00 | 74.12 |
| hsa-miR-187-5p | MIMAT0004561 | GGCUACAACACAGGACCCGGGC | 91.43 | 71.14 | 114.11 | 118.98 |
| hsa-miR-188-3p | MIMAT0004613 | CUCCCACAUGCAGGGUUUGCA | 69.78 | 74.84 | 72.98 | 69.06 |
| hsa-miR-188-5p | MIMAT0000457 | CAUCCCUUGCAUGGUGGAGGG | 130.25 | 129.52 | 83.63 | 79.40 |
| hsa-miR-18a-3p | MIMAT0002891 | ACUGCCCUAAGUGCUCCUUCUGG | 79.51 | 94.85 | 101.41 | 110.02 |
| hsa-miR-18a-5p | MIMAT0000072 | UAAGGUGCAUCUAGUGCAGAUAG | 100.26 | 126.11 | 142.80 | 163.49 |
| hsa-miR-18b-3p | MIMAT0004751 | UGCCCUAAAUGCCCCUUCUGGC | 90.40 | 116.00 | 129.86 | 127.57 |
| hsa-miR-18b-5p | MIMAT0001412 | UAAGGUGCAUCUAGUGCAGUUAG | 94.45 | 108.74 | 122.87 | 106.26 |
| hsa-miR-1908 | MIMAT0007881 | CGGCGGGGACGGCGAUUGGUC | 78.33 | 61.62 | 125.91 | 124.79 |
| hsa-miR-1909-3p | MIMAT0007883 | CGCAGGGGCCGGGUGCUCACCG | 82.01 | 82.20 | 115.62 | 116.14 |
| hsa-miR-1909-5p | MIMAT0007882 | UGAGUGCCGGUGCCUGCCCUG | 143.40 | 132.64 | 110.58 | 92.84 |
| hsa-miR-190a | MIMAT0000458 | UGAUAUGUUUGAUAUAUUAGGU | 66.13 | 75.39 | 85.77 | 90.24 |
| hsa-miR-190b | MIMAT0004929 | UGAUAUGUUUGAUAUUGGGUU | 78.15 | 63.78 | 97.83 | 98.47 |
| hsa-miR-1910 | MIMAT0007884 | CCAGUCCUGUGCCUGCCGCCU | 138.86 | 123.68 | 109.92 | 110.43 |
| hsa-miR-1911-3p | MIMAT0007886 | CACCAGGCAUUGUGGUCUCC | 84.13 | 92.67 | 85.01 | 87.34 |
| hsa-miR-1911-5p | MIMAT0007885 | UGAGUACCGCCAUGUCUGUUGGG | 111.36 | 127.21 | 94.56 | 97.06 |
| hsa-miR-1912 | MIMAT0007887 | UACCCAGAGCAUGCAGUGUGAA | 71.29 | 81.37 | 91.78 | 99.77 |
| hsa-miR-1913 | MIMAT0007888 | UCUGCCCCCUCCGCUGCUGCCA | 88.32 | 105.52 | 120.42 | 104.02 |
| hsa-miR-191-3p | MIMAT0001618 | GCUGCGCUUGGAUUUCGUCCCC | 122.77 | 149.72 | 152.22 | 151.35 |
| hsa-miR-1914-3p | MIMAT0007890 | GGAGGGGUCCCGCACUGGGAGG | 110.90 | 114.46 | 120.75 | 121.22 |
| hsa-miR-1914-5p | MIMAT0007889 | CCCUGUGCCCGGCCCACUUCUG | 125.63 | 126.14 | 106.16 | 110.82 |
| hsa-miR-1915-3p | MIMAT0007892 | CCCCAGGGCGACGCGGCGGG | 128.53 | 108.52 | 102.97 | 97.08 |
| hsa-miR-1915-5p | MIMAT0007891 | ACCUUGCCUUGCUGCCCGGGCC | 81.37 | 77.55 | 95.96 | 67.87 |
| hsa-miR-191-5p | MIMAT0000440 | CAACGGAAUCCCAAAAGCAGCUG | 94.13 | 84.13 | 131.97 | 96.76 |

| | | | | | | |
|---|---|---|---|---|---|---|
| hsa-miR-192-3p | MIMAT0004543 | CUGCCAAUUCCAUAGGUCACAG | 131.07 | 142.51 | 103.92 | 89.56 |
| hsa-miR-192-5p | MIMAT0000222 | CUGACCUAUGAAUUGACAGCC | 93.88 | 102.09 | 91.85 | 86.35 |
| hsa-miR-193a-3p | MIMAT0000459 | AACUGGCCUACAAAGUCCCAGU | 102.91 | 90.56 | 112.33 | 72.78 |
| hsa-miR-193a-5p | MIMAT0004614 | UGGGUCUUUGCGGGCGAGAUGA | 114.60 | 115.93 | 106.51 | 105.74 |
| hsa-miR-193b-3p | MIMAT0002819 | AACUGGCCCUCAAAGUCCCGCU | 81.18 | 77.31 | 115.74 | 85.93 |
| hsa-miR-193b-5p | MIMAT0004767 | CGGGGUUUUGAGGGCGAGAUGA | 166.57 | 199.38 | 141.69 | 145.03 |
| hsa-miR-194-3p | MIMAT0004671 | CCAGUGGGGCUGCUGUUAUCUG | 97.86 | 91.42 | 110.49 | 97.74 |
| hsa-miR-194-5p | MIMAT0000460 | UGUAACAGCAACUCCAUGUGGA | 172.95 | 174.42 | 60.48 | 78.78 |
| hsa-miR-195-3p | MIMAT0004615 | CCAAUAUUGGCUGUGCUGCUCC | 120.57 | 142.81 | 102.35 | 105.16 |
| hsa-miR-195-5p | MIMAT0000461 | UAGCAGCACAGAAAUAUUGGC | 59.59 | 92.58 | 78.07 | 75.30 |
| hsa-miR-196a-3p | MIMAT0004562 | CGGCAACAAGAAACUGCCUGAG | 114.75 | 108.52 | 122.90 | 120.98 |
| hsa-miR-196a-5p | MIMAT0000226 | UAGGUAGUUUCAUGUUGUUGGG | 92.57 | 92.83 | 78.04 | 68.80 |
| hsa-miR-196b-3p | MIMAT0009201 | UCGACAGCACGACACUGCCUUC | 115.52 | 104.36 | 124.30 | 91.66 |
| hsa-miR-196b-5p | MIMAT0001080 | UAGGUAGUUUCCUGUUGUUGGG | 89.02 | 74.44 | 72.24 | 54.49 |
| hsa-miR-1972 | MIMAT0009447 | UCAGGCCAGGCACAGUGGCUCA | 134.19 | 146.58 | 109.52 | 117.99 |
| hsa-miR-1972 | MIMAT0009447 | UCAGGCCAGGCACAGUGGCUCA | 117.74 | 87.84 | 87.12 | 102.80 |
| hsa-miR-1973 | MIMAT0009448 | ACCGUGCAAAGGUAGCAUA | 112.50 | 89.12 | 99.28 | 89.66 |
| hsa-miR-197-3p | MIMAT0000227 | UUCACCACCUUCUCCACCCAGC | 54.84 | 55.26 | 92.84 | 85.51 |
| hsa-miR-1974 | MIMAT0009449 | UGGUUGUAGUCCGUGCGAGAAUA | 119.67 | 101.48 | 105.57 | 112.62 |
| hsa-miR-1975 | MIMAT0009450 | CCCCCACAACCGCGCUUGACUAGCU | 106.24 | 91.42 | 69.49 | 69.18 |
| hsa-miR-1976 | MIMAT0009451 | CCUCCUGCCCUCCUUGCUGU | 64.48 | 73.64 | 93.93 | 79.20 |
| hsa-miR-1977 | MIMAT0009452 | GAUUAGGGUGCUUAGCUGUUAA | 82.93 | 90.38 | 85.36 | 110.28 |
| hsa-miR-1978 | MIMAT0009453 | GGUUUGGUCCUAGCCUUUCUA | 75.49 | 77.59 | 121.89 | 104.09 |
| hsa-miR-1979 | MIMAT0009454 | CUCCCACUGCUUCACUUGACUA | 141.23 | 137.14 | 96.26 | 124.51 |
| hsa-miR-198 | MIMAT0000228 | GGUCCAGAGGGGAGAUAGGUUC | 117.20 | 115.10 | 106.39 | 93.73 |
| hsa-miR-199a-3p | MIMAT0000232 | ACAGUAGUCUGCACAUUGGUUA | 123.08 | 121.94 | 99.48 | 94.52 |
| hsa-miR-199a-5p | MIMAT0000231 | CCCAGUGUUCAGACUACCUGUUC | 80.32 | 72.95 | 108.55 | 90.87 |
| hsa-miR-199b-5p | MIMAT0000263 | CCCAGUGUUUAGACUAUCUGUUC | 90.49 | 118.78 | 86.22 | 90.23 |
| hsa-miR-19a-3p | MIMAT0000073 | UGUGCAAAUCUAUGCAAAACUGA | 107.81 | 113.91 | 117.27 | 115.72 |
| hsa-miR-19a-5p | MIMAT0004490 | AGUUUUGCAUAGUUGCACUACA | 50.23 | 49.19 | 124.73 | 109.32 |
| hsa-miR-19b-1-5p | MIMAT0004491 | AGUUUUGCAGGUUUGCAUCCAGC | 36.55 | 45.32 | 90.42 | 76.21 |
| hsa-miR-19b-2-5p | MIMAT0004492 | AGUUUUGCAGGUUUGCAUUUCA | 37.00 | 35.70 | 103.66 | 92.57 |
| hsa-miR-19b-3p | MIMAT0000074 | UGUGCAAAUCCAUGCAAAACUGA | 154.62 | 164.41 | 94.91 | 122.60 |
| hsa-miR-200a-3p | MIMAT0000682 | UAACACUGUCUGGUAACGAUGU | 82.95 | 80.66 | 110.49 | 92.03 |
| hsa-miR-200a-5p | MIMAT0001620 | CAUCUUACCGGACAGUGCUGGA | 82.47 | 98.23 | 100.73 | 91.69 |
| hsa-miR-200b-3p | MIMAT0000318 | UAAUACUGCCUGGUAAUGAUGA | 50.55 | 52.33 | 57.75 | 66.35 |
| hsa-miR-200b-5p | MIMAT0004571 | CAUCUUACUGGGCAGCAUUGGA | 75.58 | 89.15 | 119.57 | 113.45 |
| hsa-miR-200c-3p | MIMAT0000617 | UAAUACUGCCGGGUAAUGAUGGA | 82.47 | 83.97 | 61.63 | 63.38 |
| hsa-miR-200c-5p | MIMAT0004657 | CGUCUUACCCAGCAGUGUUUGG | 108.11 | 120.61 | 102.52 | 107.89 |
| hsa-miR-202-3p | MIMAT0002811 | AGAGGUAUAGGGCAUGGGAA | 91.09 | 85.98 | 135.72 | 103.64 |
| hsa-miR-202-5p | MIMAT0002810 | UUCCUAUGCAUAUACUUCUUUG | 97.50 | 90.18 | 106.19 | 93.41 |

| | | | | | | |
|---|---|---|---|---|---|---|
| hsa-miR-203a | MIMAT0000264 | GUGAAAUGUUUAGGACCACUAG | 106.73 | 110.55 | 110.04 | 101.95 |
| hsa-miR-204-5p | MIMAT0000265 | UUCCCUUUGUCAUCCUAUGCCU | 107.48 | 101.94 | 117.07 | 120.93 |
| hsa-miR-2052 | MIMAT0009977 | UGUUUUGAUAACAGUAAUGU | 40.43 | 35.71 | 100.73 | 92.51 |
| hsa-miR-2053 | MIMAT0009978 | GUGUUAAUUAAACCUCUAUUUAC | 53.83 | 45.30 | 74.24 | 92.61 |
| hsa-miR-205-3p | MIMAT0009197 | GAUUUCAGUGGAGUGAAGUUC | 40.99 | 53.26 | 83.90 | 86.81 |
| hsa-miR-2054 | MIMAT0009979 | CUGUAAUAUAAAUUUAAUUUAUU | 102.96 | 104.72 | 106.68 | 98.26 |
| hsa-miR-205-5p | MIMAT0000266 | UCCUUCAUUCCACCGGAGUCUG | 85.84 | 97.14 | 105.47 | 119.35 |
| hsa-miR-206 | MIMAT0000462 | UGGAAUGUAAGGAAGUGUGUGG | 143.45 | 121.41 | 80.81 | 73.61 |
| hsa-miR-208a | MIMAT0000241 | AUAAGACGAGCAAAAAGCUUGU | 109.11 | 120.13 | 56.69 | 46.31 |
| hsa-miR-208b | MIMAT0004960 | AUAAGACGAACAAAAGGUUUGU | 112.15 | 124.71 | 79.73 | 80.71 |
| hsa-miR-20a-3p | MIMAT0004493 | ACUGCAUUAUGAGCACUUAAAG | 117.03 | 108.50 | 107.64 | 84.78 |
| hsa-miR-20a-5p | MIMAT0000075 | UAAAGUGCUUAUAGUGCAGGUAG | 122.90 | 124.82 | 108.55 | 124.10 |
| hsa-miR-20b-3p | MIMAT0004752 | ACUGUAGUAUGGGCACUUCCAG | 111.46 | 126.50 | 120.13 | 124.59 |
| hsa-miR-20b-5p | MIMAT0001413 | CAAAGUGCUCAUAGUGCAGGUAG | 124.49 | 105.05 | 114.87 | 111.62 |
| hsa-miR-210 | MIMAT0000267 | CUGUGCGUGUGACAGCGGCUGA | 117.72 | 95.35 | 115.90 | 97.41 |
| hsa-miR-2110 | MIMAT0010133 | UUGGGGAAACGGCCGCUGAGUG | 84.81 | 74.52 | 106.51 | 84.41 |
| hsa-miR-2113 | MIMAT0009206 | AUUUGUGCUUGGCUCUGUCAC | 146.94 | 177.83 | 120.02 | 118.87 |
| hsa-miR-2114-3p | MIMAT0011157 | CGAGCCUCAAGCAAGGGACUU | 78.42 | 79.72 | 77.84 | 82.63 |
| hsa-miR-2114-5p | MIMAT0011156 | UAGUCCCUUCCUUGAAGCGGUC | 97.35 | 108.45 | 86.13 | 84.46 |
| hsa-miR-2115-3p | MIMAT0011159 | CAUCAGAAUUCAUGGAGGCUAG | 67.88 | 70.54 | 85.66 | 79.24 |
| hsa-miR-2115-5p | MIMAT0011158 | AGCUUCCAUGACUCCUGAUGGA | 71.96 | 66.09 | 97.82 | 82.85 |
| hsa-miR-211-5p | MIMAT0000268 | UUCCCUUUGUCAUCCUUCGCCU | 112.02 | 92.64 | 93.49 | 101.68 |
| hsa-miR-2116-3p | MIMAT0011161 | CCUCCCAUGCCAAGAACUCCC | 89.02 | 77.22 | 87.87 | 79.21 |
| hsa-miR-2116-5p | MIMAT0011160 | GGUUCUUAGCAUAGGAGGUCU | 76.57 | 78.94 | 85.09 | 128.63 |
| hsa-miR-2117 | MIMAT0011162 | UGUUCUCUUUGCCAAGGACAG | 85.49 | 92.42 | 117.80 | 112.80 |
| hsa-miR-212-3p | MIMAT0000269 | UAACAGUCUCCAGUCACGGCC | 105.69 | 107.21 | 62.78 | 81.69 |
| hsa-miR-21-3p | MIMAT0004494 | CAACACCAGUCGAUGGGCUGU | 86.67 | 83.73 | 97.39 | 89.49 |
| hsa-miR-214-3p | MIMAT0000271 | ACAGCAGGCACAGACAGGCAGU | 113.07 | 138.59 | 86.42 | 93.32 |
| hsa-miR-214-5p | MIMAT0004564 | UGCCUGUCUACACUUGCUGUGC | 121.57 | 78.40 | 94.27 | 79.61 |
| hsa-miR-215 | MIMAT0000272 | AUGACCUAUGAAUUGACAGAC | 90.86 | 96.00 | 107.37 | 116.87 |
| hsa-miR-21-5p | MIMAT0000076 | UAGCUUAUCAGACUGAUGUUGA | 110.59 | 121.46 | 112.80 | 105.38 |
| hsa-miR-216a-5p | MIMAT0000273 | UAAUCUCAGCUGGCAACUGUGA | 116.50 | 124.28 | 96.76 | 99.23 |
| hsa-miR-216b | MIMAT0004959 | AAAUCUCUGCAGGCAAAUGUGA | 169.78 | 172.43 | 116.53 | 118.25 |
| hsa-miR-217 | MIMAT0000274 | UACUGCAUCAGGAACUGAUUGGA | 109.75 | 99.72 | 64.39 | 80.92 |
| hsa-miR-218-1-3p | MIMAT0004565 | AUGGUUCCGUCAAGCACCAUGG | 91.81 | 110.29 | 72.19 | 62.93 |
| hsa-miR-218-2-3p | MIMAT0004566 | CAUGGUUCUGUCAAGCACCGCG | 112.15 | 106.28 | 89.33 | 107.52 |
| hsa-miR-218-5p | MIMAT0000275 | UUGUGCUUGAUCUAACCAUGU | 83.07 | 86.29 | 99.94 | 99.21 |
| hsa-miR-219-1-3p | MIMAT0004567 | AGAGUUGAGUCUGGACGUCCCG | 78.15 | 99.72 | 111.21 | 116.45 |
| hsa-miR-219-2-3p | MIMAT0004675 | AGAAUUGUGGCUGGACAUCUGU | 49.33 | 57.21 | 138.44 | 141.06 |
| hsa-miR-219-5p | MIMAT0000276 | UGAUUGUCCAAACGCAAUUCU | 84.15 | 98.59 | 73.01 | 90.63 |
| hsa-miR-220a | MIMAT0000277 | CCACACCGUAUCUGACACUUU | 80.32 | 93.05 | 105.19 | 91.69 |

| | | | | | | |
|---|---|---|---|---|---|---|
| hsa-miR-220b | MIMAT0004908 | CCACCACCGUGUCUGACACUU | 92.26 | 82.20 | 105.94 | 104.09 |
| hsa-miR-221-3p | MIMAT0000278 | AGCUACAUUGUCUGCUGGGUUUC | 142.31 | 139.73 | 110.78 | 120.82 |
| hsa-miR-221-5p | MIMAT0004568 | ACCUGGCAUACAAUGUAGAUUU | 108.04 | 92.67 | 97.28 | 104.46 |
| hsa-miR-222-3p | MIMAT0000279 | AGCUACAUCUGGCUACUGGGU | 97.75 | 118.23 | 79.09 | 87.81 |
| hsa-miR-222-5p | MIMAT0004569 | CUCAGUAGCCAGUGUAGAUCCU | 45.34 | 106.70 | 89.53 | 91.03 |
| hsa-miR-223-3p | MIMAT0000280 | UGUCAGUUUGUCAAAUACCCCA | 113.82 | 98.12 | 82.72 | 76.97 |
| hsa-miR-223-5p | MIMAT0004570 | CGUGUAUUUGACAAGCUGAGUU | 83.86 | 74.16 | 83.23 | 82.97 |
| hsa-miR-22-3p | MIMAT0000077 | AAGCUGCCAGUUGAAGAACUGU | 56.06 | 43.12 | 78.01 | 67.08 |
| hsa-miR-224-3p | MIMAT0009198 | AAAAUGGUGCCCUAGUGACUACA | 65.71 | 61.75 | 99.32 | 110.46 |
| hsa-miR-224-5p | MIMAT0000281 | CAAGUCACUAGUGGUUCCGUU | 76.04 | 63.08 | 109.42 | 104.22 |
| hsa-miR-22-5p | MIMAT0004495 | AGUUCUUCAGUGGCAAGCUUUA | 90.29 | 103.40 | 85.12 | 117.66 |
| hsa-miR-2276 | MIMAT0011775 | UCUGCAAGUGUCAGAGGCGAGG | 89.12 | 89.44 | 98.34 | 105.48 |
| hsa-miR-2277-3p | MIMAT0011777 | UGACAGCGCCCUGCCUGGCUC | 103.84 | 120.29 | 120.76 | 115.88 |
| hsa-miR-2277-5p | MIMAT0017352 | AGCGCGGGCUGAGCGCUGCCAGUC | 62.31 | 46.63 | 95.20 | 82.85 |
| hsa-miR-2278 | MIMAT0011778 | GAGAGCAGUGUGUGUUGCCUGG | 101.10 | 126.54 | 96.74 | 97.36 |
| hsa-miR-2355-3p | MIMAT0017950 | AUUGUCCUUGCUGUUUGGAGAU | 89.56 | 97.60 | 120.80 | 114.49 |
| hsa-miR-2355-5p | MIMAT0016895 | AUCCCCAGAUACAAUGGACAA | 144.95 | 144.71 | 103.03 | 119.72 |
| hsa-miR-23a-3p | MIMAT0000078 | AUCACAUUGCCAGGGAUUUCC | 123.02 | 117.53 | 73.47 | 88.85 |
| hsa-miR-23a-5p | MIMAT0004496 | GGGGUUCCUGGGGAUGGGAUUU | 120.64 | 136.43 | 88.90 | 97.80 |
| hsa-miR-23b-3p | MIMAT0000418 | AUCACAUUGCCAGGGAUUACC | 103.77 | 112.40 | 116.79 | 98.04 |
| hsa-miR-23b-5p | MIMAT0004587 | UGGGUUCCUGGCAUGCUGAUUU | 106.45 | 78.96 | 77.14 | 73.53 |
| hsa-miR-23c | MIMAT0018000 | AUCACAUUGCCAGUGAUUACCC | 85.10 | 101.04 | 75.80 | 90.37 |
| hsa-miR-24-1-5p | MIMAT0000079 | UGCCUACUGAGCUGAUAUCAGU | 97.27 | 78.96 | 96.88 | 84.56 |
| hsa-miR-24-2-5p | MIMAT0004497 | UGCCUACUGAGCUGAAACACAG | 93.84 | 96.75 | 83.13 | 84.17 |
| hsa-miR-24-3p | MIMAT0000080 | UGGCUCAGUUCAGCAGGAACAG | 93.28 | 105.52 | 106.94 | 125.15 |
| hsa-miR-25-3p | MIMAT0000081 | CAUUGCACUUGUCUCGGUCUGA | 63.24 | 65.68 | 97.73 | 78.77 |
| hsa-miR-25-5p | MIMAT0004498 | AGGCGGAGACUUGGGCAAUUG | 99.22 | 93.27 | 91.90 | 110.65 |
| hsa-miR-26a-1-3p | MIMAT0004499 | CCUAUUCUUGGUUACUUGCACG | 93.69 | 103.54 | 103.25 | 107.42 |
| hsa-miR-26a-2-3p | MIMAT0004681 | CCUAUUCUUGAUUACUUGUUUC | 110.79 | 88.07 | 100.29 | 86.49 |
| hsa-miR-26a-5p | MIMAT0000082 | UUCAAGUAAUCCAGGAUAGGCU | 113.07 | 120.86 | 99.95 | 80.42 |
| hsa-miR-26b-3p | MIMAT0004500 | CCUGUUCUCCAUUACUUGGCUC | 98.64 | 91.73 | 82.79 | 105.50 |
| hsa-miR-26b-5p | MIMAT0000083 | UUCAAGUAAUUCAGGAUAGGU | 70.46 | 91.93 | 76.84 | 105.02 |
| hsa-miR-27a-3p | MIMAT0000084 | UUCACAGUGGCUAAGUUCCGC | 170.19 | 144.71 | 111.55 | 127.42 |
| hsa-miR-27a-5p | MIMAT0004501 | AGGGCUUAGCUGCUUGUGAGCA | 112.61 | 89.93 | 128.15 | 121.73 |
| hsa-miR-27b-3p | MIMAT0000419 | UUCACAGUGGCUAAGUUCUGC | 132.98 | 149.59 | 104.30 | 89.81 |
| hsa-miR-27b-5p | MIMAT0004588 | AGAGCUUAGCUGAUUGGUGAAC | 98.79 | 112.92 | 95.12 | 104.72 |
| hsa-miR-28-3p | MIMAT0004502 | CACUAGAUUGUGAGCUCCUGGA | 54.99 | 26.89 | 111.63 | 94.57 |
| hsa-miR-28-5p | MIMAT0000085 | AAGGAGCUCACAGUCUAUUGAG | 40.74 | 34.42 | 73.17 | 72.82 |
| hsa-miR-2861 | MIMAT0013802 | GGGGCCUGGCGGUGGGCGG | 66.92 | 88.91 | 100.30 | 94.96 |
| hsa-miR-2909 | MIMAT0013863 | GUUAGGGCCAACAUCUCUUGG | 78.10 | 95.31 | 103.97 | 79.61 |
| hsa-miR-296-3p | MIMAT0004679 | GAGGGUUGGGUGGAGGCUCUCC | 100.76 | 100.24 | 95.92 | 88.42 |

| | | | | | | |
|---|---|---|---|---|---|---|
| hsa-miR-296-5p | MIMAT0000690 | AGGGCCCCCCCUCAAUCCUGU | 75.42 | 93.65 | 95.40 | 99.61 |
| hsa-miR-297 | MIMAT0004450 | AUGUAUGUGUGCAUGUGCAUG | 60.75 | 60.73 | 84.53 | 79.10 |
| hsa-miR-298 | MIMAT0004901 | AGCAGAAGCAGGGAGGUUCUCCCA | 84.61 | 98.50 | 108.71 | 97.57 |
| hsa-miR-299-3p | MIMAT0000687 | UAUGUGGGAUGGUAAACCGCUU | 128.60 | 104.32 | 99.51 | 110.03 |
| hsa-miR-299-5p | MIMAT0002890 | UGGUUUACCGUCCCACAUACAU | 96.31 | 85.59 | 94.84 | 82.38 |
| hsa-miR-29a-3p | MIMAT0000086 | UAGCACCAUCUGAAAUCGGUUA | 54.08 | 79.25 | 67.99 | 84.28 |
| hsa-miR-29a-5p | MIMAT0004503 | ACUGAUUUCUUUUGGUGUUCAG | 112.31 | 125.79 | 110.65 | 129.72 |
| hsa-miR-29b-1-5p | MIMAT0004514 | GCUGGUUUCAUAUGGUGGUUUAGA | 93.52 | 101.83 | 125.15 | 124.59 |
| hsa-miR-29b-2-5p | MIMAT0004515 | CUGGUUUCACAUGGUGGCUUAG | 74.65 | 61.98 | 108.65 | 97.69 |
| hsa-miR-29b-3p | MIMAT0000100 | UAGCACCAUUUGAAAUCAGUGUU | 118.09 | 132.64 | 101.97 | 72.19 |
| hsa-miR-29c-3p | MIMAT0000681 | UAGCACCAUUUGAAAUCGGUUA | 73.93 | 91.96 | 44.61 | 55.13 |
| hsa-miR-29c-5p | MIMAT0004673 | UGACCGAUUUCUCCUGGUGUUC | 95.15 | 90.44 | 113.92 | 113.19 |
| hsa-miR-300 | MIMAT0004903 | UAUACAAGGGCAGACUCUCUCU | 126.73 | 137.37 | 97.91 | 116.74 |
| hsa-miR-301a-3p | MIMAT0000688 | CAGUGCAAUAGUAUUGUCAAAGC | 86.15 | 103.73 | 113.09 | 93.98 |
| hsa-miR-301b | MIMAT0004958 | CAGUGCAAUGAUAUUGUCAAAGC | 200.53 | 207.16 | 116.74 | 113.85 |
| hsa-miR-302a-3p | MIMAT0000684 | UAAGUGCUUCCAUGUUUUGGUGA | 123.05 | 121.62 | 146.53 | 148.85 |
| hsa-miR-302a-5p | MIMAT0000683 | ACUUAAACGUGGAUGUACUUGCU | 120.90 | 72.56 | 107.17 | 98.25 |
| hsa-miR-302b-3p | MIMAT0000715 | UAAGUGCUUCCAUGUUUUAGUAG | 123.02 | 126.75 | 129.86 | 126.30 |
| hsa-miR-302b-5p | MIMAT0000714 | ACUUUAACAUGGAAGUGCUUUC | 88.64 | 86.87 | 96.20 | 93.50 |
| hsa-miR-302c-3p | MIMAT0000717 | UAAGUGCUUCCAUGUUUCAGUGG | 185.42 | 233.52 | 130.29 | 123.47 |
| hsa-miR-302c-5p | MIMAT0000716 | UUUAACAUGGGGGUACCUGCUG | 116.50 | 106.78 | 112.77 | 109.16 |
| hsa-miR-302d-3p | MIMAT0000718 | UAAGUGCUUCCAUGUUUGAGUGU | 89.25 | 89.85 | 134.07 | 144.28 |
| hsa-miR-302d-5p | MIMAT0004685 | ACUUUAACAUGGAGGCACUUGC | 92.67 | 115.93 | 71.08 | 76.21 |
| hsa-miR-302e | MIMAT0005931 | UAAGUGCUUCCAUGCUU | 146.11 | 139.37 | 143.10 | 133.24 |
| hsa-miR-302f | MIMAT0005932 | UAAUUGCUUCCAUGUUU | 118.05 | 99.53 | 82.11 | 105.64 |
| hsa-miR-3065-3p | MIMAT0015378 | UCAGCACCAGGAUAUUGUUGGAG | 90.69 | 78.54 | 99.51 | 91.43 |
| hsa-miR-3065-5p | MIMAT0015066 | UCAACAAAAUCACUGAUGCUGGA | 128.50 | 114.95 | 108.00 | 122.54 |
| hsa-miR-3074-3p | MIMAT0015027 | GAUAUCAGCUCAGUAGGCACCG | 42.83 | 52.05 | 85.07 | 65.69 |
| hsa-miR-30a-3p | MIMAT0000088 | CUUUCAGUCGGAUGUUUGCAGC | 129.95 | 118.27 | 69.39 | 81.23 |
| hsa-miR-30a-5p | MIMAT0000087 | UGUAAACAUCCUCGACUGGAAG | 94.27 | 78.71 | 104.53 | 98.77 |
| hsa-miR-30b-3p | MIMAT0004589 | CUGGGAGGUGGAUGUUUACUUC | 118.82 | 128.78 | 102.76 | 80.18 |
| hsa-miR-30b-5p | MIMAT0000420 | UGUAAACAUCCUACACUCAGCU | 118.05 | 104.07 | 107.06 | 112.21 |
| hsa-miR-30c-1-3p | MIMAT0004674 | CUGGGAGAGGGUUGUUUACUCC | 102.06 | 104.65 | 134.62 | 121.16 |
| hsa-miR-30c-2-3p | MIMAT0004550 | CUGGGAGAAGGCUGUUUACUCU | 121.07 | 107.99 | 110.42 | 127.52 |
| hsa-miR-30c-5p | MIMAT0000244 | UGUAAACAUCCUACACUCUCAGC | 62.44 | 55.93 | 81.77 | 93.80 |
| hsa-miR-30d-3p | MIMAT0004551 | CUUUCAGUCAGAUGUUUGCUGC | 102.91 | 120.86 | 86.13 | 83.06 |
| hsa-miR-30d-5p | MIMAT0000245 | UGUAAACAUCCCCGACUGGAAG | 69.99 | 78.45 | 119.30 | 109.34 |
| hsa-miR-30e-3p | MIMAT0000693 | CUUUCAGUCGGAUGUUUACAGC | 104.62 | 27.13 | 78.88 | 44.10 |
| hsa-miR-30e-5p | MIMAT0000692 | UGUAAACAUCCUUGACUGGAAG | 101.04 | 89.22 | 119.54 | 109.60 |
| hsa-miR-3115 | MIMAT0014977 | AUAUGGGUUUACUAGUUGGU | 81.25 | 117.34 | 68.31 | 90.93 |
| hsa-miR-3116 | MIMAT0014978 | UGCCUGGAACAUAGUAGGGACU | 84.89 | 82.42 | 109.60 | 137.02 |

| | | | | | | |
|---|---|---|---|---|---|---|
| hsa-miR-3117-3p | MIMAT0014979 | AUAGGACUCAUAUAGUGCCAG | 85.77 | 97.58 | 84.64 | 83.11 |
| hsa-miR-3118 | MIMAT0014980 | UGUGACUGCAUUAUGAAAAUUCU | 108.26 | 69.40 | 112.27 | 101.62 |
| hsa-miR-3119 | MIMAT0014981 | UGGCUUUUAACUUUGAUGGC | 66.25 | 61.63 | 86.07 | 104.30 |
| hsa-miR-3120-3p | MIMAT0014982 | CACAGCAAGUGUAGACAGGCA | 97.91 | 109.31 | 122.03 | 114.28 |
| hsa-miR-3121-3p | MIMAT0014983 | UAAAUAGAGUAGGCAAAGGACA | 125.85 | 115.49 | 71.29 | 76.78 |
| hsa-miR-3122 | MIMAT0014984 | GUUGGGACAAGAGGACGGUCUU | 76.87 | 107.66 | 97.28 | 96.58 |
| hsa-miR-3123 | MIMAT0014985 | CAGAGAAUUGUUUAAUC | 175.93 | 142.40 | 127.65 | 101.27 |
| hsa-miR-3124-5p | MIMAT0014986 | UUCGCGGGCGAAGGCAAAGUC | 92.68 | 101.94 | 120.80 | 108.06 |
| hsa-miR-3125 | MIMAT0014988 | UAGAGGAAGCUGUGGAGAGA | 100.35 | 103.93 | 97.84 | 95.78 |
| hsa-miR-3126-3p | MIMAT0015377 | CAUCUGGCAUCCGUCACACAGA | 89.56 | 107.36 | 100.00 | 73.74 |
| hsa-miR-3126-5p | MIMAT0014989 | UGAGGGACAGAUGCCAGAAGCA | 121.87 | 115.42 | 101.78 | 105.09 |
| hsa-miR-3127-5p | MIMAT0014990 | AUCAGGGCUUGUGGAAUGGGAAG | 109.22 | 112.44 | 124.42 | 104.76 |
| hsa-miR-3128 | MIMAT0014991 | UCUGGCAAGUAAAAAACUCUCAU | 65.20 | 78.81 | 107.64 | 121.30 |
| hsa-miR-3129-5p | MIMAT0014992 | GCAGUAGUGUAGAGAUUGGUUU | 117.45 | 127.33 | 82.32 | 79.95 |
| hsa-miR-3130-3p | MIMAT0014994 | GCUGCACCGGAGACUGGGUAA | 79.22 | 72.37 | 100.31 | 107.81 |
| hsa-miR-3130-5p | MIMAT0014995 | UACCCAGUCUCCGGUGCAGCC | 80.28 | 89.43 | 95.97 | 93.77 |
| hsa-miR-3131 | MIMAT0014996 | UCGAGGACUGGUGGAAGGGCCUU | 83.26 | 124.16 | 115.03 | 102.51 |
| hsa-miR-3132 | MIMAT0014997 | UGGGUAGAGAAGGAGCUCAGAGGA | 93.66 | 90.84 | 95.07 | 101.72 |
| hsa-miR-3133 | MIMAT0014998 | UAAAGAACUCUUAAAACCCAAU | 60.02 | 68.42 | 75.39 | 78.60 |
| hsa-miR-3134 | MIMAT0015000 | UGAUGGAUAAAAGACUACAUAUU | 86.91 | 99.64 | 88.12 | 110.22 |
| hsa-miR-3135a | MIMAT0015001 | UGCCUAGGCUGAGACUGCAGUG | 84.81 | 112.63 | 85.66 | 101.23 |
| hsa-miR-3136-5p | MIMAT0015003 | CUGACUGAAUAGGUAGGGUCAUU | 78.12 | 82.82 | 97.29 | 119.72 |
| hsa-miR-3137 | MIMAT0015005 | UCUGUAGCCUGGGAGCAAUGGGGU | 95.35 | 78.96 | 140.39 | 138.44 |
| hsa-miR-3138 | MIMAT0015006 | UGUGGACAGUGAGGUAGAGGGAGU | 70.84 | 65.69 | 106.19 | 114.43 |
| hsa-miR-3139 | MIMAT0015007 | UAGGAGCUCAACAGAUGCCUGUU | 84.11 | 60.73 | 65.33 | 61.94 |
| hsa-miR-31-3p | MIMAT0004504 | UGCUAUGCCAACAUAUUGCCAU | 110.60 | 106.77 | 112.33 | 139.14 |
| hsa-miR-3140-3p | MIMAT0015008 | AGCUUUUGGGAAUUCAGGUAGU | 85.77 | 99.88 | 93.82 | 111.38 |
| hsa-miR-3141 | MIMAT0015010 | GAGGGCGGGUGGAGGAGGA | 92.30 | 88.04 | 99.57 | 116.86 |
| hsa-miR-3142 | MIMAT0015011 | AAGGCCUUUCUGAACCUUCAGA | 101.98 | 77.22 | 124.81 | 83.66 |
| hsa-miR-3143 | MIMAT0015012 | AUAACAUUGUAAAGCGCUUCUUUCG | 130.06 | 146.40 | 114.93 | 126.83 |
| hsa-miR-3144-3p | MIMAT0015015 | AUAUACCUGUUCGGUCUCUUUA | 125.38 | 120.28 | 84.98 | 118.38 |
| hsa-miR-3144-5p | MIMAT0015014 | AGGGGACCAAAGAGAUAUAUAG | 73.21 | 65.07 | 76.00 | 76.95 |
| hsa-miR-3145-3p | MIMAT0015016 | AGAUAUUUUGAGUGUUUGGAAUUG | 79.06 | 105.20 | 82.01 | 95.87 |
| hsa-miR-3146 | MIMAT0015018 | CAUGCUAGGAUAGAAAGAAUGG | 98.13 | 121.46 | 84.00 | 87.14 |
| hsa-miR-3147 | MIMAT0015019 | GGUUGGGCAGUGAGGAGGGUGUGA | 77.69 | 82.76 | 121.40 | 115.60 |
| hsa-miR-3148 | MIMAT0015021 | UGGAAAAAACUGGUGUGUGCUU | 108.63 | 126.76 | 133.10 | 128.28 |
| hsa-miR-3149 | MIMAT0015022 | UUUGUAUGGAUAUGUGUGUGUAU | 118.82 | 107.26 | 78.23 | 88.35 |
| hsa-miR-3150a-3p | MIMAT0015023 | CUGGGGAGAUCCUCGAGGUUGG | 59.00 | 75.14 | 111.55 | 118.10 |
| hsa-miR-3150b-3p | MIMAT0018194 | UGAGGAGAUCGUCGAGGUUGG | 90.38 | 112.71 | 99.24 | 123.77 |
| hsa-miR-3151 | MIMAT0015024 | GGUGGGGCAAUGGGAUCAGGU | 86.15 | 75.83 | 103.59 | 92.02 |
| hsa-miR-3152-3p | MIMAT0015025 | UGUGUUAGAAUAGGGGCAAUAA | 121.87 | 104.72 | 100.62 | 112.83 |

| | | | | | | |
|---|---|---|---|---|---|---|
| hsa-miR-3153 | MIMAT0015026 | GGGGAAAGCGAGUAGGGACAUUU | 164.33 | 134.47 | 135.73 | 134.87 |
| hsa-miR-3154 | MIMAT0015028 | CAGAAGGGGAGUUGGGAGCAGA | 87.59 | 88.70 | 83.14 | 87.96 |
| hsa-miR-3155a | MIMAT0015029 | CCAGGCUCUGCAGUGGGAACU | 150.31 | 129.05 | 114.93 | 120.93 |
| hsa-miR-3156-5p | MIMAT0015030 | AAAGAUCUGGAAGUGGGAGACA | 104.81 | 109.88 | 102.28 | 89.45 |
| hsa-miR-3157-5p | MIMAT0015031 | UUCAGCCAGGCUAGUGCAGUCU | 82.20 | 69.57 | 82.66 | 85.28 |
| hsa-miR-3158-3p | MIMAT0015032 | AAGGGCUUCCUCUCUGCAGGAC | 106.21 | 102.17 | 98.92 | 114.52 |
| hsa-miR-3159 | MIMAT0015033 | UAGGAUUACAAGUGUCGGCCAC | 164.05 | 180.89 | 114.39 | 130.66 |
| hsa-miR-31-5p | MIMAT0000089 | AGGCAAGAUGCUGGCAUAGCU | 84.58 | 86.58 | 133.23 | 137.00 |
| hsa-miR-3160-3p | MIMAT0015034 | AGAGCUGAGACUAGAAAGCCCA | 132.73 | 116.71 | 98.24 | 93.50 |
| hsa-miR-3161 | MIMAT0015035 | CUGAUAAGAACAGAGGCCCAGAU | 84.74 | 85.68 | 90.08 | 92.22 |
| hsa-miR-3162-5p | MIMAT0015036 | UUAGGGAGUAGAAGGGUGGGGAG | 86.91 | 106.34 | 116.01 | 87.77 |
| hsa-miR-3163 | MIMAT0015037 | UAUAAAAUGAGGGCAGUAAGAC | 97.53 | 124.25 | 125.86 | 124.01 |
| hsa-miR-3164 | MIMAT0015038 | UGUGACUUUAAGGGAAAUGGCG | 101.11 | 117.37 | 108.67 | 104.62 |
| hsa-miR-3165 | MIMAT0015039 | AGGUGGAUGCAAUGUGACCUCA | 42.63 | 46.61 | 61.34 | 75.15 |
| hsa-miR-3166 | MIMAT0015040 | CGCAGACAAUGCCUACUGGCCUA | 75.38 | 82.09 | 93.08 | 76.78 |
| hsa-miR-3167 | MIMAT0015042 | AGGAUUUCAGAAAUACUGGUGU | 91.78 | 108.28 | 84.29 | 105.46 |
| hsa-miR-3168 | MIMAT0015043 | GAGUUCUACAGUCAGAC | 111.67 | 84.98 | 100.56 | 90.23 |
| hsa-miR-3169 | MIMAT0015044 | UAGGACUGUGCUUGGCACAUAG | 56.96 | 52.87 | 119.13 | 116.48 |
| hsa-miR-3170 | MIMAT0015045 | CUGGGGUUCUGAGACAGACAGU | 133.06 | 143.82 | 139.03 | 151.18 |
| hsa-miR-3171 | MIMAT0015046 | AGAUGUAUGGAAUCUGUAUAUAUC | 85.29 | 92.31 | 93.26 | 101.67 |
| hsa-miR-3173-3p | MIMAT0015048 | AAAGGAGGAAAUAGGCAGGCCA | 73.28 | 81.16 | 101.82 | 102.79 |
| hsa-miR-3174 | MIMAT0015051 | UAGUGAGUUAGAGAUGCAGAGCC | 60.53 | 76.59 | 92.20 | 98.05 |
| hsa-miR-3175 | MIMAT0015052 | CGGGGAGAGAACGCAGUGACGU | 81.51 | 95.31 | 112.50 | 94.60 |
| hsa-miR-3176 | MIMAT0015053 | ACUGGCCUGGGACUACCGG | 125.63 | 97.58 | 126.67 | 111.61 |
| hsa-miR-3177-3p | MIMAT0015054 | UGCACGGCACUGGGGACACGU | 28.82 | 37.96 | 92.00 | 82.32 |
| hsa-miR-3178 | MIMAT0015055 | GGGGCGCGGCCGGAUCG | 95.97 | 97.61 | 119.44 | 119.88 |
| hsa-miR-3179 | MIMAT0015056 | AGAAGGGGUGAAAUUUAAACGU | 81.94 | 88.40 | 83.28 | 102.68 |
| hsa-miR-3180 | MIMAT0018178 | UGGGGCGGAGCUUCCGGAG | 73.63 | 113.65 | 105.04 | 116.69 |
| hsa-miR-3180-3p | MIMAT0015058 | UGGGGCGGAGCUUCCGGAGGCC | 91.12 | 72.66 | 121.33 | 137.02 |
| hsa-miR-3180-5p | MIMAT0015057 | CUUCCAGACGCUCCGCCCCACGUCG | 88.36 | 103.54 | 101.78 | 105.09 |
| hsa-miR-3181 | MIMAT0015061 | AUCGGGCCCUCGGCGCCGG | 54.91 | 60.53 | 118.18 | 78.40 |
| hsa-miR-3182 | MIMAT0015062 | GCUUCUGUAGUGUAGUC | 134.06 | 152.53 | 117.47 | 102.36 |
| hsa-miR-3183 | MIMAT0015063 | GCCUCUCUCGGAGUCGCUCGGA | 113.73 | 109.08 | 122.79 | 117.15 |
| hsa-miR-3184-5p | MIMAT0015064 | UGAGGGGCCUCAGACCGAGCUUUU | 65.14 | 53.57 | 103.50 | 95.82 |
| hsa-miR-3185 | MIMAT0015065 | AGAAGAAGGCGGUCGGUCUGCGG | 138.13 | 153.06 | 104.45 | 120.13 |
| hsa-miR-3186-3p | MIMAT0015068 | UCACGCGGAGAGAUGGCUUUG | 59.19 | 41.21 | 85.60 | 91.97 |
| hsa-miR-3186-5p | MIMAT0015067 | CAGGCGUCUGUCUACGUGGCUU | 72.11 | 99.17 | 85.09 | 77.57 |
| hsa-miR-3187-3p | MIMAT0015069 | UUGGCCAUGGGGCUGCGCGG | 93.46 | 124.71 | 108.53 | 131.65 |
| hsa-miR-3188 | MIMAT0015070 | AGAGGCUUUGUGCGGAUACGGGG | 131.77 | 107.26 | 106.19 | 90.30 |
| hsa-miR-3189-3p | MIMAT0015071 | CCCUUGGGUCUGAUGGGGUAG | 88.45 | 64.50 | 95.55 | 84.50 |
| hsa-miR-3190 | MIMAT0015073 | UGUGGAAGGUAGACGGCCAGAGA | 123.81 | 131.49 | 125.29 | 108.38 |

| | | | | | | |
|---|---|---|---|---|---|---|
| hsa-miR-3191-3p | MIMAT0015075 | UGGGGACGUAGCUGGCCAGACAG | 95.16 | 88.36 | 112.97 | 107.16 |
| hsa-miR-3192 | MIMAT0015076 | UCUGGGAGGUUGUAGCAGUGGAA | 92.30 | 71.48 | 70.70 | 61.29 |
| hsa-miR-3193 | MIMAT0015077 | UCCUGCGUAGGAUCUGAGGAGU | 74.69 | 98.79 | 87.39 | 113.64 |
| hsa-miR-3194-5p | MIMAT0015078 | GGCCAGCCACCAGGAGGGCUG | 127.73 | 108.45 | 81.87 | 64.62 |
| hsa-miR-3195 | MIMAT0015079 | CGCGCCGGGCCCGGGUU | 111.97 | 108.00 | 95.89 | 82.13 |
| hsa-miR-3196 | MIMAT0015080 | CGGGGCGGCAGGGGCCUC | 95.16 | 88.36 | 98.29 | 114.86 |
| hsa-miR-3197 | MIMAT0015082 | GGAGGCGCAGGCUCGGAAAGGCG | 94.56 | 108.13 | 138.78 | 127.39 |
| hsa-miR-3198 | MIMAT0015083 | GUGGAGUCCUGGGGAAUGGAGA | 151.77 | 180.89 | 83.60 | 71.91 |
| hsa-miR-3199 | MIMAT0015084 | AGGGACUGCCUUAGGAGAAAGUU | 121.87 | 116.46 | 94.50 | 89.87 |
| hsa-miR-3200-3p | MIMAT0015085 | CACCUUGCGCUACUCAGGUCUG | 89.95 | 115.43 | 82.88 | 93.80 |
| hsa-miR-3200-5p | MIMAT0017392 | AAUCUGAGAAGGCGCACAAGGU | 83.03 | 74.48 | 108.48 | 102.55 |
| hsa-miR-3201 | MIMAT0015086 | GGGAUAUGAAGAAAAAU | 117.60 | 123.63 | 113.33 | 111.27 |
| hsa-miR-3202 | MIMAT0015089 | UGGAAGGGAGAAGAGCUUUAAU | 125.70 | 99.88 | 98.58 | 97.85 |
| hsa-miR-320a | MIMAT0000510 | AAAAGCUGGGUUGAGAGGGCGA | 70.62 | 68.41 | 98.12 | 85.54 |
| hsa-miR-320b | MIMAT0005792 | AAAAGCUGGGUUGAGAGGGCAA | 120.48 | 111.87 | 127.59 | 135.09 |
| hsa-miR-320c | MIMAT0005793 | AAAAGCUGGGUUGAGAGGGU | 86.83 | 98.74 | 97.88 | 123.20 |
| hsa-miR-320d | MIMAT0006764 | AAAAGCUGGGUUGAGAGGA | 112.50 | 122.45 | 92.20 | 78.84 |
| hsa-miR-320e | MIMAT0015072 | AAAGCUGGGUUGAGAAGG | 118.23 | 114.14 | 105.12 | 115.34 |
| hsa-miR-323a-3p | MIMAT0000755 | CACAUUACACGGUCGACCUCU | 73.44 | 86.87 | 69.63 | 81.51 |
| hsa-miR-323a-5p | MIMAT0004696 | AGGUGGUCCGUGGCGCGUUCGC | 85.38 | 91.52 | 92.99 | 90.38 |
| hsa-miR-323b-3p | MIMAT0015050 | CCCAAUACACGGUCGACCUCUU | 69.58 | 52.18 | 79.91 | 74.92 |
| hsa-miR-323b-5p | MIMAT0001630 | AGGUUGUCCGUGGUGAGUUCGCA | 107.40 | 90.18 | 111.59 | 97.69 |
| hsa-miR-32-3p | MIMAT0004505 | CAAUUUAGUGUGUGUGAUAUUU | 112.02 | 128.25 | 112.75 | 123.17 |
| hsa-miR-324-3p | MIMAT0000762 | ACUGCCCCAGGUGCUGCUGG | 96.55 | 117.16 | 101.63 | 109.47 |
| hsa-miR-324-5p | MIMAT0000761 | CGCAUCCCCUAGGGCAUUGGUGU | 74.34 | 82.25 | 103.99 | 83.06 |
| hsa-miR-325 | MIMAT0000771 | CCUAGUAGGUGUCCAGUAAGUGU | 94.39 | 84.31 | 109.45 | 101.47 |
| hsa-miR-32-5p | MIMAT0000090 | UAUUGCACAUUACUAAGUUGCA | 86.67 | 74.03 | 99.25 | 88.63 |
| hsa-miR-326 | MIMAT0000756 | CCUCUGGGCCCUUCCUCCAG | 76.41 | 97.04 | 78.06 | 61.68 |
| hsa-miR-328 | MIMAT0000752 | CUGGCCCUCUCUGCCCUUCCGU | 183.06 | 180.43 | 84.95 | 103.13 |
| hsa-miR-329 | MIMAT0001629 | AACACACCUGGUUAACCUCUUU | 76.53 | 71.45 | 82.37 | 98.82 |
| hsa-miR-330-3p | MIMAT0000751 | GCAAAGCACACGGCCUGCAGAGA | 140.42 | 154.18 | 105.68 | 113.43 |
| hsa-miR-330-5p | MIMAT0004693 | UCUCUGGGCCUGUGUCUUAGGC | 102.21 | 102.13 | 141.60 | 121.45 |
| hsa-miR-331-3p | MIMAT0000760 | GCCCCUGGGCCUAUCCUAGAA | 112.24 | 130.82 | 125.31 | 99.03 |
| hsa-miR-331-5p | MIMAT0004700 | CUAGGUAUGGUCCCAGGGAUCC | 107.42 | 110.88 | 94.68 | 78.28 |
| hsa-miR-335-3p | MIMAT0004703 | UUUUUCAUUAUUGCUCCUGACC | 81.56 | 85.62 | 87.24 | 78.35 |
| hsa-miR-335-5p | MIMAT0000765 | UCAAGAGCAAUAACGAAAAAUGU | 84.55 | 86.47 | 116.49 | 114.82 |
| hsa-miR-337-3p | MIMAT0000754 | CUCCUAUAUGAUGCCUUUCUUC | 91.12 | 101.94 | 113.33 | 110.74 |
| hsa-miR-337-5p | MIMAT0004695 | GAACGGCUUCAUACAGGAGUU | 127.73 | 109.53 | 102.13 | 111.81 |
| hsa-miR-338-3p | MIMAT0000763 | UCCAGCAUCAGUGAUUUUGUUG | 110.65 | 124.17 | 77.06 | 105.17 |
| hsa-miR-338-5p | MIMAT0004701 | AACAAUAUCCUGGUGCUGAGUG | 98.24 | 83.48 | 87.19 | 82.18 |
| hsa-miR-339-3p | MIMAT0004702 | UGAGCGCCUCGACGACAGAGCCG | 121.51 | 95.78 | 73.14 | 78.33 |

| | | | | | | |
|---|---|---|---|---|---|---|
| hsa-miR-339-5p | MIMAT0000764 | UCCCUGUCCUCCAGGAGCUCACG | 122.90 | 128.71 | 79.13 | 85.54 |
| hsa-miR-33a-3p | MIMAT0004506 | CAAUGUUUCCACAGUGCAUCAC | 72.94 | 78.40 | 80.46 | 70.81 |
| hsa-miR-33a-5p | MIMAT0000091 | GUGCAUUGUAGUUGCAUUGCA | 105.87 | 110.97 | 65.48 | 65.39 |
| hsa-miR-33b-3p | MIMAT0004811 | CAGUGCCUCGGCAGUGCAGCCC | 98.24 | 111.45 | 100.67 | 100.76 |
| hsa-miR-33b-5p | MIMAT0003301 | GUGCAUUGCUGUUGCAUUGC | 75.23 | 113.91 | 76.14 | 69.34 |
| hsa-miR-340-3p | MIMAT0000750 | UCCGUCUCAGUUACUUUAUAGC | 111.36 | 121.39 | 103.66 | 113.70 |
| hsa-miR-340-5p | MIMAT0004692 | UUAUAAAGCAAUGAGACUGAUU | 73.74 | 88.44 | 141.77 | 110.64 |
| hsa-miR-342-3p | MIMAT0000753 | UCUCACACAGAAAUCGCACCCGU | 118.09 | 100.43 | 96.21 | 91.04 |
| hsa-miR-342-5p | MIMAT0004694 | AGGGGUGCUAUCUGUGAUUGA | 98.70 | 79.78 | 106.67 | 90.93 |
| hsa-miR-345-5p | MIMAT0000772 | GCUGACUCCUAGUCCAGGGCUC | 108.92 | 87.21 | 81.18 | 82.90 |
| hsa-miR-346 | MIMAT0000773 | UGUCUGCCCGCAUGCCUGCCUCU | 115.52 | 110.67 | 102.48 | 116.48 |
| hsa-miR-34a-3p | MIMAT0004557 | CAAUCAGCAAGUAUACUGCCCU | 70.46 | 98.74 | 78.84 | 66.74 |
| hsa-miR-34a-5p | MIMAT0000255 | UGGCAGUGUCUUAGCUGGUUGU | 37.93 | 39.04 | 68.79 | 75.93 |
| hsa-miR-34b-3p | MIMAT0004676 | CAAUCACUAACUCCACUGCCAU | 67.80 | 70.55 | 98.20 | 104.07 |
| hsa-miR-34b-5p | MIMAT0000685 | UAGGCAGUGUCAUUAGCUGAUUG | 84.31 | 73.10 | 132.29 | 128.37 |
| hsa-miR-34c-3p | MIMAT0004677 | AAUCACUAACCACACGGCCAGG | 147.12 | 161.18 | 142.49 | 124.44 |
| hsa-miR-34c-5p | MIMAT0000686 | AGGCAGUGUAGUUAGCUGAUUGC | 33.46 | 39.89 | 80.23 | 91.66 |
| hsa-miR-3605-3p | MIMAT0017982 | CCUCCGUGUUACCUGUCCUCUAG | 97.59 | 97.07 | 105.37 | 102.44 |
| hsa-miR-3605-5p | MIMAT0017981 | UGAGGAUGGAUAGCAAGGAAGCC | 102.06 | 100.90 | 118.63 | 102.92 |
| hsa-miR-3606-5p | MIMAT0017983 | UUAGUGAAGGCUAUUUUAAUU | 100.09 | 88.94 | 83.16 | 91.98 |
| hsa-miR-3607-3p | MIMAT0017985 | ACUGUAAACGCUUUCUGAUG | 67.69 | 101.66 | 95.93 | 127.87 |
| hsa-miR-3607-5p | MIMAT0017984 | GCAUGUGAUGAAGCAAAUCAGU | 113.58 | 135.74 | 111.96 | 114.43 |
| hsa-miR-3609 | MIMAT0017986 | CAAAGUGAUGAGUAAUACUGGCUG | 106.30 | 109.48 | 109.49 | 105.28 |
| hsa-miR-3610 | MIMAT0017987 | GAAUCGGAAAGGAGGCGCCG | 125.39 | 151.82 | 88.27 | 77.49 |
| hsa-miR-3611 | MIMAT0017988 | UUGUGAAGAAAGAAAUUCUUA | 96.95 | 90.18 | 86.00 | 83.11 |
| hsa-miR-3612 | MIMAT0017989 | AGGAGGCAUCUUGAGAAAUGGA | 48.32 | 67.43 | 97.29 | 91.03 |
| hsa-miR-3613-3p | MIMAT0017991 | ACAAAAAAAAAAGCCCAACCCUUC | 101.42 | 92.48 | 106.73 | 98.81 |
| hsa-miR-3613-5p | MIMAT0017990 | UGUUGUACUUUUUUUUUUGUUC | 67.79 | 73.85 | 108.15 | 112.09 |
| hsa-miR-361-3p | MIMAT0004682 | UCCCCCAGGUGUGAUUCUGAUUU | 131.44 | 109.64 | 99.29 | 93.35 |
| hsa-miR-3614-3p | MIMAT0017993 | UAGCCUUCAGAUCUUGGUGUUUU | 127.90 | 118.96 | 113.37 | 114.52 |
| hsa-miR-3614-5p | MIMAT0017992 | CCACUUGGAUCUGAAGGCUGCCC | 120.40 | 128.01 | 87.39 | 110.00 |
| hsa-miR-3615 | MIMAT0017994 | UCUCUCGGCUCCUCGCGGCUC | 166.78 | 162.80 | 134.76 | 151.73 |
| hsa-miR-361-5p | MIMAT0000703 | UUAUCAGAAUCUCCAGGGGUAC | 91.08 | 102.49 | 97.84 | 97.84 |
| hsa-miR-3616-3p | MIMAT0017996 | CGAGGGCAUUUCAUGAUGCAGGC | 101.30 | 94.64 | 112.08 | 104.31 |
| hsa-miR-3616-5p | MIMAT0017995 | AUGAAGUGCACUCAUGAUAUGU | 132.55 | 145.68 | 88.55 | 80.55 |
| hsa-miR-3617-5p | MIMAT0017997 | AAAGACAUAGUUGCAAGAUGGG | 125.85 | 144.71 | 103.50 | 109.19 |
| hsa-miR-3618 | MIMAT0017998 | UGUCUACAUUAAUGAAAAGAGC | 119.16 | 137.73 | 110.67 | 127.90 |
| hsa-miR-3619-5p | MIMAT0017999 | UCAGCAGGCAGGCUGGUGCAGC | 98.13 | 121.46 | 67.47 | 76.42 |
| hsa-miR-3620-3p | MIMAT0018001 | UCACCCUGCAUCCCGCACCCAG | 113.57 | 126.62 | 100.18 | 112.83 |
| hsa-miR-3621 | MIMAT0018002 | CGCGGGUCGGGGUCUGCAGG | 110.44 | 87.21 | 96.87 | 92.24 |
| hsa-miR-3622a-3p | MIMAT0018004 | UCACCUGACCUCCCAUGCCUGU | 103.98 | 89.93 | 100.96 | 111.13 |

| | | | | | | |
|---|---|---|---|---|---|---|
| hsa-miR-3622a-5p | MIMAT0018003 | CAGGCACGGGAGCUCAGGUGAG | 102.83 | 99.83 | 106.68 | 100.42 |
| hsa-miR-3622b-3p | MIMAT0018006 | UCACCUGAGCUCCCGUGCCUG | 104.40 | 88.09 | 86.08 | 93.62 |
| hsa-miR-3622b-5p | MIMAT0018005 | AGGCAUGGGAGGUCAGGUGA | 37.17 | 55.14 | 69.57 | 70.45 |
| hsa-miR-362-3p | MIMAT0004683 | AACACACCUAUUCAAGGAUUCA | 98.59 | 86.38 | 84.42 | 90.57 |
| hsa-miR-362-5p | MIMAT0000705 | AAUCCUUGGAACCUAGGUGUGAGU | 70.92 | 76.43 | 103.66 | 79.10 |
| hsa-miR-363-3p | MIMAT0000707 | AAUUGCACGGUAUCCAUCUGUA | 129.28 | 84.59 | 104.27 | 130.04 |
| hsa-miR-363-5p | MIMAT0003385 | CGGGUGGAUCACGAUGCAAUUU | 36.49 | 37.31 | 100.66 | 89.33 |
| hsa-miR-3646 | MIMAT0018065 | AAAAUGAAAUGAGCCCAGCCCA | 116.58 | 67.71 | 90.67 | 87.29 |
| hsa-miR-3647-3p | MIMAT0018067 | AGAAAAUUUUUGUGUGUCUGAUC | 57.70 | 50.29 | 72.07 | 80.51 |
| hsa-miR-3647-5p | MIMAT0018066 | CUGAAGUGAUGAUUCACAUUCAU | 137.02 | 129.37 | 120.90 | 117.62 |
| hsa-miR-3648 | MIMAT0018068 | AGCCGCGGGGAUCGCCGAGGG | 131.01 | 93.15 | 102.76 | 102.36 |
| hsa-miR-3649 | MIMAT0018069 | AGGGACCUGAGUGUCUAAG | 135.51 | 134.47 | 104.27 | 105.38 |
| hsa-miR-365* | MIMAT0009199 | AGGGACUUUCAGGGGCAGCUGU | 83.51 | 75.96 | 108.04 | 95.95 |
| hsa-miR-3650 | MIMAT0018070 | AGGUGUGUCUGUAGAGUCC | 61.93 | 54.72 | 89.12 | 79.96 |
| hsa-miR-3651 | MIMAT0018071 | CAUAGCCCGGUCGCUGGUACAUGA | 71.53 | 68.86 | 124.78 | 117.51 |
| hsa-miR-3652 | MIMAT0018072 | CGGCUGGAGGUGUGAGGA | 88.64 | 81.25 | 135.29 | 139.64 |
| hsa-miR-3653 | MIMAT0018073 | CUAAGAAGUUGACUGAAG | 147.98 | 155.08 | 95.20 | 87.14 |
| hsa-miR-3654 | MIMAT0018074 | GACUGGACAAGCUGAGGAA | 112.21 | 137.75 | 76.50 | 101.08 |
| hsa-miR-3655 | MIMAT0018075 | GCUUGUCGCUGCGGUGUUGCU | 117.39 | 96.81 | 121.69 | 93.26 |
| hsa-miR-3656 | MIMAT0018076 | GGCGGGUGCGGGGGUGG | 127.86 | 102.34 | 120.02 | 113.59 |
| hsa-miR-3657 | MIMAT0018077 | UGUGUCCCAUUAUUGGUGAUU | 51.79 | 47.88 | 98.84 | 92.24 |
| hsa-miR-3658 | MIMAT0018078 | UUUAAGAAAACACCAUGGAGAU | 74.77 | 87.84 | 86.67 | 95.19 |
| hsa-miR-3659 | MIMAT0018080 | UGAGUGUUGUCUACGAGGGCA | 110.05 | 114.67 | 107.07 | 92.05 |
| hsa-miR-365a-3p | MIMAT0000710 | UAAUGCCCCUAAAAAUCCUUAU | 90.34 | 73.74 | 95.73 | 109.13 |
| hsa-miR-3660 | MIMAT0018081 | ACUGACAGGAGAGCAUUUUGA | 86.57 | 92.08 | 69.76 | 81.69 |
| hsa-miR-3661 | MIMAT0018082 | UGACCUGGGACUCGGACAGCUG | 112.92 | 116.71 | 100.96 | 82.14 |
| hsa-miR-3662 | MIMAT0018083 | GAAAAUGAUGAGUAGUGACUGAUG | 87.05 | 93.24 | 92.80 | 96.16 |
| hsa-miR-3663-3p | MIMAT0018085 | UGAGCACCACACAGGCCGGGCGC | 103.33 | 84.49 | 31.32 | 23.35 |
| hsa-miR-3663-5p | MIMAT0018084 | GCUGGUCUGCGUGGUGCUCGG | 94.66 | 112.02 | 92.89 | 89.09 |
| hsa-miR-3664-5p | MIMAT0018086 | AACUCUGUCUUCACUCAUGAGU | 108.69 | 82.95 | 108.92 | 96.92 |
| hsa-miR-3665 | MIMAT0018087 | AGCAGGUGCGGGGCGGCG | 3.89 | 19.52 | 98.40 | 85.53 |
| hsa-miR-3666 | MIMAT0018088 | CAGUGCAAGUGUAGAUGCCGA | 120.99 | 107.12 | 91.64 | 83.06 |
| hsa-miR-3667-3p | MIMAT0018090 | ACCUUCCUCUCCAUGGGUCUUU | 67.88 | 62.37 | 92.46 | 71.03 |
| hsa-miR-3667-5p | MIMAT0018089 | AAAGACCCAUUGAGGAGAAGGU | 113.73 | 119.79 | 80.66 | 94.20 |
| hsa-miR-3668 | MIMAT0018091 | AAUGUAGAGAUUGAUCAAAAU | 99.82 | 107.27 | 103.68 | 103.16 |
| hsa-miR-3669 | MIMAT0018092 | ACGGAAUAUGUAUACGGAAUAUA | 100.75 | 123.17 | 103.27 | 99.75 |
| hsa-miR-3670 | MIMAT0018093 | AGAGCUCACAGCUGUCCUUCUCUA | 107.40 | 105.03 | 89.03 | 93.02 |
| hsa-miR-3671 | MIMAT0018094 | AUCAAAUAAGGACUAGUCUGCA | 126.56 | 108.22 | 116.88 | 108.79 |
| hsa-miR-3672 | MIMAT0018095 | AUGAGACUCAUGUAAAACAUCUU | 109.40 | 132.11 | 114.06 | 124.81 |
| hsa-miR-3673 | MIMAT0018096 | AUGGAAUGUAUAUACGGAAUA | 112.48 | 112.18 | 81.18 | 86.62 |
| hsa-miR-367-3p | MIMAT0000719 | AAUUGCACUUUAGCAAUGGUGA | 51.75 | 52.20 | 95.89 | 102.36 |

| | | | | | | |
|---|---|---|---|---|---|---|
| hsa-miR-3674 | MIMAT0018097 | AUUGUAGAACCUAAGAUUGGCC | 57.75 | 59.08 | 88.17 | 80.59 |
| hsa-miR-3675-3p | MIMAT0018099 | CAUCUCUAAGGAACUCCCCCAA | 95.99 | 94.26 | 117.61 | 112.83 |
| hsa-miR-3675-5p | MIMAT0018098 | UAUGGGGCUUCUGUAGAGAUUUC | 124.45 | 136.85 | 105.37 | 109.44 |
| hsa-miR-367-5p | MIMAT0004686 | ACUGUUGCUAAUAUGCAACUCU | 96.08 | 93.38 | 85.57 | 63.36 |
| hsa-miR-3676-3p | MIMAT0018100 | CCGUGUUUCCCCCACGCUUU | 140.19 | 93.26 | 111.20 | 103.77 |
| hsa-miR-3677-3p | MIMAT0018101 | CUCGUGGGCUCUGGCCACGGCC | 102.38 | 102.94 | 98.92 | 100.02 |
| hsa-miR-3678-3p | MIMAT0018103 | CUGCAGAGUUUGUACGGACCGG | 71.47 | 87.49 | 87.95 | 81.58 |
| hsa-miR-3678-5p | MIMAT0018102 | UCCGUACAAACUCUGCUGUG | 106.21 | 104.72 | 95.52 | 84.56 |
| hsa-miR-3679-3p | MIMAT0018105 | CUUCCCCCCAGUAAUCUUCAUC | 120.91 | 118.50 | 95.52 | 100.75 |
| hsa-miR-3679-5p | MIMAT0018104 | UGAGGAUAUGGCAGGGAAGGGGA | 69.92 | 77.92 | 94.50 | 71.10 |
| hsa-miR-3680-3p | MIMAT0018107 | UUUUGCAUGACCCUGGGAGUAGG | 152.65 | 96.52 | 93.60 | 81.78 |
| hsa-miR-3680-5p | MIMAT0018106 | GACUCACUCACAGGAUUGUGCA | 94.07 | 104.21 | 113.19 | 100.02 |
| hsa-miR-3681-3p | MIMAT0018109 | ACACAGUGCUUCAUCCACUACU | 108.77 | 109.48 | 94.84 | 75.69 |
| hsa-miR-3681-5p | MIMAT0018108 | UAGUGGAUGAUGCACUCUGUGC | 68.54 | 68.32 | 114.40 | 106.98 |
| hsa-miR-3682-3p | MIMAT0018110 | UGAUGAUACAGGUGGAGGUAG | 93.63 | 121.93 | 96.72 | 118.81 |
| hsa-miR-3683 | MIMAT0018111 | UGCGACAUUGGAAGUAGUAUCA | 88.96 | 94.26 | 93.14 | 87.46 |
| hsa-miR-3684 | MIMAT0018112 | UUAGACCUAGUACACGUCCUU | 118.38 | 88.93 | 82.93 | 79.64 |
| hsa-miR-3685 | MIMAT0018113 | UUUCCUACCCUACCUGAAGACU | 102.06 | 101.66 | 90.42 | 114.52 |
| hsa-miR-3686 | MIMAT0018114 | AUCUGUAAGAGAAAGUAAAUGA | 80.02 | 85.08 | 80.56 | 93.74 |
| hsa-miR-3687 | MIMAT0018115 | CCCGGACAGGCGUUCGUGCGACGU | 112.73 | 119.13 | 118.45 | 110.54 |
| hsa-miR-3688-3p | MIMAT0018116 | UAUGGAAAGACUUUGCCACUCU | 138.10 | 121.79 | 103.49 | 118.05 |
| hsa-miR-3689a-3p | MIMAT0018118 | CUGGGAGGUGUGAUAUCGUGGU | 130.83 | 110.67 | 119.47 | 116.76 |
| hsa-miR-3689a-5p | MIMAT0018117 | UGUGAUAUCAUGGUUCCUGGGA | 93.69 | 125.81 | 100.31 | 119.88 |
| hsa-miR-3689b-3p | MIMAT0018181 | CUGGGAGGUGUGAUAUUGUGGU | 89.22 | 94.14 | 128.80 | 113.59 |
| hsa-miR-3689b-5p | MIMAT0018180 | UGUGAUAUCAUGGUUCCUGGGA | 101.30 | 100.88 | 97.34 | 114.05 |
| hsa-miR-3690 | MIMAT0018119 | ACCUGGACCCAGCGUAGACAAAG | 91.75 | 83.49 | 104.92 | 116.48 |
| hsa-miR-3691-5p | MIMAT0018120 | AGUGGAUGAUGGAGACUCGGUAC | 102.30 | 91.52 | 113.82 | 106.07 |
| hsa-miR-3692-3p | MIMAT0018122 | GUUCCACACUGACACUGCAGAAGU | 93.57 | 131.08 | 112.22 | 114.07 |
| hsa-miR-3692-5p | MIMAT0018121 | CCUGCUGGUCAGGAGUGGAUACUG | 137.84 | 140.69 | 106.39 | 111.86 |
| hsa-miR-369-3p | MIMAT0000721 | AAUAAUACAUGGUUGAUCUUU | 116.11 | 113.15 | 138.23 | 155.59 |
| hsa-miR-369-5p | MIMAT0001621 | AGAUCGACCGUGUUAUAUUCGC | 105.87 | 108.74 | 116.98 | 115.99 |
| hsa-miR-370 | MIMAT0000722 | GCCUGCUGGGGUGGAACCUGGU | 92.72 | 101.48 | 101.47 | 97.85 |
| hsa-miR-3713 | MIMAT0018164 | GGUAUCCGUUUGGGGAUGGU | 81.53 | 70.60 | 106.15 | 102.48 |
| hsa-miR-3714 | MIMAT0018165 | GAAGGCAGCAGUGCUCCCCUGU | 68.39 | 74.18 | 101.17 | 87.86 |
| hsa-miR-371a-3p | MIMAT0000723 | AAGUGCCGCCAUCUUUUGAGUGU | 135.14 | 135.97 | 129.29 | 128.84 |
| hsa-miR-371a-5p | MIMAT0004687 | ACUCAAACUGUGGGGGCACU | 139.61 | 137.14 | 101.10 | 110.15 |
| hsa-miR-372 | MIMAT0000724 | AAAGUGCUGCGACAUUUGAGCGU | 86.91 | 115.93 | 121.86 | 128.30 |
| hsa-miR-373-3p | MIMAT0000726 | GAAGUGCUUCGAUUUUGGGGUGU | 139.07 | 135.84 | 138.34 | 158.15 |
| hsa-miR-373-5p | MIMAT0000725 | ACUCAAAAUGGGGGCGCUUUCC | 151.73 | 161.56 | 111.33 | 92.81 |
| hsa-miR-374a-3p | MIMAT0004688 | CUUAUCAGAUUGUAUUGUAAUU | 80.02 | 97.14 | 82.37 | 97.39 |
| hsa-miR-374a-5p | MIMAT0000727 | UUAUAAUACAACCUGAUAAGUG | 87.82 | 96.20 | 114.49 | 84.95 |

| | | | | | | |
|---|---|---|---|---|---|---|
| hsa-miR-374b-3p | MIMAT0004956 | CUUAGCAGGUUGUAUUAUCAUU | 49.17 | 42.60 | 114.46 | 90.70 |
| hsa-miR-374b-5p | MIMAT0004955 | AUAUAAUACAACCUGCUAAGUG | 64.48 | 70.12 | 116.79 | 117.31 |
| hsa-miR-374c-5p | MIMAT0018443 | AUAAUACAACCUGCUAAGUGCU | 60.53 | 57.53 | 96.54 | 93.74 |
| hsa-miR-375 | MIMAT0000728 | UUUGUUCGUUCGGCUCGCGUGA | 115.48 | 101.41 | 97.56 | 93.74 |
| hsa-miR-376a-3p | MIMAT0000729 | AUCAUAGAGGAAAAUCCACGU | 77.76 | 102.14 | 94.06 | 84.77 |
| hsa-miR-376a-5p | MIMAT0003386 | GUAGAUUCUCCUUCUAUGAGUA | 104.10 | 94.86 | 84.07 | 96.59 |
| hsa-miR-376b-3p | MIMAT0002172 | AUCAUAGAGGAAAAUCCAUGUU | 111.64 | 104.67 | 102.79 | 87.26 |
| hsa-miR-376c-3p | MIMAT0000720 | AACAUAGAGGAAAUUCCACGU | 86.18 | 69.76 | 82.27 | 67.03 |
| hsa-miR-377-3p | MIMAT0000730 | AUCACACAAAGGCAACUUUUGU | 98.23 | 106.76 | 93.14 | 109.20 |
| hsa-miR-377-5p | MIMAT0004689 | AGAGGUUGCCCUUGGUGAAUUC | 57.65 | 56.03 | 76.09 | 69.78 |
| hsa-miR-378a-3p | MIMAT0000732 | ACUGGACUUGGAGUCAGAAGG | 105.43 | 116.51 | 111.95 | 112.54 |
| hsa-miR-378a-5p | MIMAT0000731 | CUCCUGACUCCAGGUCCUGUGU | 76.95 | 105.18 | 98.12 | 106.33 |
| hsa-miR-378b | MIMAT0014999 | ACUGGACUUGGAGGCAGAA | 135.51 | 142.06 | 125.60 | 133.26 |
| hsa-miR-378c | MIMAT0016847 | ACUGGACUUGGAGUCAGAAGAGUGG | 106.30 | 91.63 | 101.17 | 110.82 |
| hsa-miR-379-3p | MIMAT0004690 | UAUGUAACAUGGUCCACUAACU | 122.50 | 136.81 | 102.48 | 102.16 |
| hsa-miR-379-5p | MIMAT0000733 | UGGUAGACUAUGGAACGUAGG | 98.45 | 88.91 | 69.61 | 60.31 |
| hsa-miR-380-3p | MIMAT0000735 | UAUGUAAUAUGGUCCACAUCUU | 82.26 | 82.76 | 97.85 | 94.58 |
| hsa-miR-380-5p | MIMAT0000734 | UGGUUGACCAUAGAACAUGCGC | 65.49 | 58.77 | 134.01 | 130.53 |
| hsa-miR-381-3p | MIMAT0000736 | UAUACAAGGGCAAGCUCUCUGU | 104.62 | 91.26 | 126.82 | 110.46 |
| hsa-miR-382-5p | MIMAT0000737 | GAAGUUGUUCGUGGUGGAUUCG | 97.14 | 87.75 | 102.56 | 98.51 |
| hsa-miR-383 | MIMAT0000738 | AGAUCAGAAGGUGAUUGUGGCU | 49.96 | 61.66 | 125.15 | 132.29 |
| hsa-miR-384 | MIMAT0001075 | AUUCCUAGAAAUUGUUCAUA | 92.00 | 117.16 | 74.67 | 84.91 |
| hsa-miR-3907 | MIMAT0018179 | AGGUGCUCCAGGCUGGCUCACA | 49.80 | 49.98 | 81.21 | 88.26 |
| hsa-miR-3908 | MIMAT0018182 | GAGCAAUGUAGGUAGACUGUUU | 71.96 | 89.05 | 96.39 | 73.13 |
| hsa-miR-3909 | MIMAT0018183 | UGUCCUCUAGGGCCUGCAGUCU | 126.95 | 142.06 | 80.80 | 85.00 |
| hsa-miR-3910 | MIMAT0018184 | AAAGGCAUAAAACCAAGACA | 81.97 | 61.29 | 81.32 | 99.55 |
| hsa-miR-3911 | MIMAT0018185 | UGUGUGGAUCCUGGAGGAGGCA | 73.21 | 57.48 | 97.33 | 106.98 |
| hsa-miR-3912 | MIMAT0018186 | UAACGCAUAAUAUGGACAUGU | 96.86 | 118.84 | 86.55 | 116.34 |
| hsa-miR-3913-5p | MIMAT0018187 | UUUGGGACUGAUCUUGAUGUCU | 125.31 | 115.46 | 108.72 | 103.35 |
| hsa-miR-3914 | MIMAT0018188 | AAGGAACCAGAAAAUGAGAAGU | 119.95 | 112.63 | 105.04 | 120.32 |
| hsa-miR-3915 | MIMAT0018189 | UUGAGGAAAAGAUGGUCUUAUU | 97.89 | 101.57 | 93.08 | 90.55 |
| hsa-miR-3916 | MIMAT0018190 | AAGAGGAAGAAAUGGCUGGUUCUCAG | 74.50 | 90.71 | 106.23 | 122.56 |
| hsa-miR-3917 | MIMAT0018191 | GCUCGGACUGAGCAGGUGGG | 107.79 | 93.61 | 101.73 | 123.09 |
| hsa-miR-3918 | MIMAT0018192 | ACAGGGCCGCAGAUGGAGACU | 72.78 | 87.36 | 91.69 | 85.87 |
| hsa-miR-3919 | MIMAT0018193 | GCAGAGAACAAAGGACUCAGU | 78.10 | 71.66 | 86.92 | 107.97 |
| hsa-miR-3920 | MIMAT0018195 | ACUGAUUAUCUUAACUCUCUGA | 123.81 | 122.45 | 103.97 | 83.26 |
| hsa-miR-3921 | MIMAT0018196 | UCUCUGAGUACCAUAUGCCUUGU | 82.51 | 103.53 | 98.95 | 105.67 |
| hsa-miR-3922-3p | MIMAT0018197 | UCUGGCCUUGACUUGACUCUUU | 75.55 | 116.04 | 105.33 | 80.17 |
| hsa-miR-3923 | MIMAT0018198 | AACUAGUAAUGUUGGAUUAGGG | 86.15 | 79.32 | 98.84 | 102.15 |
| hsa-miR-3924 | MIMAT0018199 | AUAUGUAUAUGUGACUGCUACU | 107.57 | 88.52 | 92.16 | 84.36 |
| hsa-miR-3925-5p | MIMAT0018200 | AAGAGAACUGAAAGUGGAGCCU | 87.28 | 90.12 | 117.40 | 99.17 |

| | | | | | | |
|---|---|---|---|---|---|---|
| hsa-miR-3926 | MIMAT0018201 | UGGCCAAAAAGCAGGCAGAGA | 116.71 | 119.00 | 97.29 | 112.80 |
| hsa-miR-3927-3p | MIMAT0018202 | CAGGUAGAUAUUUGAUAGGCAU | 96.95 | 90.95 | 82.26 | 92.53 |
| hsa-miR-3928 | MIMAT0018205 | GGAGGAACCUUGGAGCUUCGGC | 83.26 | 101.94 | 101.73 | 94.20 |
| hsa-miR-3929 | MIMAT0018206 | GAGGCUGAUGUGAGUAGACCACU | 96.57 | 93.26 | 90.93 | 103.23 |
| hsa-miR-3934-5p | MIMAT0018349 | UCAGGUGUGGAAACUGAGGCAG | 95.89 | 99.17 | 109.49 | 105.67 |
| hsa-miR-3935 | MIMAT0018350 | UGUAGAUACGAGCACCAGCCAC | 67.87 | 59.83 | 77.45 | 69.89 |
| hsa-miR-3936 | MIMAT0018351 | UAAGGGGUGUAUGGCAGAUGCA | 83.79 | 96.86 | 84.61 | 106.26 |
| hsa-miR-3937 | MIMAT0018352 | ACAGGCGGCUGUAGCAAUGGGGG | 89.12 | 117.65 | 103.25 | 89.91 |
| hsa-miR-3938 | MIMAT0018353 | AAUUCCCUUGUAGAUAACCCGG | 94.07 | 116.71 | 90.08 | 90.60 |
| hsa-miR-3939 | MIMAT0018355 | UACGCGCAGACCACAGGAUGUC | 91.43 | 83.30 | 118.36 | 113.19 |
| hsa-miR-3940-3p | MIMAT0018356 | CAGCCCGGAUCCCAGCCCACUU | 131.99 | 111.31 | 103.03 | 123.37 |
| hsa-miR-3941 | MIMAT0018357 | UUACACACAACUGAGGAUCAUA | 79.70 | 87.38 | 81.24 | 81.18 |
| hsa-miR-3942-5p | MIMAT0018358 | AAGCAAUACUGUUACCUGAAAU | 86.68 | 94.90 | 93.44 | 87.25 |
| hsa-miR-3943 | MIMAT0018359 | UAGCCCCCAGGCUUCACUUGGCG | 115.78 | 91.67 | 114.53 | 113.65 |
| hsa-miR-3944-3p | MIMAT0018360 | UUCGGGCUGGCCUGCUGCUCCGG | 128.50 | 147.49 | 83.47 | 72.66 |
| hsa-miR-3945 | MIMAT0018361 | AGGGCAUAGGAGAGGGUUGAUAU | 105.11 | 91.67 | 99.33 | 117.54 |
| hsa-miR-409-3p | MIMAT0001639 | GAAUGUUGCUCGGUGAACCCCU | 89.12 | 94.64 | 68.91 | 79.01 |
| hsa-miR-409-5p | MIMAT0001638 | AGGUUACCCGAGCAACUUUGCAU | 101.74 | 98.34 | 91.44 | 94.22 |
| hsa-miR-410 | MIMAT0002171 | AAUAUAACACAGAUGGCCUGU | 86.25 | 78.13 | 120.83 | 119.59 |
| hsa-miR-411-3p | MIMAT0004813 | UAUGUAACACGGUCCACUAACC | 113.07 | 110.81 | 118.49 | 112.90 |
| hsa-miR-411-5p | MIMAT0003329 | UAGUAGACCGUAUAGCGUACG | 92.18 | 96.39 | 99.51 | 96.18 |
| hsa-miR-412 | MIMAT0002170 | ACUUCACCUGGUCCACUAGCCGU | 117.10 | 74.16 | 69.14 | 75.67 |
| hsa-miR-421 | MIMAT0003339 | AUCAACAGACAUUAAUUGGGCGC | 104.26 | 84.09 | 86.65 | 67.60 |
| hsa-miR-422a | MIMAT0001339 | ACUGGACUUAGGGUCAGAAGGC | 132.40 | 119.29 | 143.20 | 146.67 |
| hsa-miR-423-3p | MIMAT0001340 | AGCUCGGUCUGAGGCCCCUCAGU | 104.04 | 95.64 | 95.89 | 82.26 |
| hsa-miR-423-5p | MIMAT0004748 | UGAGGGGCAGAGAGCGAGACUUU | 89.15 | 82.78 | 126.58 | 115.28 |
| hsa-miR-424-3p | MIMAT0004749 | CAAAACGUGAGGCGCUGCUAU | 74.39 | 67.76 | 121.21 | 120.82 |
| hsa-miR-424-5p | MIMAT0001341 | CAGCAGCAAUUCAUGUUUUGAA | 81.40 | 85.91 | 86.20 | 83.08 |
| hsa-miR-4251 | MIMAT0016883 | CCUGAGAAAAGGGCCAA | 121.17 | 108.29 | 82.88 | 100.53 |
| hsa-miR-4252 | MIMAT0016886 | GGCCACUGAGUCAGCACCA | 89.70 | 83.49 | 101.60 | 112.83 |
| hsa-miR-4253 | MIMAT0016882 | AGGGCAUGUCCAGGGGGU | 63.86 | 50.97 | 96.80 | 80.17 |
| hsa-miR-425-3p | MIMAT0001343 | AUCGGGAAUGUCGUGUCCGCCC | 85.25 | 77.76 | 97.34 | 77.81 |
| hsa-miR-4254 | MIMAT0016884 | GCCUGGAGCUACUCCACCAUCUC | 88.79 | 68.32 | 81.87 | 81.24 |
| hsa-miR-4255 | MIMAT0016885 | CAGUGUUCAGAGAUGGA | 81.51 | 84.88 | 110.13 | 101.49 |
| hsa-miR-425-5p | MIMAT0003393 | AAUGACACGAUCACUCCCGUUGA | 50.61 | 74.16 | 92.31 | 101.25 |
| hsa-miR-4256 | MIMAT0016877 | AUCUGACCUGAUGAAGGU | 67.19 | 73.75 | 90.24 | 80.42 |
| hsa-miR-4257 | MIMAT0016878 | CCAGAGGUGGGGACUGAG | 114.60 | 102.85 | 103.96 | 87.77 |
| hsa-miR-4258 | MIMAT0016879 | CCCCGCCACCGCCUUGG | 92.18 | 86.47 | 87.87 | 102.51 |
| hsa-miR-4259 | MIMAT0016880 | CAGUUGGGUCUAGGGGUCAGGA | 88.45 | 67.99 | 122.59 | 119.47 |
| hsa-miR-4260 | MIMAT0016881 | CUUGGGGCAUGGAGUCCCA | 115.22 | 83.30 | 100.62 | 100.53 |
| hsa-miR-4261 | MIMAT0016890 | AGGAAACAGGGACCCA | 132.67 | 140.53 | 147.55 | 152.13 |

| | | | | | | |
|---|---|---|---|---|---|---|
| hsa-miR-4262 | MIMAT0016894 | GACAUUCAGACUACCUG | 102.38 | 119.01 | 90.08 | 92.77 |
| hsa-miR-4263 | MIMAT0016898 | AUUCUAAGUGCCUUGGCC | 100.80 | 119.86 | 94.67 | 89.41 |
| hsa-miR-4264 | MIMAT0016899 | ACUCAGUCAUGGUCAUU | 78.10 | 80.70 | 129.08 | 103.92 |
| hsa-miR-4265 | MIMAT0016891 | CUGUGGGCUCAGCUCUGGG | 51.79 | 67.17 | 116.49 | 117.54 |
| hsa-miR-4266 | MIMAT0016892 | CUAGGAGGCCUUGGCC | 86.83 | 71.63 | 108.65 | 96.14 |
| hsa-miR-4267 | MIMAT0016893 | UCCAGCUCGGUGGCAC | 93.69 | 89.44 | 69.41 | 76.29 |
| hsa-miR-4268 | MIMAT0016896 | GGCUCCUCCUCUCAGGAUGUG | 109.22 | 95.01 | 82.76 | 85.15 |
| hsa-miR-4269 | MIMAT0016897 | GCAGGCACAGACAGCCCUGGC | 99.78 | 102.06 | 84.61 | 105.09 |
| hsa-miR-4270 | MIMAT0016900 | UCAGGGAGUCAGGGGAGGGC | 89.70 | 92.53 | 115.34 | 103.92 |
| hsa-miR-4271 | MIMAT0016901 | GGGGGAAGAAAAGGUGGGG | 84.61 | 63.63 | 70.70 | 65.87 |
| hsa-miR-4272 | MIMAT0016902 | CAUUCAACUAGUGAUUGU | 108.92 | 68.53 | 94.99 | 75.29 |
| hsa-miR-4273 | MIMAT0016903 | GUGUUCUCUGAUGGACAG | 100.35 | 95.60 | 113.92 | 106.07 |
| hsa-miR-4274 | MIMAT0016906 | CAGCAGUCCCUCCCCCUG | 103.59 | 104.29 | 80.69 | 70.84 |
| hsa-miR-4275 | MIMAT0016905 | CCAAUUACCACUUCUUU | 94.31 | 100.27 | 97.76 | 108.31 |
| hsa-miR-4276 | MIMAT0016904 | CUCAGUGACUCAUGUGC | 63.86 | 66.15 | 79.73 | 75.88 |
| hsa-miR-4277 | MIMAT0016908 | GCAGUUCUGAGCACAGUACAC | 87.95 | 84.97 | 88.38 | 98.17 |
| hsa-miR-4278 | MIMAT0016910 | CUAGGGGGUUUGCCCUUG | 80.76 | 68.86 | 89.34 | 94.96 |
| hsa-miR-4279 | MIMAT0016909 | CUCUCCUCCCGGCUUC | 145.42 | 133.13 | 129.58 | 146.11 |
| hsa-miR-4280 | MIMAT0016911 | GAGUGUAGUUCUGAGCAGAGC | 111.97 | 119.13 | 102.27 | 114.43 |
| hsa-miR-4281 | MIMAT0016907 | GGGUCCCGGGGAGGGGGG | 109.68 | 108.00 | 115.02 | 107.81 |
| hsa-miR-4282 | MIMAT0016912 | UAAAAUUUGCAUCCAGGA | 100.64 | 100.32 | 78.00 | 96.12 |
| hsa-miR-4283 | MIMAT0016914 | UGGGGCUCAGCGAGUUU | 109.27 | 113.05 | 90.08 | 106.86 |
| hsa-miR-4284 | MIMAT0016915 | GGGCUCACAUCACCCCAU | 116.91 | 115.06 | 111.99 | 132.55 |
| hsa-miR-4285 | MIMAT0016913 | GCGGCGAGUCCGACUCAU | 104.07 | 100.75 | 117.25 | 113.19 |
| hsa-miR-4286 | MIMAT0016916 | ACCCCACUCCUGGUACC | 145.18 | 148.60 | 124.41 | 100.02 |
| hsa-miR-4287 | MIMAT0016917 | UCUCCCUUGAGGGCACUUU | 93.46 | 104.11 | 99.47 | 101.62 |
| hsa-miR-4288 | MIMAT0016918 | UUGUCUGCUGAGUUUCC | 72.19 | 80.86 | 73.85 | 82.87 |
| hsa-miR-4289 | MIMAT0016920 | GCAUUGUGCAGGGCUAUCA | 116.75 | 111.36 | 95.52 | 109.93 |
| hsa-miR-429 | MIMAT0001536 | UAAUACUGUCUGGUAAAACCGU | 55.60 | 50.10 | 77.25 | 54.10 |
| hsa-miR-4290 | MIMAT0016921 | UGCCCUCCUUUCUUCCCUC | 66.51 | 91.83 | 103.97 | 117.29 |
| hsa-miR-4291 | MIMAT0016922 | UUCAGCAGGAACAGCU | 78.45 | 92.41 | 84.12 | 106.26 |
| hsa-miR-4292 | MIMAT0016919 | CCCCUGGGCCGGCCUUGG | 130.76 | 155.15 | 110.90 | 126.01 |
| hsa-miR-4293 | MIMAT0016848 | CAGCCUGACAGGAACAG | 67.87 | 68.18 | 69.39 | 75.56 |
| hsa-miR-4294 | MIMAT0016849 | GGGAGUCUACAGCAGGG | 125.63 | 113.84 | 118.91 | 87.86 |
| hsa-miR-4295 | MIMAT0016844 | CAGUGCAAUGUUUUCCUU | 137.45 | 114.79 | 123.87 | 97.44 |
| hsa-miR-4296 | MIMAT0016845 | AUGUGGGCUCAGGCUCA | 75.55 | 66.15 | 105.33 | 109.67 |
| hsa-miR-4297 | MIMAT0016846 | UGCCUUCCUGUCUGUG | 121.08 | 86.27 | 108.24 | 137.95 |
| hsa-miR-4298 | MIMAT0016852 | CUGGGACAGGAGGAGGAGGCAG | 126.37 | 110.67 | 131.11 | 123.09 |
| hsa-miR-4299 | MIMAT0016851 | GCUGGUGACAUGAGAGGC | 121.17 | 129.71 | 93.41 | 112.01 |
| hsa-miR-4300 | MIMAT0016853 | UGGGAGCUGGACUACUUC | 110.02 | 102.74 | 102.83 | 89.05 |
| hsa-miR-4301 | MIMAT0016850 | UCCCACUACUUCACUUGUGA | 104.07 | 115.83 | 112.81 | 98.95 |

| | | | | | | |
|---|---|---|---|---|---|---|
| hsa-miR-4302 | MIMAT0016855 | CCAGUGUGGCUCAGCGAG | 105.21 | 123.41 | 66.19 | 84.93 |
| hsa-miR-4303 | MIMAT0016856 | UUCUGAGCUGAGGACAG | 80.28 | 89.25 | 31.88 | 36.02 |
| hsa-miR-4304 | MIMAT0016854 | CCGGCAUGUCCAGGGCA | 104.02 | 94.62 | 105.87 | 115.26 |
| hsa-miR-4305 | MIMAT0016857 | CCUAGACACCUCCAGUUC | 69.24 | 86.27 | 99.71 | 82.85 |
| hsa-miR-4306 | MIMAT0016858 | UGGAGAGAAAGGCAGUA | 81.00 | 62.90 | 92.53 | 100.55 |
| hsa-miR-4307 | MIMAT0016860 | AAUGUUUUUUCCUGUUUCC | 99.34 | 79.64 | 99.25 | 90.75 |
| hsa-miR-4308 | MIMAT0016861 | UCCCUGGAGUUUCUUCUU | 88.96 | 94.77 | 111.49 | 90.36 |
| hsa-miR-4309 | MIMAT0016859 | CUGGAGUCUAGGAUUCCA | 86.23 | 83.70 | 101.73 | 94.59 |
| hsa-miR-4310 | MIMAT0016862 | GCAGCAUUCAUGUCCC | 107.48 | 82.42 | 111.73 | 99.48 |
| hsa-miR-4311 | MIMAT0016863 | GAAAGAGAGCUGAGUGUG | 105.13 | 105.57 | 111.09 | 111.17 |
| hsa-miR-4312 | MIMAT0016864 | GGCCUUGUUCCUGUCCCCA | 66.28 | 62.63 | 104.02 | 92.29 |
| hsa-miR-4313 | MIMAT0016865 | AGCCCCCUGGCCCCAAACCC | 122.86 | 124.31 | 115.82 | 109.18 |
| hsa-miR-431-3p | MIMAT0004757 | CAGGUCGUCUUGCAGGGCUUCU | 79.00 | 65.89 | 112.33 | 117.31 |
| hsa-miR-4314 | MIMAT0016868 | CUCUGGGAAAUGGGACAG | 102.07 | 105.03 | 74.31 | 75.51 |
| hsa-miR-4315 | MIMAT0016866 | CCGCUUUCUGAGCUGGAC | 93.75 | 95.28 | 117.61 | 85.53 |
| hsa-miR-431-5p | MIMAT0001625 | UGUCUUGCAGGCCGUCAUGCA | 74.01 | 53.57 | 98.76 | 84.88 |
| hsa-miR-4316 | MIMAT0016867 | GGUGAGGCUAGCUGGUG | 29.73 | 76.16 | 93.96 | 81.14 |
| hsa-miR-4317 | MIMAT0016872 | ACAUUGCCAGGGAGUUU | 95.23 | 112.38 | 100.09 | 95.03 |
| hsa-miR-4318 | MIMAT0016869 | CACUGUGGGUACAUGCU | 114.25 | 120.62 | 84.61 | 93.80 |
| hsa-miR-4319 | MIMAT0016870 | UCCCUGAGCAAAGCCAC | 89.02 | 84.88 | 96.87 | 97.03 |
| hsa-miR-4320 | MIMAT0016871 | GGGAUUCUGUAGCUUCCU | 60.75 | 79.16 | 82.40 | 84.46 |
| hsa-miR-4321 | MIMAT0016874 | UUAGCGGUGGACCGCCCUGCG | 116.14 | 112.44 | 103.96 | 132.55 |
| hsa-miR-4322 | MIMAT0016873 | CUGUGGGCUCAGCGCGUGGGG | 103.84 | 95.01 | 109.07 | 99.86 |
| hsa-miR-4323 | MIMAT0016875 | CAGCCCCACAGCCUCAGA | 93.46 | 80.25 | 120.27 | 118.78 |
| hsa-miR-432-3p | MIMAT0002815 | CUGGAUGGCUCCUCCAUGUCU | 119.74 | 129.95 | 127.80 | 98.41 |
| hsa-miR-4324 | MIMAT0016876 | CCCUGAGACCCUAACCUUAA | 119.99 | 121.16 | 102.49 | 100.51 |
| hsa-miR-4325 | MIMAT0016887 | UUGCACUUGUCUCAGUGA | 81.50 | 93.15 | 96.38 | 112.87 |
| hsa-miR-432-5p | MIMAT0002814 | UCUUGGAGUAGGUCAUUGGGUGG | 137.46 | 138.43 | 101.47 | 88.82 |
| hsa-miR-4326 | MIMAT0016888 | UGUUCCUCUGUCUCCCAGAC | 42.06 | 47.72 | 92.00 | 84.46 |
| hsa-miR-4327 | MIMAT0016889 | GGCUUGCAUGGGGGACUGG | 130.76 | 110.70 | 94.46 | 79.92 |
| hsa-miR-4328 | MIMAT0016926 | CCAGUUUUCCCAGGAUU | 95.66 | 98.78 | 72.21 | 87.93 |
| hsa-miR-4329 | MIMAT0016923 | CCUGAGACCCUAGUUCCAC | 127.11 | 119.40 | 78.44 | 87.47 |
| hsa-miR-433 | MIMAT0001627 | AUCAUGAUGGGCUCCUCGGUGU | 113.56 | 98.09 | 89.06 | 98.09 |
| hsa-miR-4330 | MIMAT0016924 | CCUCAGAUCAGAGCCUUGC | 81.77 | 60.69 | 91.75 | 103.70 |
| hsa-miR-448 | MIMAT0001532 | UUGCAUAUGUAGGAUGUCCCAU | 99.73 | 111.45 | 114.07 | 110.13 |
| hsa-miR-449a | MIMAT0001541 | UGGCAGUGUAUUGUUAGCUGGU | 21.82 | 27.24 | 45.59 | 62.93 |
| hsa-miR-449b-3p | MIMAT0009203 | CAGCCACAACUACCCUGCCACU | 103.77 | 94.79 | 131.84 | 107.89 |
| hsa-miR-449b-5p | MIMAT0003327 | AGGCAGUGUAUUGUUAGCUGGC | 36.17 | 26.71 | 65.57 | 60.88 |
| hsa-miR-449c-3p | MIMAT0013771 | UUGCUAGUUGCACUCCUCUCUGU | 145.70 | 142.40 | 101.73 | 105.67 |
| hsa-miR-449c-5p | MIMAT0010251 | UAGGCAGUGUAUUGCUAGCGGCUGU | 91.29 | 66.05 | 94.37 | 84.42 |
| hsa-miR-450a-5p | MIMAT0001545 | UUUUGCGAUGUGUUCCUAAUAU | 169.53 | 177.24 | 156.37 | 178.96 |

| | | | | | | |
|---|---|---|---|---|---|---|
| hsa-miR-450b-3p | MIMAT0004910 | UUGGGAUCAUUUUGCAUCCAUA | 95.41 | 92.40 | 98.49 | 107.28 |
| hsa-miR-450b-5p | MIMAT0004909 | UUUUGCAAUAUGUUCCUGAAUA | 36.30 | 34.18 | 86.13 | 94.62 |
| hsa-miR-451a | MIMAT0001631 | AAACCGUUACCAUUACUGAGUU | 80.71 | 89.15 | 96.72 | 133.15 |
| hsa-miR-452-3p | MIMAT0001636 | CUCAUCUGCAAAGAAGUAAGUG | 86.91 | 92.39 | 101.40 | 84.50 |
| hsa-miR-452-5p | MIMAT0001635 | AACUGUUUGCAGAGGAAACUGA | 136.07 | 110.20 | 85.24 | 87.48 |
| hsa-miR-453 | MIMAT0001630 | AGGUUGUCCGUGGUGAGUUCGCA | 102.07 | 99.83 | 116.00 | 128.83 |
| hsa-miR-454-3p | MIMAT0003885 | UAGUGCAAUAUUGCUUAUAGGGU | 133.80 | 134.07 | 102.83 | 106.86 |
| hsa-miR-454-5p | MIMAT0003884 | ACCCUAUCAAUAUUGUCUCUGC | 129.91 | 129.36 | 143.18 | 158.56 |
| hsa-miR-455-3p | MIMAT0004784 | GCAGUCCAUGGGCAUAUACAC | 129.22 | 126.39 | 86.78 | 115.88 |
| hsa-miR-455-5p | MIMAT0003150 | UAUGUGCCUUUGGACUACAUCG | 94.45 | 75.34 | 126.30 | 112.87 |
| hsa-miR-466 | MIMAT0015002 | AUACACAUACACGCAACACACAU | 102.26 | 111.18 | 87.04 | 88.59 |
| hsa-miR-483-3p | MIMAT0002173 | UCACUCCUCUCCUCCCGUCUU | 128.29 | 126.11 | 96.63 | 95.38 |
| hsa-miR-483-5p | MIMAT0004761 | AAGACGGGAGGAAAGAAGGGAG | 87.08 | 90.20 | 119.87 | 107.38 |
| hsa-miR-484 | MIMAT0002174 | UCAGGCUCAGUCCCCUCCCGAU | 150.67 | 155.24 | 100.29 | 108.68 |
| hsa-miR-485-3p | MIMAT0002176 | GUCAUACACGGCUCUCCUCUCU | 117.43 | 134.25 | 94.49 | 131.44 |
| hsa-miR-485-5p | MIMAT0002175 | AGAGGCUGGCCGUGAUGAAUUC | 68.03 | 76.05 | 111.06 | 125.66 |
| hsa-miR-486-3p | MIMAT0004762 | CGGGGCAGCUCAGUACAGGAU | 96.58 | 87.82 | 111.05 | 115.17 |
| hsa-miR-486-5p | MIMAT0002177 | UCCUGUACUGAGCUGCCCCGAG | 157.32 | 116.04 | 109.60 | 104.30 |
| hsa-miR-487a | MIMAT0002178 | AAUCAUACAGGGACAUCCAGUU | 101.88 | 117.04 | 95.51 | 92.10 |
| hsa-miR-487b | MIMAT0003180 | AAUCGUACAGGGUCAUCCACUU | 137.84 | 139.42 | 114.89 | 120.07 |
| hsa-miR-488-3p | MIMAT0004763 | UUGAAAGGCUAUUUCUUGGUC | 109.93 | 95.32 | 102.79 | 110.63 |
| hsa-miR-488-5p | MIMAT0002804 | CCCAGAUAAUGGCACUCUCAA | 86.25 | 100.83 | 108.17 | 88.82 |
| hsa-miR-489 | MIMAT0002805 | GUGACAUCACAUAUACGGCAGC | 115.96 | 87.66 | 107.82 | 93.01 |
| hsa-miR-490-3p | MIMAT0002806 | CAACCUGGAGGACUCCAUGCUG | 120.76 | 107.21 | 106.88 | 117.51 |
| hsa-miR-490-5p | MIMAT0004764 | CCAUGGAUCUCCAGGUGGGU | 57.96 | 61.77 | 101.37 | 97.88 |
| hsa-miR-491-3p | MIMAT0004765 | CUUAUGCAAGAUUCCCUUCUAC | 159.71 | 132.00 | 88.08 | 101.90 |
| hsa-miR-491-5p | MIMAT0002807 | AGUGGGGAACCCUUCCAUGAGG | 117.07 | 88.40 | 102.32 | 99.78 |
| hsa-miR-492 | MIMAT0002812 | AGGACCUGCGGGACAAGAUUCUU | 92.16 | 105.77 | 95.89 | 109.37 |
| hsa-miR-493-3p | MIMAT0003161 | UGAAGGUCUACUGUGUGCCAGG | 153.63 | 201.98 | 117.11 | 123.28 |
| hsa-miR-493-5p | MIMAT0002813 | UUGUACAUGGUAGGCUUUCAUU | 101.74 | 92.99 | 109.45 | 100.99 |
| hsa-miR-494 | MIMAT0002816 | UGAAACAUACACGGGAAACCUC | 82.37 | 70.77 | 137.80 | 136.08 |
| hsa-miR-495-3p | MIMAT0002817 | AAACAAACAUGGUGCACUUCUU | 92.70 | 86.00 | 102.44 | 81.54 |
| hsa-miR-496 | MIMAT0002818 | UGAGUAUUACAUGGCCAAUCUC | 105.48 | 110.99 | 120.69 | 107.03 |
| hsa-miR-497-3p | MIMAT0004768 | CAAACCACACUGUGGUGUUAGA | 91.66 | 85.42 | 112.75 | 72.08 |
| hsa-miR-497-5p | MIMAT0002820 | CAGCAGCACACUGUGGUUUGU | 63.98 | 64.95 | 70.39 | 75.90 |
| hsa-miR-498 | MIMAT0002824 | UUUCAAGCCAGGGGGCGUUUUUC | 62.30 | 70.60 | 96.28 | 109.01 |
| hsa-miR-499a-3p | MIMAT0004772 | AACAUCACAGCAAGUCUGUGCU | 89.25 | 69.32 | 117.05 | 97.13 |
| hsa-miR-499a-5p | MIMAT0002870 | UUAAGACUUGCAGUGAUGUUU | 119.13 | 111.85 | 85.46 | 88.41 |
| hsa-miR-500a-3p | MIMAT0002871 | AUGCACCUGGGCAAGGAUUCUG | 95.38 | 101.98 | 98.84 | 119.47 |
| hsa-miR-500a-5p | MIMAT0004773 | UAAUCCUUGCUACCUGGGUGAGA | 177.47 | 179.25 | 120.91 | 117.15 |
| hsa-miR-500b | MIMAT0016925 | AAUCCUUGCUACCUGGGU | 90.69 | 104.32 | 97.29 | 99.74 |

| | | | | | | |
|---|---|---|---|---|---|---|
| hsa-miR-501-3p | MIMAT0004774 | AAUGCACCCGGGCAAGGAUUCU | 160.99 | 168.08 | 89.47 | 97.61 |
| hsa-miR-501-5p | MIMAT0002872 | AAUCCUUUGUCCCUGGGUGAGA | 113.18 | 101.18 | 86.50 | 92.84 |
| hsa-miR-502-3p | MIMAT0004775 | AAUGCACCUGGGCAAGGAUUCA | 85.83 | 87.74 | 120.13 | 121.16 |
| hsa-miR-502-5p | MIMAT0002873 | AUCCUUGCUAUCUGGGUGCUA | 82.88 | 82.25 | 77.36 | 80.68 |
| hsa-miR-503-5p | MIMAT0002874 | UAGCAGCGGGAACAGUUCUGCAG | 57.78 | 71.44 | 67.78 | 77.43 |
| hsa-miR-504 | MIMAT0002875 | AGACCCUGGUCUGCACUCUAUC | 83.53 | 86.10 | 105.04 | 95.19 |
| hsa-miR-505-3p | MIMAT0002876 | CGUCAACACUUGCUGGUUUCCU | 108.66 | 99.59 | 115.06 | 121.02 |
| hsa-miR-505-5p | MIMAT0004776 | GGGAGCCAGGAAGUAUUGAUGU | 89.48 | 77.48 | 111.52 | 103.18 |
| hsa-miR-506-3p | MIMAT0002878 | UAAGGCACCCUUCUGAGUAGA | 133.97 | 129.18 | 143.52 | 148.45 |
| hsa-miR-507 | MIMAT0002879 | UUUUGCACCUUUUGGAGUGAA | 58.04 | 68.11 | 78.87 | 76.11 |
| hsa-miR-508-3p | MIMAT0002880 | UGAUUGUAGCCUUUUGGAGUAGA | 102.10 | 144.70 | 109.86 | 74.53 |
| hsa-miR-508-5p | MIMAT0004778 | UACUCCAGAGGGCGUCACUCAUG | 32.03 | 46.16 | 75.53 | 59.94 |
| hsa-miR-509-3-5p | MIMAT0004975 | UACUGCAGACGUGGCAAUCAUG | 71.69 | 63.10 | 134.57 | 107.57 |
| hsa-miR-509-3p | MIMAT0002881 | UGAUUGGUACGUCUGUGGGUAG | 111.32 | 105.74 | 114.21 | 114.76 |
| hsa-miR-509-5p | MIMAT0004779 | UACUGCAGACAGUGGCAAUCA | 35.04 | 32.29 | 129.14 | 130.42 |
| hsa-miR-510 | MIMAT0002882 | UACUCAGGAGAGUGGCAAUCAC | 36.15 | 43.58 | 87.51 | 115.88 |
| hsa-miR-511 | MIMAT0002808 | GUGUCUUUUGCUCUGCAGUCA | 139.11 | 125.92 | 116.77 | 97.31 |
| hsa-miR-512-3p | MIMAT0002823 | AAGUGCUGUCAUAGCUGAGGUC | 156.83 | 167.92 | 95.63 | 99.30 |
| hsa-miR-512-5p | MIMAT0002822 | CACUCAGCCUUGAGGGCACUUUC | 116.50 | 85.27 | 76.32 | 59.02 |
| hsa-miR-513a-3p | MIMAT0004777 | UAAAUUUCACCUUUCUGAGAAGG | 121.07 | 100.43 | 107.66 | 69.81 |
| hsa-miR-513a-5p | MIMAT0002877 | UUCACAGGGAGGUGUCAU | 73.99 | 79.16 | 85.07 | 83.39 |
| hsa-miR-513b | MIMAT0005788 | UUCACAAGGAGGUGUCAUUUAU | 82.38 | 104.36 | 75.16 | 98.36 |
| hsa-miR-513c-5p | MIMAT0005789 | UUCUCAAGGAGGUGUCGUUUAU | 118.20 | 108.17 | 129.61 | 129.02 |
| hsa-miR-514a-3p | MIMAT0002883 | AUUGACACUUCUGUGAGUAGA | 71.42 | 66.15 | 87.06 | 76.14 |
| hsa-miR-514b-3p | MIMAT0015088 | AUUGACACCUCUGUGAGUGGA | 89.12 | 99.83 | 82.65 | 87.18 |
| hsa-miR-514b-5p | MIMAT0015087 | UUCUCAAGAGGGAGGCAAUCAU | 136.03 | 112.65 | 109.49 | 110.03 |
| hsa-miR-515-3p | MIMAT0002827 | GAGUGCCUUCUUUUGGAGCGUU | 174.45 | 173.51 | 119.73 | 118.78 |
| hsa-miR-515-5p | MIMAT0002826 | UUCUCCAAAAGAAAGCACUUUCUG | 120.57 | 88.57 | 70.82 | 104.45 |
| hsa-miR-516a-3p | MIMAT0002860 | UGCUUCCUUUCAGAGGGU | 98.63 | 99.52 | 94.91 | 113.53 |
| hsa-miR-516a-5p | MIMAT0004770 | UUCUCGAGGAAAGAAGCACUUUC | 121.17 | 102.34 | 112.26 | 97.36 |
| hsa-miR-516b-5p | MIMAT0002859 | AUCUGGAGGUAAGAAGCACUUU | 118.46 | 94.27 | 75.95 | 67.73 |
| hsa-miR-517-5p | MIMAT0002851 | CCUCUAGAUGGAAGCACUGUCU | 122.36 | 98.88 | 103.33 | 107.69 |
| hsa-miR-517a-3p | MIMAT0002852 | AUCGUGCAUCCCUUUAGAGUGU | 76.44 | 99.01 | 61.60 | 42.38 |
| hsa-miR-517b | MIMAT0002857 | UCGUGCAUCCCUUUAGAGUGUU | 150.89 | 144.26 | 107.34 | 122.39 |
| hsa-miR-517c-3p | MIMAT0002866 | AUCGUGCAUCCUUUUAGAGUGU | 87.25 | 85.64 | 128.09 | 105.51 |
| hsa-miR-518a-3p | MIMAT0002863 | GAAAGCGCUUCCCUUUGCUGGA | 99.79 | 85.50 | 104.04 | 90.63 |
| hsa-miR-518a-5p | MIMAT0005457 | CUGCAAAGGGAAGCCCUUUC | 22.37 | 17.67 | 59.23 | 41.25 |
| hsa-miR-518b | MIMAT0002844 | CAAAGCGCUCCCCUUUAGAGGU | 109.27 | 89.65 | 85.65 | 92.22 |
| hsa-miR-518c-3p | MIMAT0002848 | CAAAGCGCUUCUCUUUAGAGUGU | 101.10 | 133.28 | 102.83 | 108.05 |
| hsa-miR-518c-5p | MIMAT0002847 | UCUCUGGAGGGAAGCACUUUCUG | 107.03 | 91.11 | 95.78 | 97.19 |
| hsa-miR-518d-3p | MIMAT0002864 | CAAAGCGCUUCCCUUUGGAGC | 84.89 | 113.87 | 98.93 | 67.30 |

| | | | | | | |
|---|---|---|---|---|---|---|
| hsa-miR-518d-5p | MIMAT0005456 | CUCUAGAGGGAAGCACUUUCUG | 144.45 | 145.19 | 120.18 | 127.57 |
| hsa-miR-518e-3p | MIMAT0002861 | AAAGCGCUUCCCUUCAGAGUG | 116.49 | 98.12 | 118.83 | 117.92 |
| hsa-miR-518e-5p | MIMAT0005450 | CUCUAGAGGGAAGCGCUUUCUG | 92.67 | 76.46 | 82.22 | 112.60 |
| hsa-miR-518f-3p | MIMAT0002842 | GAAAGCGCUUCUCUUUAGAGG | 100.65 | 89.88 | 80.88 | 113.61 |
| hsa-miR-518f-5p | MIMAT0002841 | CUCUAGAGGGAAGCACUUUCUC | 106.81 | 113.45 | 85.45 | 87.29 |
| hsa-miR-519a-3p | MIMAT0002869 | AAAGUGCAUCCUUUUAGAGUGU | 125.97 | 131.43 | 137.97 | 161.83 |
| hsa-miR-519b-3p | MIMAT0002837 | AAAGUGCAUCCUUUUAGAGGUU | 134.15 | 120.33 | 115.46 | 124.00 |
| hsa-miR-519c-3p | MIMAT0002832 | AAAGUGCAUCUUUUUAGAGGAU | 113.20 | 121.58 | 114.13 | 116.31 |
| hsa-miR-519d | MIMAT0002853 | CAAAGUGCCUCCCUUUAGAGUG | 80.76 | 86.29 | 124.42 | 114.57 |
| hsa-miR-519e-3p | MIMAT0002829 | AAGUGCCUCCUUUUAGAGUGUU | 99.19 | 122.33 | 82.26 | 76.10 |
| hsa-miR-519e-5p | MIMAT0002828 | UUCUCCAAAAGGGAGCACUUUC | 108.53 | 106.16 | 68.25 | 73.52 |
| hsa-miR-520a-3p | MIMAT0002834 | AAAGUGCUUCCCUUUGGACUGU | 137.73 | 136.43 | 138.20 | 151.71 |
| hsa-miR-520a-5p | MIMAT0002833 | CUCCAGAGGGAAGUACUUUCU | 37.98 | 45.06 | 83.22 | 78.56 |
| hsa-miR-520b | MIMAT0002843 | AAAGUGCUUCCUUUUAGAGGG | 97.08 | 118.93 | 112.27 | 134.03 |
| hsa-miR-520c-3p | MIMAT0002846 | AAAGUGCUUCCUUUUAGAGGGU | 106.85 | 119.01 | 122.03 | 103.16 |
| hsa-miR-520d-3p | MIMAT0002856 | AAAGUGCUUCUCUUUGGUGGGU | 166.82 | 203.57 | 132.14 | 136.45 |
| hsa-miR-520d-5p | MIMAT0002855 | CUACAAAGGGAAGCCCUUUC | 162.63 | 147.31 | 104.33 | 87.81 |
| hsa-miR-520e | MIMAT0002825 | AAAGUGCUUCCUUUUUGAGGG | 100.08 | 101.57 | 94.08 | 123.82 |
| hsa-miR-520f | MIMAT0002830 | AAGUGCUUCCUUUUAGAGGGUU | 165.34 | 152.49 | 164.35 | 140.85 |
| hsa-miR-520g | MIMAT0002858 | ACAAAGUGCUUCCCUUUAGAGUGU | 125.60 | 107.31 | 114.13 | 118.77 |
| hsa-miR-520h | MIMAT0002867 | ACAAAGUGCUUCCCUUUAGAGU | 78.13 | 84.50 | 89.97 | 88.43 |
| hsa-miR-521 | MIMAT0002854 | AACGCACUUCCCUUUAGAGUGU | 101.53 | 90.65 | 78.01 | 102.80 |
| hsa-miR-522-3p | MIMAT0002868 | AAAAUGGUUCCCUUUAGAGUGU | 46.90 | 31.45 | 117.11 | 110.56 |
| hsa-miR-523-3p | MIMAT0002840 | GAACGCGCUUCCCUAUAGAGGGU | 101.58 | 100.63 | 115.05 | 107.89 |
| hsa-miR-524-3p | MIMAT0002850 | GAAGGCGCUUCCCUUUGGAGU | 62.30 | 63.63 | 81.67 | 92.34 |
| hsa-miR-524-5p | MIMAT0002849 | CUACAAAGGGAAGCACUUUCUC | 127.02 | 106.37 | 119.06 | 109.29 |
| hsa-miR-525-3p | MIMAT0002839 | GAAGGCGCUUCCCUUUAGAGCG | 93.83 | 117.46 | 116.10 | 141.40 |
| hsa-miR-525-5p | MIMAT0002838 | CUCCAGAGGGAUGCACUUUCU | 111.66 | 129.42 | 107.17 | 95.06 |
| hsa-miR-526b-3p | MIMAT0002836 | GAAAGUGCUUCCUUUUAGAGGC | 176.36 | 184.28 | 86.13 | 110.03 |
| hsa-miR-526b-5p | MIMAT0002835 | CUCUUGAGGGAAGCACUUUCUGU | 63.86 | 92.18 | 109.07 | 100.01 |
| hsa-miR-532-3p | MIMAT0004780 | CCUCCCACACCCAAGGCUUGCA | 112.93 | 92.18 | 77.60 | 75.35 |
| hsa-miR-532-5p | MIMAT0002888 | CAUGCCUUGAGUGUAGGACCGU | 137.87 | 163.67 | 120.42 | 117.54 |
| hsa-miR-539-5p | MIMAT0003163 | GGAGAAAUUAUCCUUGGUGUGU | 140.97 | 167.01 | 109.60 | 117.17 |
| hsa-miR-541-3p | MIMAT0004920 | UGGUGGGCACAGAAUCUGGACU | 64.74 | 57.94 | 120.81 | 124.53 |
| hsa-miR-541-5p | MIMAT0004919 | AAAGGAUUCUGCUGUCGGUCCCACU | 116.50 | 97.56 | 111.06 | 108.53 |
| hsa-miR-542-3p | MIMAT0003389 | UGUGACAGAUUGAUAACUGAAA | 58.49 | 67.39 | 81.94 | 95.48 |
| hsa-miR-542-5p | MIMAT0003340 | UCGGGGAUCAUCAUGUCACGAGA | 94.13 | 105.24 | 103.09 | 106.62 |
| hsa-miR-543 | MIMAT0004954 | AAACAUUCGCGGUGCACUUCUU | 88.46 | 128.52 | 103.39 | 101.72 |
| hsa-miR-544a | MIMAT0003164 | AUUCUGCAUUUUUAGCAAGUUC | 51.10 | 50.43 | 107.64 | 87.16 |
| hsa-miR-544b | MIMAT0015004 | ACCUGAGGUUGUGCAUUUCUAA | 92.43 | 62.61 | 92.60 | 80.02 |
| hsa-miR-545-3p | MIMAT0003165 | UCAGCAAACAUUUAUUGUGUGC | 141.67 | 113.87 | 97.44 | 91.55 |

| | | | | | | |
|---|---|---|---|---|---|---|
| hsa-miR-545-5p | MIMAT0004785 | UCAGUAAAUGUUUAUUAGAUGA | 87.58 | 97.14 | 68.67 | 75.43 |
| hsa-miR-548a-3p | MIMAT0003251 | CAAAACUGGCAAUUACUUUUGC | 118.09 | 97.04 | 134.19 | 110.36 |
| hsa-miR-548a-5p | MIMAT0004803 | AAAAGUAAUUGCGAGUUUUACC | 101.58 | 103.71 | 108.79 | 104.72 |
| hsa-miR-548aa | MIMAT0018447 | AAAAACCACAAUUACUUUUGCACCA | 63.78 | 75.14 | 102.08 | 104.73 |
| hsa-miR-548b-3p | MIMAT0003254 | CAAGAACCUCAGUUGCUUUUGU | 105.89 | 84.06 | 92.80 | 107.03 |
| hsa-miR-548b-5p | MIMAT0004798 | AAAAGUAAUUGUGGUUUUGGCC | 45.67 | 53.01 | 115.05 | 102.82 |
| hsa-miR-548c-3p | MIMAT0003285 | CAAAAAUCUCAAUUACUUUUGC | 97.03 | 74.89 | 99.24 | 83.90 |
| hsa-miR-548c-5p | MIMAT0004806 | AAAAGUAAUUGCGGUUUUUGCC | 68.64 | 69.55 | 113.10 | 123.73 |
| hsa-miR-548d-3p | MIMAT0003323 | CAAAAACCACAGUUUCUUUUGC | 116.12 | 111.10 | 96.20 | 115.00 |
| hsa-miR-548d-5p | MIMAT0004812 | AAAAGUAAUUGUGGUUUUUGCC | 113.16 | 113.11 | 77.21 | 86.91 |
| hsa-miR-548e | MIMAT0005874 | AAAAACUGAGACUACUUUUGCA | 145.78 | 153.49 | 110.18 | 119.82 |
| hsa-miR-548f | MIMAT0005895 | AAAAACUGUAAUUACUUUU | 124.25 | 108.41 | 106.76 | 94.04 |
| hsa-miR-548g-3p | MIMAT0005912 | AAAACUGUAAUUACUUUUGUAC | 105.04 | 105.26 | 74.24 | 91.18 |
| hsa-miR-548h-5p | MIMAT0005928 | AAAAGUAAUCGCGGUUUUUGUC | 94.87 | 111.85 | 97.00 | 96.21 |
| hsa-miR-548i | MIMAT0005935 | AAAAGUAAUUGCGGAUUUUGCC | 124.92 | 105.03 | 110.61 | 107.81 |
| hsa-miR-548j | MIMAT0005875 | AAAAGUAAUUGCGGUCUUUGGU | 86.87 | 84.89 | 83.43 | 76.72 |
| hsa-miR-548k | MIMAT0005882 | AAAAGUACUUGCGGAUUUUGCU | 87.44 | 83.31 | 121.86 | 127.96 |
| hsa-miR-548l | MIMAT0005889 | AAAAGUAUUUGCGGGUUUUGUC | 89.33 | 97.14 | 100.34 | 116.48 |
| hsa-miR-548m | MIMAT0005917 | CAAAGGUAUUUGUGGUUUUUG | 138.79 | 137.26 | 117.45 | 105.50 |
| hsa-miR-548n | MIMAT0005916 | CAAAAGUAAUUGUGGAUUUUGU | 69.21 | 64.79 | 114.78 | 124.00 |
| hsa-miR-548o-3p | MIMAT0005919 | CCAAAACUGCAGUUACUUUUGC | 115.90 | 131.30 | 108.92 | 119.18 |
| hsa-miR-548p | MIMAT0005934 | UAGCAAAAACUGCAGUUACUUU | 53.83 | 46.66 | 111.03 | 92.61 |
| hsa-miR-548q | MIMAT0011163 | GCUGGUGCAAAAGUAAUGGCGG | 107.40 | 119.13 | 116.49 | 121.05 |
| hsa-miR-548s | MIMAT0014987 | AUGGCCAAAACUGCAGUUAUUUU | 88.97 | 92.06 | 92.33 | 103.68 |
| hsa-miR-548t-5p | MIMAT0015009 | CAAAAGUGAUCGUGGUUUUUG | 129.26 | 134.97 | 74.13 | 100.27 |
| hsa-miR-548u | MIMAT0015013 | CAAAGACUGCAAUUACUUUUGCG | 37.60 | 43.57 | 92.51 | 111.74 |
| hsa-miR-548v | MIMAT0015020 | AGCUACAGUUACUUUUGCACCA | 114.94 | 119.66 | 105.87 | 112.02 |
| hsa-miR-548w | MIMAT0015060 | AAAAGUAACUGCGGUUUUUGCCU | 126.02 | 110.08 | 104.02 | 116.45 |
| hsa-miR-548x | MIMAT0015081 | UAAAAACUGCAAUUACUUUCA | 91.77 | 90.27 | 85.77 | 104.51 |
| hsa-miR-548y | MIMAT0018354 | AAAAGUAAUCACUGUUUUUGCC | 126.17 | 116.38 | 136.58 | 121.78 |
| hsa-miR-548z | MIMAT0018446 | CAAAAACCGCAAUUACUUUUGCA | 88.46 | 93.25 | 109.52 | 99.01 |
| hsa-miR-549a | MIMAT0003333 | UGACAACUAUGGAUGAGCUCU | 93.20 | 119.07 | 75.75 | 82.51 |
| hsa-miR-550a-3p | MIMAT0003257 | UGUCUUACUCCCUCAGGCACAU | 85.49 | 106.31 | 115.03 | 104.09 |
| hsa-miR-550a-5p | MIMAT0004800 | AGUGCCUGAGGGAGUAAGAGCCC | 93.78 | 106.37 | 81.72 | 85.83 |
| hsa-miR-550b-3p | MIMAT0018445 | UCUUACUCCCUCAGGCACUG | 116.06 | 77.66 | 98.69 | 71.76 |
| hsa-miR-551a | MIMAT0003214 | GCGACCCACUCUUGGUUUCCA | 134.06 | 119.13 | 126.79 | 127.27 |
| hsa-miR-551b-3p | MIMAT0003233 | GCGACCCAUACUUGGUUUCAG | 120.80 | 124.02 | 125.47 | 120.36 |
| hsa-miR-551b-5p | MIMAT0004794 | GAAAUCAAGCGUGGGUGAGACC | 110.44 | 119.87 | 89.52 | 80.18 |
| hsa-miR-552 | MIMAT0003215 | AACAGGUGACUGGUUAGACAA | 54.23 | 63.07 | 68.29 | 80.92 |
| hsa-miR-553 | MIMAT0003216 | AAAACGGUGAGAUUUUGUUUU | 145.63 | 121.75 | 117.24 | 96.63 |
| hsa-miR-554 | MIMAT0003217 | GCUAGUCCUGACUCAGCCAGU | 71.42 | 99.95 | 88.43 | 97.82 |

| | | | | | | |
|---|---|---|---|---|---|---|
| hsa-miR-555 | MIMAT0003219 | AGGGUAAGCUGAACCUCUGAU | 98.26 | 117.65 | 114.53 | 119.88 |
| hsa-miR-556-3p | MIMAT0004793 | AUAUUACCAUUAGCUCAUCUUU | 166.97 | 214.59 | 139.64 | 151.56 |
| hsa-miR-556-5p | MIMAT0003220 | GAUGAGCUCAUUGUAAUAUGAG | 71.76 | 83.73 | 105.94 | 112.97 |
| hsa-miR-557 | MIMAT0003221 | GUUUGCACGGGUGGGCCUUGUCU | 81.77 | 80.92 | 107.82 | 92.22 |
| hsa-miR-558 | MIMAT0003222 | UGAGCUGCUGUACCAAAAU | 94.20 | 83.14 | 94.27 | 91.42 |
| hsa-miR-559 | MIMAT0003223 | UAAAGUAAAUAUGCACCAAAA | 105.62 | 90.38 | 132.43 | 114.57 |
| hsa-miR-561-3p | MIMAT0003225 | CAAAGUUUAAGAUCCUUGAAGU | 122.98 | 103.15 | 117.76 | 116.60 |
| hsa-miR-562 | MIMAT0003226 | AAAGUAGCUGUACCAUUUGC | 102.21 | 78.40 | 115.49 | 129.94 |
| hsa-miR-563 | MIMAT0003227 | AGGUUGACAUACGUUUCCC | 106.18 | 95.35 | 143.68 | 126.54 |
| hsa-miR-564 | MIMAT0003228 | AGGCACGGUGUCAGCAGGC | 100.47 | 118.21 | 103.73 | 101.09 |
| hsa-miR-566 | MIMAT0003230 | GGGCGCCUGUGAUCCCAAC | 113.13 | 108.91 | 104.51 | 92.30 |
| hsa-miR-567 | MIMAT0003231 | AGUAUGUUCUUCCAGGACAGAAC | 117.48 | 89.11 | 121.89 | 106.66 |
| hsa-miR-568 | MIMAT0003232 | AUGUAUAAAUGUAUACACAC | 94.24 | 94.35 | 99.47 | 114.49 |
| hsa-miR-569 | MIMAT0003234 | AGUUAAUGAAUCCUGGAAAGU | 81.25 | 81.01 | 67.65 | 99.76 |
| hsa-miR-570-3p | MIMAT0003235 | CGAAAACAGCAAUUACCUUUGC | 49.76 | 66.94 | 71.24 | 77.34 |
| hsa-miR-571 | MIMAT0003236 | UGAGUUGGCCAUCUGAGUGAG | 107.67 | 78.40 | 118.41 | 120.47 |
| hsa-miR-572 | MIMAT0003237 | GUCCGCUCGGCGGUGGCCCA | 88.36 | 77.57 | 103.25 | 103.53 |
| hsa-miR-573 | MIMAT0003238 | CUGAAGUGAUGUGUAACUGAUCAG | 136.29 | 125.80 | 125.60 | 133.26 |
| hsa-miR-574-3p | MIMAT0003239 | CACGCUCAUGCACACACCCACA | 113.10 | 125.65 | 94.22 | 100.65 |
| hsa-miR-574-5p | MIMAT0004795 | UGAGUGUGUGUGUGUGAGUGUGU | 103.66 | 110.34 | 119.31 | 99.30 |
| hsa-miR-575 | MIMAT0003240 | GAGCCAGUUGGACAGGAGC | 101.05 | 119.30 | 104.48 | 125.53 |
| hsa-miR-576-3p | MIMAT0004796 | AAGAUGUGGAAAAAUUGGAAUC | 99.52 | 77.80 | 108.49 | 113.36 |
| hsa-miR-576-5p | MIMAT0003241 | AUUCUAAUUUCUCCACGUCUUU | 121.62 | 139.37 | 98.94 | 119.51 |
| hsa-miR-577 | MIMAT0003242 | UAGAUAAAAUAUUGGUACCUG | 129.01 | 118.23 | 118.17 | 106.30 |
| hsa-miR-578 | MIMAT0003243 | CUUCUUGUGCUCUAGGAUUGU | 36.24 | 25.42 | 92.55 | 96.92 |
| hsa-miR-579 | MIMAT0003244 | UUCAUUUGGUAUAAACCGCGAUU | 119.91 | 101.82 | 88.83 | 78.31 |
| hsa-miR-580 | MIMAT0003245 | UUGAGAAUGAUGAAUCAUUAGG | 84.13 | 110.29 | 83.34 | 77.92 |
| hsa-miR-581 | MIMAT0003246 | UCUUGUGUUCUCUAGAUCAGU | 119.23 | 95.48 | 112.77 | 117.65 |
| hsa-miR-582-3p | MIMAT0004797 | UAACUGGUUGAACAACUGAACC | 99.16 | 114.48 | 91.38 | 110.64 |
| hsa-miR-582-5p | MIMAT0003247 | UUACAGUUGUUCAACCAGUUACU | 189.17 | 216.00 | 73.30 | 89.05 |
| hsa-miR-583 | MIMAT0003248 | CAAAGAGGAAGGUCCCAUUAC | 114.94 | 109.23 | 91.18 | 92.58 |
| hsa-miR-584-5p | MIMAT0003249 | UUAUGGUUUGCCUGGGACUGAG | 69.99 | 68.86 | 79.47 | 76.33 |
| hsa-miR-585 | MIMAT0003250 | UGGGCGUAUCUGUAUGCUA | 98.12 | 86.08 | 97.84 | 127.84 |
| hsa-miR-586 | MIMAT0003252 | UAUGCAUUGUAUUUUUAGGUCC | 134.76 | 154.64 | 101.47 | 113.44 |
| hsa-miR-587 | MIMAT0003253 | UUUCCAUAGGUGAUGAGUCAC | 106.78 | 108.51 | 89.13 | 67.39 |
| hsa-miR-588 | MIMAT0003255 | UUGGCCACAAUGGGUUAGAAC | 110.85 | 123.14 | 93.08 | 84.88 |
| hsa-miR-589-3p | MIMAT0003256 | UCAGAACAAAUGCCGGUUCCCAGA | 112.30 | 105.47 | 103.22 | 109.01 |
| hsa-miR-589-5p | MIMAT0004799 | UGAGAACCACGUCUGCUCUGAG | 96.49 | 83.97 | 91.42 | 97.85 |
| hsa-miR-590-3p | MIMAT0004801 | UAAUUUUAUGUAUAAGCUAGU | 98.12 | 98.37 | 103.94 | 107.26 |
| hsa-miR-590-5p | MIMAT0003258 | GAGCUUAUUCAUAAAAGUGCAG | 132.32 | 132.09 | 110.04 | 96.57 |
| hsa-miR-591 | MIMAT0003259 | AGACCAUGGGUUCUCAUUGU | 81.30 | 78.96 | 86.34 | 92.05 |

| | | | | | | |
|---|---|---|---|---|---|---|
| hsa-miR-592 | MIMAT0003260 | UUGUGUCAAUAUGCGAUGAUGU | 119.44 | 112.16 | 111.90 | 120.11 |
| hsa-miR-593-3p | MIMAT0004802 | UGUCUCUGCUGGGGUUUCU | 87.86 | 115.09 | 109.29 | 96.21 |
| hsa-miR-593-5p | MIMAT0003261 | AGGCACCAGCCAGGCAUUGCUCAGC | 49.59 | 52.20 | 123.81 | 118.36 |
| hsa-miR-595 | MIMAT0003263 | GAAGUGUGCCGUGGUGUGUCU | 131.07 | 136.90 | 100.66 | 100.55 |
| hsa-miR-596 | MIMAT0003264 | AAGCCUGCCCGGCUCCUCGGG | 130.00 | 95.35 | 58.61 | 61.93 |
| hsa-miR-597 | MIMAT0003265 | UGUGUCACUCGAUGACCACUGU | 178.63 | 153.83 | 66.09 | 85.66 |
| hsa-miR-598 | MIMAT0003266 | UACGUCAUCGUUGUCAUCGUCA | 117.10 | 111.45 | 99.30 | 80.76 |
| hsa-miR-599 | MIMAT0003267 | GUUGUGUCAGUUUAUCAAAC | 112.18 | 124.25 | 89.86 | 88.60 |
| hsa-miR-600 | MIMAT0003268 | ACUUACAGACAAGAGCCUUGCUC | 134.72 | 149.58 | 103.97 | 120.53 |
| hsa-miR-601 | MIMAT0003269 | UGGUCUAGGAUUGUUGGAGGAG | 92.80 | 94.52 | 86.00 | 106.79 |
| hsa-miR-602 | MIMAT0003270 | GACACGGGCGACAGCUGCGGCCC | 120.85 | 119.45 | 134.62 | 145.56 |
| hsa-miR-603 | MIMAT0003271 | CACACACUGCAAUUACUUUUGC | 120.21 | 87.86 | 99.24 | 99.90 |
| hsa-miR-604 | MIMAT0003272 | AGGCUGCGGAAUUCAGGAC | 133.97 | 119.08 | 137.67 | 124.68 |
| hsa-miR-605 | MIMAT0003273 | UAAAUCCCAUGGUGCCUUCUCCU | 137.68 | 135.98 | 80.57 | 86.13 |
| hsa-miR-606 | MIMAT0003274 | AAACUACUGAAAAUCAAAGAU | 111.50 | 105.12 | 91.75 | 95.78 |
| hsa-miR-607 | MIMAT0003275 | GUUCAAAUCCAGAUCUAUAAC | 90.25 | 97.94 | 94.76 | 111.40 |
| hsa-miR-608 | MIMAT0003276 | AGGGGUGGUGUUGGGACAGCUCCGU | 41.01 | 42.25 | 125.30 | 122.49 |
| hsa-miR-609 | MIMAT0003277 | AGGGUGUUUCUCUCAUCUCU | 81.87 | 88.57 | 100.80 | 124.03 |
| hsa-miR-610 | MIMAT0003278 | UGAGCUAAAUGUGUGCUGGGA | 167.45 | 190.86 | 124.00 | 131.65 |
| hsa-miR-611 | MIMAT0003279 | GCGAGGACCCCUCGGGGUCUGAC | 116.54 | 120.62 | 115.02 | 95.75 |
| hsa-miR-612 | MIMAT0003280 | GCUGGGCAGGGCUUCUGAGCUCCUU | 70.35 | 70.08 | 88.83 | 94.42 |
| hsa-miR-613 | MIMAT0003281 | AGGAAUGUUCCUUCUUUGCC | 134.37 | 109.55 | 91.69 | 78.40 |
| hsa-miR-614 | MIMAT0003282 | GAACGCCUGUUCUUGCCAGGUGG | 89.95 | 112.65 | 119.47 | 113.59 |
| hsa-miR-615-3p | MIMAT0003283 | UCCGAGCCUGGGUCUCCCUCUU | 54.98 | 47.66 | 115.99 | 129.44 |
| hsa-miR-615-5p | MIMAT0004804 | GGGGGUCCCCGGUGCUCGGAUC | 120.93 | 121.01 | 99.79 | 99.48 |
| hsa-miR-616-3p | MIMAT0004805 | AGUCAUUGGAGGGUUUGAGCAG | 132.32 | 115.43 | 123.90 | 115.96 |
| hsa-miR-616-5p | MIMAT0003284 | ACUCAAAACCCUUCAGUGACUU | 118.77 | 112.40 | 117.82 | 94.15 |
| hsa-miR-617 | MIMAT0003286 | AGACUUCCCAUUUGAAGGUGGC | 45.95 | 78.08 | 83.47 | 80.71 |
| hsa-miR-618 | MIMAT0003287 | AAACUCUACUUGUCCUUCUGAGU | 97.03 | 108.61 | 85.83 | 105.64 |
| hsa-miR-619 | MIMAT0003288 | GACCUGGACAUGUUUGUGCCCAGU | 118.19 | 125.35 | 83.43 | 123.09 |
| hsa-miR-620 | MIMAT0003289 | AUGGAGAUAGAUAUAGAAAU | 121.70 | 89.85 | 96.16 | 92.90 |
| hsa-miR-621 | MIMAT0003290 | GGCUAGCAACAGCGCUUACCU | 79.47 | 112.01 | 113.45 | 127.01 |
| hsa-miR-622 | MIMAT0003291 | ACAGUCUGCUGAGGUUGGAGC | 94.45 | 90.18 | 90.50 | 101.59 |
| hsa-miR-623 | MIMAT0003292 | AUCCCUUGCAGGGGCUGUUGGGU | 73.99 | 108.45 | 101.60 | 81.24 |
| hsa-miR-624-3p | MIMAT0004807 | CACAAGGUAUUGGUAUUACCU | 84.73 | 97.65 | 104.54 | 118.42 |
| hsa-miR-624-5p | MIMAT0003293 | UAGUACCAGUACCUUGUGUUCA | 92.82 | 83.62 | 134.99 | 126.51 |
| hsa-miR-625-3p | MIMAT0004808 | GACUAUAGAACUUUCCCCCUCA | 110.23 | 109.75 | 102.98 | 112.64 |
| hsa-miR-625-5p | MIMAT0003294 | AGGGGGAAAGUUCUAUAGUCC | 113.73 | 87.27 | 90.08 | 77.57 |
| hsa-miR-626 | MIMAT0003295 | AGCUGUCUGAAAAUGUCUU | 87.14 | 87.23 | 95.28 | 90.54 |
| hsa-miR-627 | MIMAT0003296 | GUGAGUCUCUAAGAAAAGAGGA | 117.60 | 124.17 | 99.75 | 101.53 |
| hsa-miR-628-3p | MIMAT0003297 | UCUAGUAAGAGUGGCAGUCGA | 94.97 | 91.77 | 100.55 | 96.31 |

| | | | | | | |
|---|---|---|---|---|---|---|
| hsa-miR-628-5p | MIMAT0004809 | AUGCUGACAUAUUUACUAGAGG | 36.17 | 37.29 | 87.81 | 76.46 |
| hsa-miR-629-3p | MIMAT0003298 | GUUCUCCCAACGUAAGCCCAGC | 85.17 | 98.23 | 75.40 | 72.00 |
| hsa-miR-629-5p | MIMAT0004810 | UGGGUUUACGUUGGGAGAACU | 69.99 | 70.60 | 87.15 | 89.40 |
| hsa-miR-630 | MIMAT0003299 | AGUAUUCUGUACCAGGGAAGGU | 168.45 | 149.05 | 107.61 | 75.35 |
| hsa-miR-631 | MIMAT0003300 | AGACCUGGCCCAGACCUCAGC | 113.81 | 123.69 | 87.93 | 94.95 |
| hsa-miR-632 | MIMAT0003302 | GUGUCUGCUUCCUGUGGGA | 107.81 | 106.51 | 76.14 | 107.27 |
| hsa-miR-633 | MIMAT0003303 | CUAAUAGUAUCUACCACAAUAAA | 86.25 | 85.91 | 92.53 | 82.26 |
| hsa-miR-634 | MIMAT0003304 | AACCAGCACCCCAACUUUGGAC | 40.19 | 67.38 | 149.97 | 150.99 |
| hsa-miR-635 | MIMAT0003305 | ACUUGGGCACUGAAACAAUGUCC | 77.40 | 91.11 | 83.02 | 94.48 |
| hsa-miR-636 | MIMAT0003306 | UGUGCUUGCUCGUCCCGCCCGCA | 123.78 | 106.25 | 103.68 | 92.53 |
| hsa-miR-637 | MIMAT0003307 | ACUGGGGGCUUUCGGGCUCUGCGU | 85.38 | 84.55 | 95.55 | 114.24 |
| hsa-miR-638 | MIMAT0003308 | AGGGAUCGCGGGCGGGUGGCGGCCU | 81.94 | 90.45 | 108.17 | 99.90 |
| hsa-miR-639 | MIMAT0003309 | AUCGCUGCGGUUGCGAGCGCUGU | 83.07 | 69.73 | 100.30 | 85.15 |
| hsa-miR-640 | MIMAT0003310 | AUGAUCCAGGAACCUGCCUCU | 91.75 | 77.92 | 98.29 | 80.02 |
| hsa-miR-641 | MIMAT0003311 | AAAGACAUAGGAUAGAGUCACCUC | 87.68 | 114.19 | 87.51 | 99.53 |
| hsa-miR-642a-5p | MIMAT0003312 | GUCCCUCUCCAAAUGUGUCUUG | 101.22 | 92.81 | 121.46 | 126.07 |
| hsa-miR-642b-3p | MIMAT0018444 | AGACACAUUUGGAGAGGGACCC | 69.80 | 85.48 | 93.46 | 86.51 |
| hsa-miR-643 | MIMAT0003313 | ACUUGUAUGCUAGCUCAGGUAG | 137.30 | 103.77 | 89.86 | 103.74 |
| hsa-miR-644a | MIMAT0003314 | AGUGUGGCUUUCUUAGAGC | 63.07 | 61.89 | 79.84 | 77.96 |
| hsa-miR-645 | MIMAT0003315 | UCUAGGCUGGUACUGCUGA | 121.16 | 103.71 | 113.91 | 110.43 |
| hsa-miR-646 | MIMAT0003316 | AAGCAGCUGCCUCUGAGGC | 65.35 | 84.36 | 69.58 | 90.23 |
| hsa-miR-647 | MIMAT0003317 | GUGGCUGCACUCACUUCCUUC | 97.61 | 97.30 | 82.19 | 98.82 |
| hsa-miR-648 | MIMAT0003318 | AAGUGUGCAGGGCACUGGU | 94.90 | 112.36 | 105.58 | 116.13 |
| hsa-miR-649 | MIMAT0003319 | AAACCUGUGUUGUUCAAGAGUC | 71.95 | 116.54 | 85.70 | 107.17 |
| hsa-miR-650 | MIMAT0003320 | AGGAGGCAGCGCUCUCAGGAC | 94.28 | 81.19 | 119.84 | 123.73 |
| hsa-miR-651 | MIMAT0003321 | UUUAGGAUAAGCUUGACUUUUG | 72.23 | 77.48 | 85.09 | 81.03 |
| hsa-miR-652-3p | MIMAT0003322 | AAUGGCGCCACUAGGGUUGUG | 73.10 | 105.15 | 117.80 | 112.21 |
| hsa-miR-653 | MIMAT0003328 | GUGUUGAAACAAUCUCUACUG | 35.41 | 45.32 | 84.29 | 95.20 |
| hsa-miR-654-3p | MIMAT0004814 | UAUGUCUGCUGACCAUCACCUU | 165.29 | 171.83 | 87.06 | 82.51 |
| hsa-miR-654-5p | MIMAT0003330 | UGGUGGGCCGCAGAACAUGUGC | 96.31 | 70.55 | 102.90 | 116.48 |
| hsa-miR-655 | MIMAT0003331 | AUAAUACAUGGUUAACCUCUUU | 37.88 | 35.70 | 76.69 | 62.99 |
| hsa-miR-656 | MIMAT0003332 | AAUAUUAUACAGUCAACCUCU | 109.51 | 101.46 | 88.58 | 81.86 |
| hsa-miR-657 | MIMAT0003335 | GGCAGGUUCUCACCCUCUCUAGG | 103.21 | 113.15 | 103.41 | 131.52 |
| hsa-miR-658 | MIMAT0003336 | GGCGGAGGGAAGUAGGUCCGUUGGU | 98.26 | 102.80 | 115.02 | 104.70 |
| hsa-miR-659-3p | MIMAT0003337 | CUUGGUUCAGGGAGGGUCCCCA | 91.72 | 89.94 | 93.27 | 91.50 |
| hsa-miR-660-5p | MIMAT0003338 | UACCCAUUGCAUAUCGGAGUUG | 99.82 | 110.59 | 88.72 | 109.69 |
| hsa-miR-661 | MIMAT0003324 | UGCCUGGGUCUCUGGCCUGCGCGU | 85.17 | 81.37 | 77.64 | 77.33 |
| hsa-miR-662 | MIMAT0003325 | UCCCACGUUGUGGCCCAGCAG | 142.02 | 148.84 | 104.51 | 120.61 |
| hsa-miR-663a | MIMAT0003326 | AGGCGGGGCGCCGCGGGACCGC | 94.28 | 108.69 | 93.54 | 115.28 |
| hsa-miR-663b | MIMAT0005867 | GGUGGCCCGGCCGUGCCUGAGG | 109.04 | 117.53 | 120.75 | 105.99 |
| hsa-miR-664a-3p | MIMAT0005949 | UAUUCAUUUAUCCCCAGCCUACA | 126.57 | 128.25 | 137.13 | 116.48 |

| | | | | | | |
|---|---|---|---|---|---|---|
| hsa-miR-664a-5p | MIMAT0005948 | ACUGGCUAGGGAAAAUGAUUGGAU | 94.37 | 92.67 | 90.59 | 92.90 |
| hsa-miR-665 | MIMAT0004952 | ACCAGGAGGCUGAGGCCCCU | 78.74 | 78.19 | 97.22 | 108.24 |
| hsa-miR-668 | MIMAT0003881 | UGUCACUCGGCUCGGCCCACUAC | 165.38 | 150.52 | 102.71 | 103.56 |
| hsa-miR-670 | MIMAT0010357 | GUCCCUGAGUGUAUGUGGUG | 68.55 | 62.72 | 76.27 | 93.80 |
| hsa-miR-671-3p | MIMAT0004819 | UCCGGUUCUCAGGGCUCCACC | 105.48 | 90.56 | 93.37 | 70.21 |
| hsa-miR-671-5p | MIMAT0003880 | AGGAAGCCCUGGAGGGGCUGGAG | 109.95 | 112.56 | 103.46 | 84.10 |
| hsa-miR-675-3p | MIMAT0006790 | CUGUAUGCCCUCACCGCUCA | 88.75 | 121.94 | 77.66 | 69.22 |
| hsa-miR-675-5p | MIMAT0004284 | UGGUGCGGAGAGGGCCCACAGUG | 90.28 | 89.87 | 94.29 | 98.64 |
| hsa-miR-676-3p | MIMAT0018204 | CUGUCCUAAGGUUGUUGAGUU | 101.25 | 92.18 | 118.13 | 121.46 |
| hsa-miR-676-5p | MIMAT0018203 | UCUUCAACCUCAGGACUUGCA | 112.15 | 134.47 | 74.40 | 74.27 |
| hsa-miR-708-3p | MIMAT0004927 | CAACUAGACUGUGAGCUUCUAG | 93.41 | 103.93 | 114.80 | 98.90 |
| hsa-miR-708-5p | MIMAT0004926 | AAGGAGCUUACAAUCUAGCUGGG | 61.24 | 74.31 | 49.38 | 50.05 |
| hsa-miR-711 | MIMAT0012734 | GGGACCCAGGGAGAGACGUAAG | 83.84 | 97.62 | 122.59 | 105.74 |
| hsa-miR-7-1-3p | MIMAT0004553 | CAACAAAUCACAGUCUGCCAUA | 94.57 | 117.88 | 99.91 | 111.71 |
| hsa-miR-718 | MIMAT0012735 | CUUCCGCCCCGCCGGGCGUCG | 63.19 | 74.97 | 86.20 | 102.11 |
| hsa-miR-720 | MIMAT0005954 | UCUCGCUGGGGCCUCCA | 141.27 | 151.30 | 92.18 | 113.06 |
| hsa-miR-7-2-3p | MIMAT0004554 | CAACAAAUCCCAGUCUACCUAA | 54.23 | 63.78 | 86.13 | 58.65 |
| hsa-miR-744-3p | MIMAT0004946 | CUGUUGCCACUAACCUCAACCU | 57.82 | 61.09 | 84.11 | 75.93 |
| hsa-miR-744-5p | MIMAT0004945 | UGCGGGGCUAGGGCUAACAGCA | 107.81 | 122.87 | 105.94 | 90.05 |
| hsa-miR-758-3p | MIMAT0003879 | UUUGUGACCUGGUCCACUAACC | 105.83 | 99.86 | 80.04 | 80.50 |
| hsa-miR-759 | MIMAT0010497 | GCAGAGUGCAAACAAUUUUGAC | 82.42 | 86.01 | 81.71 | 81.63 |
| hsa-miR-7-5p | MIMAT0000252 | UGGAAGACUAGUGAUUUUGUUGU | 110.83 | 135.51 | 134.86 | 155.41 |
| hsa-miR-760 | MIMAT0004957 | CGGCUCUGGGUCUGUGGGGA | 93.26 | 88.51 | 94.77 | 90.46 |
| hsa-miR-761 | MIMAT0010364 | GCAGCAGGGUGAAACUGACACA | 94.41 | 114.64 | 106.16 | 95.78 |
| hsa-miR-762 | MIMAT0010313 | GGGGCUGGGGCCGGGGCCGAGC | 83.84 | 88.91 | 109.07 | 101.17 |
| hsa-miR-764 | MIMAT0010367 | GCAGGUGCUCACUUGUCCUCCU | 39.51 | 32.29 | 89.52 | 98.47 |
| hsa-miR-765 | MIMAT0003945 | UGGAGGAGAAGGAAGGUGAUG | 96.26 | 110.60 | 70.10 | 86.99 |
| hsa-miR-766-3p | MIMAT0003888 | ACUCCAGCCCCACAGCCUCAGC | 80.97 | 81.03 | 57.97 | 61.65 |
| hsa-miR-767-3p | MIMAT0003883 | UCUGCUCAUACCCCAUGGUUUCU | 101.25 | 103.97 | 97.73 | 107.04 |
| hsa-miR-767-5p | MIMAT0003882 | UGCACCAUGGUUGUCUGAGCAUG | 68.52 | 44.52 | 101.30 | 94.95 |
| hsa-miR-768-3p | MIMAT0003947 | UCACAAUGCUGACACUCAAACUGCUGAC | 71.40 | 69.62 | 106.92 | 122.69 |
| hsa-miR-768-5p | MIMAT0003946 | GUUGGAGGAUGAAAGUACGGAGUGAU | 98.34 | 87.37 | 102.16 | 91.09 |
| hsa-miR-769-3p | MIMAT0003887 | CUGGGAUCUCCGGGGUCUUGGUU | 105.62 | 91.73 | 97.34 | 95.00 |
| hsa-miR-769-5p | MIMAT0003886 | UGAGACCUCUGGGUUCUGAGCU | 75.82 | 55.14 | 106.71 | 96.57 |
| hsa-miR-770-5p | MIMAT0003948 | UCCAGUACCACGUGUCAGGGCCA | 87.59 | 97.61 | 99.33 | 100.81 |
| hsa-miR-801 | MIMAT0004209 | GAUUGCUCUGCGUGCGGAAUCGAC | 88.20 | 95.78 | 94.28 | 81.16 |
| hsa-miR-802 | MIMAT0004185 | CAGUAACAAAGAUUCAUCCUUGU | 94.61 | 76.71 | 89.70 | 90.71 |
| hsa-miR-873-5p | MIMAT0004953 | GCAGGAACUUGUGAGUCUCCU | 80.07 | 95.50 | 70.08 | 81.67 |
| hsa-miR-874 | MIMAT0004911 | CUGCCCUGGCCCGAGGGACCGA | 137.68 | 91.52 | 79.84 | 102.48 |
| hsa-miR-875-3p | MIMAT0004923 | CCUGGAAACACUGAGGUUGUG | 122.38 | 125.22 | 115.55 | 95.51 |
| hsa-miR-875-5p | MIMAT0004922 | UAUACCUCAGUUUUAUCAGGUG | 37.73 | 43.79 | 101.95 | 83.14 |

| | | | | | | |
|---|---|---|---|---|---|---|
| hsa-miR-876-3p | MIMAT0004925 | UGGUGGUUUACAAAGUAAUUCA | 73.85 | 78.13 | 87.32 | 116.72 |
| hsa-miR-876-5p | MIMAT0004924 | UGGAUUUCUUUGUGAAUCACCA | 97.12 | 113.45 | 60.51 | 65.37 |
| hsa-miR-877-3p | MIMAT0004950 | UCCUCUUCUCCCUCCUCCCAG | 117.51 | 100.18 | 102.96 | 99.90 |
| hsa-miR-877-5p | MIMAT0004949 | GUAGAGGAGAUGGCGCAGGG | 73.02 | 73.64 | 108.42 | 67.22 |
| hsa-miR-885-3p | MIMAT0004948 | AGGCAGCGGGGUGUAGUGGAUA | 57.89 | 61.98 | 86.57 | 84.46 |
| hsa-miR-885-5p | MIMAT0004947 | UCCAUUACACUACCCUGCCUCU | 106.79 | 86.32 | 89.21 | 109.80 |
| hsa-miR-886-3p | MIMAT0004906 | CGCGGGUGCUUACUGACCCUU | 100.35 | 81.40 | 109.54 | 122.87 |
| hsa-miR-887 | MIMAT0004951 | GUGAACGGGCGCCAUCCCGAGG | 62.30 | 76.71 | 106.15 | 98.88 |
| hsa-miR-888-3p | MIMAT0004917 | GACUGACACCUCUUUGGGUGAA | 108.17 | 115.69 | 92.25 | 78.37 |
| hsa-miR-888-5p | MIMAT0004916 | UACUCAAAAAGCUGUCAGUCA | 102.82 | 112.93 | 92.87 | 101.02 |
| hsa-miR-889 | MIMAT0004921 | UUAAUAUCGGACAACCAUUGU | 72.63 | 67.97 | 93.21 | 87.02 |
| hsa-miR-890 | MIMAT0004912 | UACUUGGAAAGGCAUCAGUUG | 65.22 | 88.07 | 81.41 | 98.38 |
| hsa-miR-891a | MIMAT0004902 | UGCAACGAACCUGAGCCACUGA | 92.67 | 101.13 | 89.47 | 94.62 |
| hsa-miR-891b | MIMAT0004913 | UGCAACUUACCUGAGUCAUUGA | 50.33 | 53.01 | 62.65 | 55.22 |
| hsa-miR-892a | MIMAT0004907 | CACUGUGUCCUUUCUGCGUAG | 104.75 | 95.86 | 82.90 | 82.43 |
| hsa-miR-892b | MIMAT0004918 | CACUGGCUCCUUUCUGGGUAGA | 83.79 | 71.63 | 110.61 | 114.82 |
| hsa-miR-920 | MIMAT0004970 | GGGGAGCUGUGGAAGCAGUA | 93.07 | 123.77 | 88.24 | 77.31 |
| hsa-miR-921 | MIMAT0004971 | CUAGUGAGGGACAGAACCAGGAUUC | 91.53 | 101.11 | 101.03 | 115.22 |
| hsa-miR-922 | MIMAT0004972 | GCAGCAGAGAAUAGGACUACGUC | 93.52 | 97.60 | 129.61 | 105.75 |
| hsa-miR-923 | MIMAT0004973 | GUCAGCGGAGGAAAAGAAACU | 118.77 | 89.12 | 128.60 | 112.90 |
| hsa-miR-924 | MIMAT0004974 | AGAGUCUUGUGAUGUCUUGC | 93.20 | 96.02 | 92.84 | 106.62 |
| hsa-miR-92a-1-5p | MIMAT0004507 | AGGUUGGGAUCGGUUGCAAUGCU | 108.45 | 114.19 | 97.01 | 109.34 |
| hsa-miR-92a-2-5p | MIMAT0004508 | GGGUGGGGAUUUGUUGCAUUAC | 67.38 | 85.27 | 66.47 | 75.69 |
| hsa-miR-92a-3p | MIMAT0000092 | UAUUGCACUUGUCCCGGCCUGU | 48.52 | 38.78 | 93.53 | 93.68 |
| hsa-miR-92b-3p | MIMAT0003218 | UAUUGCACUCGUCCCGGCCUCC | 57.25 | 59.50 | 103.66 | 84.91 |
| hsa-miR-92b-5p | MIMAT0004792 | AGGGACGGGACGCGGUGCAGUG | 111.46 | 105.36 | 99.51 | 116.88 |
| hsa-miR-933 | MIMAT0004976 | UGUGCGCAGGGAGACCUCUCCC | 65.24 | 77.59 | 101.95 | 123.13 |
| hsa-miR-93-3p | MIMAT0004509 | ACUGCUGAGCUAGCACUUCCCG | 124.65 | 82.27 | 142.11 | 122.47 |
| hsa-miR-934 | MIMAT0004977 | UGUCUACUACUGGAGACACUGG | 132.53 | 142.14 | 108.65 | 109.76 |
| hsa-miR-935 | MIMAT0004978 | CCAGUUACCGCUUCCGCUACCGC | 187.10 | 169.72 | 144.41 | 140.35 |
| hsa-miR-93-5p | MIMAT0000093 | CAAAGUGCUGUUCGUGCAGGUAG | 86.70 | 83.41 | 116.88 | 130.70 |
| hsa-miR-936 | MIMAT0004979 | ACAGUAGAGGGAGGAAUCGCAG | 116.68 | 118.78 | 103.33 | 111.71 |
| hsa-miR-937-3p | MIMAT0004980 | AUCCGCGCUCUGACUCUCUGCC | 119.13 | 105.37 | 112.64 | 111.38 |
| hsa-miR-938 | MIMAT0004981 | UGCCCUUAAAGGUGAACCCAGU | 123.02 | 111.39 | 121.89 | 126.93 |
| hsa-miR-939-5p | MIMAT0004982 | UGGGGAGCUGAGGCUCUGGGGGUG | 110.59 | 108.45 | 112.27 | 130.58 |
| hsa-miR-9-3p | MIMAT0000442 | AUAAAGCUAGAUAACCGAAAGU | 107.51 | 105.23 | 100.13 | 105.42 |
| hsa-miR-940 | MIMAT0004983 | AAGGCAGGGCCCCCGCUCCCC | 77.40 | 57.21 | 96.75 | 119.53 |
| hsa-miR-941 | MIMAT0004984 | CACCCGGCUGUGUGCACAUGUGC | 136.56 | 146.05 | 119.31 | 108.48 |
| hsa-miR-942 | MIMAT0004985 | UCUUCUCUGUUUUGGCCAUGUG | 111.97 | 110.97 | 90.50 | 78.62 |
| hsa-miR-943 | MIMAT0004986 | CUGACUGUUGCCGUCCUCCAG | 78.15 | 118.04 | 92.26 | 112.17 |
| hsa-miR-944 | MIMAT0004987 | AAAUUAUUGUACAUCGGAUGAG | 128.29 | 122.80 | 110.05 | 94.11 |

| | | | | | | |
|---|---|---|---|---|---|---|
| hsa-miR-95 | MIMAT0000094 | UUCAACGGGUAUUUAUUGAGCA | 136.14 | 127.26 | 109.80 | 86.46 |
| hsa-miR-9-5p | MIMAT0000441 | UCUUUGGUUAUCUAGCUGUAUGA | 55.86 | 71.68 | 75.75 | 83.11 |
| hsa-miR-96-3p | MIMAT0004510 | AAUCAUGUGCAGUGCCAAUAUG | 100.66 | 77.09 | 116.55 | 104.68 |
| hsa-miR-96-5p | MIMAT0000095 | UUUGGCACUAGCACAUUUUUGCU | 114.35 | 112.02 | 120.88 | 109.32 |
| hsa-miR-98-5p | MIMAT0000096 | UGAGGUAGUAAGUUGUAUUGUU | 22.99 | 38.54 | 68.25 | 57.76 |
| hsa-miR-99a-3p | MIMAT0004511 | CAAGCUCGCUUCUAUGGGUCUG | 122.63 | 101.32 | 88.53 | 92.63 |
| hsa-miR-99a-5p | MIMAT0000097 | AACCCGUAGAUCCGAUCUUGUG | 102.71 | 82.64 | 104.46 | 91.93 |
| hsa-miR-99b-3p | MIMAT0004678 | CAAGCUCGUGUCUGUGGGUCCG | 140.73 | 160.55 | 117.33 | 98.37 |
| hsa-miR-99b-5p | MIMAT0000689 | CACCCGUAGAACCGACCUUGCG | 116.91 | 134.23 | 93.36 | 101.82 |
