## Supplementray Tables 2-4 for "MicroRNA-541-3p alters lipoproteins to reduce atherosclerosis by degrading Znf101 and Casz1 transcription factors"

**Table S2. Physical parameters in different mice transduced with different shRNAs measured using dual-energy x-ray absorptiometry (DEXA).** Mice were transduced with different shRNAs as described in Fig 4. DEXA was performed 3 months after the injection of viruses. All KD mice showed significantly less fat weight. Data are average +SD, Student *t*-test compared to controls.

| DEXA parameters | shCtrl (n=4) | shCasz1 (n=4) | shZfp961 (n=4) | sh(C+Z) (n=5) |
|---|---|---|---|---|
| Total Weight (gms) | 23.11 ± 0.55 | 24.60 ± 1.32<br>P = 1.72 | 23.59 ± 0.18<br>P = 0.86 | 21.31 ± 1.50,<br>P = 0.06 |
| Soft Weight (gms) | 21.91 ± 0.49 | 23.61 ± 1.42<br>P = 0.13 | 22.63 ± 0.13<br>P = 0.70 | 20.34 ± 1.60<br>P = 0.14 |
| Lean Weight (gms) | 12.82 ± 2.92 | 16.20 ± 2.92<br>P = 0.19 | 16.38 ± 0.95<br>P = 0.17 | 13.99 ± 2.75<br>P = 0.88 |
| Fat Weight (gms) | 10.59 ± 0.75 | 7.41 ± 1.69<br>**P = 0.001** | 6.25 ± 0.86<br>**P = 0.0002** | 6.35 ±1.75<br>**p<0.0001** |
| Fat (% of body weight) | 46.46 ± 4.05 | 31.72 ± 8.83<br>**P = 0.003** | 27.64 ± 3.92<br>**P = 0.0005** | 31.56 ± 0.94<br>**P = 0.0003** |
| Bone Mineral Content (gms) | 1.25 ± 0.17 | 0.98 ± 0.11<br>P = 0.08 | 0.96 ± 0.11<br>P = 0.052 | 0.97 ± 0.18<br>**P = 0.05** |
| Bone Mineral Density (mg/cm$^2$) | 88.66 ± 3.82 | 87.76 ± 3.72<br>P = 0.99 | 90.52 ± 7.65<br>P = 0.94 | 88.42 ± 6.85<br>P = 0.99 |

**Table S3. Total weight gain was less in different mice transduced with shRNAs to KD Casz1 and Zfp961.** Mice were transduced with different shRNAs as described in Fig 5. DEXA was performed 3 months after the injection of viruses. All KD mice showed significantly less total; soft, lean and fat weights. Data are average +SD, Student *t*-test compared to controls.

| DEXA Parameters | mPcsk9+shCtrl (n=4) | mPcsk9+shCasz1 (n=4) | mPcsk9+shZfp961 (n=4) | mPcsk9+sh(C+Z) (n=5) |
|---|---|---|---|---|
| Total Weight (gms) | 40.97 ± 0.82 | 34.18 ± 2.58 **P = 0.024** | 36.17 ± 1.47 **P = 0.001** | 30.58 ± 3.16 **P = 0.0004** |
| Soft Weight (gms) | 40.00 ± 0.81 | 33.28 ± 2.59 **P = 0.002** | 35.21 ± 1.38 **P = 0.001** | 29.61 ± 3.17 **P = 0.002** |
| Lean Weight (gms) | 27.05 ± 0.49 | 22.83 ± 1.35 **P = 0.001** | 23.47 ± 2.61 **P = 0.041** | 20.07 ± 2.54 **P = 0.0004** |
| Fat Weight (gms) | 13.17 ± 0.58 | 10.45 ± 1.37 **P = 0.013** | 10.24 ± 1.09 **P = 0.005** | 9.54 ± 1.01 **P = 0.0006** |
| Fat (% of body weight) | 32.93 ± 1.12 | 31.31 ± 1.92 P = 0.19 | 33.25 ± 8.01 P = 0.94 | 32.27 ± 2.46 P = 0.64 |
| Bone Mineral Content (gms) | 0.98 ± 0.01 | 0.90 ± 0.02 **P = 0.003** | 0.96 ± 0.10 P = 0.82 | 0.96 ± 0.08 P = 0.76 |
| Bone Mineral Density ($mg/cm^2$) | 86.32 ± 1.50 | 85.38 ± 2.26 P = 0.51 | 86.12 ± 0.77 P = 0.81 | 89.55 ± 2.30 P = 0.04 |

Data are average +SD. Student t-test.

Table S4. Genetic associations of *MIR541*, *ZNZF101* and *CASZ1* genes with plasma lipid and lipoproteins.

| *Gene_name* | *chr_pos_ref_alt* | *SNP_effect_allele* | *allele_frequency* | *Trait* | *Beta* | *Std_error* | *p_value* |
|---|---|---|---|---|---|---|---|
| **miR-541-3p** | 14_101529005_A_G | rs7161194-A | 6.61E-01 | Cholesterol | 3.86E-03 | 2.62E-03 | 1.41E-01 |
| | 14_101529005_A_G | rs7161194-A | 6.61E-01 | LDL direct | 1.37E-03 | 2.60E-03 | 5.99E-01 |
| | 14_101529005_A_G | rs7161194-A | 6.61E-01 | Apolipoprotein B | -1.79E-03 | 2.71E-03 | 5.07E-01 |
| | 14_101529005_A_G | rs7161194-A | 6.61E-01 | HDL cholesterol | 1.53E-02 | 2.75E-03 | **2.63E-08** |
| | 14_101529005_A_G | rs7161194-A | 6.61E-01 | Apolipoprotein A | 1.29E-02 | 2.72E-03 | 2.07E-06 |
| | 14_101529005_A_G | rs7161194-A | 6.61E-01 | Triglycerides | -7.21E-03 | 2.70E-03 | 7.50E-03 |
| **CASZ1** | 1_10798489_C_G | rs34071855-G | 3.42E-01 | Cholesterol | -1.84E-02 | 2.52E-03 | **2.53E-13** |
| | 1_10798489_C_G | rs34071855-G | 3.42E-01 | LDL direct | -1.65E-02 | 2.50E-03 | **3.87E-11** |
| | 1_10798489_C_G | rs34071855-G | 3.42E-01 | Apolipoprotein B | -1.82E-02 | 2.60E-03 | **2.49E-12** |
| | 1_10798489_C_G | rs34071855-G | 3.42E-01 | HDL cholesterol | -5.27E-04 | 2.65E-03 | 8.42E-01 |
| | 1_10798489_C_G | rs34071855-G | 3.42E-01 | Apolipoprotein A | -3.65E-03 | 2.61E-03 | 1.63E-01 |
| | 1_10798489_C_G | rs34071855-G | 3.42E-01 | Triglycerides | -1.38E-02 | 2.59E-03 | 1.08E-07 |
| **ZNF101** | 19_19789528_A_G | rs2304130-G | 8.91E-02 | Cholesterol | -8.91E-02 | 4.14E-03 | **1.21E-102** |
| | 19_19789528_A_G | rs2304130-G | 8.91E-02 | LDL direct | -7.70E-02 | 4.12E-03 | **6.27E-78** |
| | 19_19789528_A_G | rs2304130-G | 8.91E-02 | Apolipoprotein B | -6.96E-02 | 4.28E-03 | **1.85E-59** |
| | 19_19789528_A_G | rs2304130-G | 8.91E-02 | HDL cholesterol | -1.78E-02 | 4.33E-03 | 4.10E-05 |
| | 19_19789528_A_G | rs2304130-G | 8.91E-02 | Apolipoprotein A | -8.65E-03 | 4.29E-03 | 4.36E-02 |
| | 19_19789528_A_G | rs2304130-G | 8.91E-02 | Triglycerides | -7.94E-02 | 4.26E-03 | **1.60E-77** |
